# Chromosome scale assembly of allopolyploid genome of the diatom *Fistulifera solaris*

**DOI:** 10.1101/2021.11.10.468027

**Authors:** Yoshiaki Maeda, Kahori Watanabe, Ryosuke Kobayashi, Tomoko Yoshino, Chris Bowler, Mitsufumi Matsumoto, Tsuyoshi Tanaka

## Abstract

Microalgae including diatoms are of interest for environmentally-friendly manufacturing such as biofuel production. However, only a very few of their genomes have been elucidated owing to their diversified and complex evolutionary history. The genome of the marine oleaginous diatom *Fistulifera solaris*, an allopolyploid diatom possessing two subgenomes, has been analyzed previously by pyrosequencing. However, many unsolved regions and unconnected scaffolds remained. Here we report the entire chromosomal structure of the genome of *F. solaris* strain JPCC DA0580 using a long-read nanopore sequencing platform. From just one single run using a MinION flow-cell, the chromosome scale assembly with telomere-to-telomere resolution was achieved for 41 out of 44 chromosomes. Centromere regions were also predicted from the chromosomes, and we discovered conserved motifs in the predicted regions. The function of the motifs was experimentally confirmed by successful transformation of the diatom via bacterial conjugation. This discovery provides insights into chromosome replication, facilitating the rational design of artificial chromosomes for large-scale metabolic engineering of diatoms. The chromosome scale assembly also suggests the potential existence of multi-copy mini-chromosomes and tandemly repeated lipogenesis genes related to the oleaginous phenotype of *F. solaris*. The nanopore sequencing also solved the sequential arrangement of the repeat region in the *F. solaris* mitochondrial genome. Findings of this study will be useful to understand and further engineer the oleaginous phenotype of *F. solaris*.

## Introduction

Serious damage caused by global warming has already been documented ^1, 2^. Facing this issue and finding solutions is the overriding priority for humankind to sustain the planet. Every possible measure needs to be taken to decrease the emission of green-house gases including CO_2_. Among them, development of renewable fuels instead of conventional fossil fuels is one of the hopeful options to be considered. Research into renewable fuels has passed from a phase of searching for fuel sources from edible biomass (e.g., bioethanol derived from starch and biodiesel derived from edible plant oil) to non-edible cellulose-based biomass. Microalgal biomass has additionally been recognized as a promising feedstock due to no-competition with food production, higher biomass productivity than cellulose-based biomass ^3^, as well as the potential of microalgae-derived oil to be used as an alternative to jet-fuels ^4, 5, 6^, which cannot be mixed with bioethanol and biodiesel produced in previous generations of biofuel research.

Some microalgae accumulate high levels of oil in their cells, and can serve as production hosts of microalgal oil to be converted to renewable fuels. Such oleaginous microalgae including *Nannochrolopsis gagitana* ^7^, *Chlorella vulgaris* ^8^, and *Phaeodactylum tricornutum* ^9^ have been investigated all over the world by means of multi-omics analyses, to elucidate the molecular basis supporting their oil accumulation phenotypes. In particular, genome analyses have provided genetic insights into lipid metabolism, and clues for enhancement of lipid production by metabolic engineering. Among such promising microalgae, we have focused on the oleaginous diatom, *Fistulifera solaris*, which accumulates significant amounts of lipids up to ~65 wt% of its dry cell weight ^10^. Multi-omics analyses including genomics ^11, 12^, transcriptomics ^11^, proteomics ^13, 14^ and metabolimcs^15^ have been performed for this diatom, and the biological factors involved in its excellent lipid productivity have been partially elucidated. However, we have to point out the shortcomings of our previous analyses which were based on genomic information obtained from pyrosequencing of short sequence reads ^12^.

In general, short read sequencing technology such as the pyrosequencing and Illumina technology generate relatively accurate but poorly contiguous assemblies which tend to contain unresolved repeat regions and unlinked contigs or scaffolds ^16^. In our case, the previous analysis generated an *F. solaris* genome assembly containing 117 gaps with unresolved sequences, and 53 out of 295 scaffolds unlinked to the hypothetically determined pseudo-chromosomes ^12^. In addition, only 9 out of 84 pseudo-chromosomes achieved telomere-to-telomere resolution. These facts strongly indicate that the previous assembly did not fully cover the entire sequence of the *F. solaris* genome, and might overlook uncertain genomic regions which are related to its oleaginous phenotype. Furthermore, the limits of the current genomic information might negatively impact elucidation of the complex genomic features of this diatom, in particular allopolyploidy caused by interspecies hybridization between distinct parental species ^11^. Allopolyploidy is considered to be one of the major driving forces generating plant diversity, and has been intensively studied in important agricultural plants such as *Triticum aestivum* (wheat) ^17^, *Gossypium hirsutum* (cotton) ^18^, *Brassica* (oil seed) ^19^ and *Elaeis guineensis* strain *Tenera* (African oil palm) ^20^. In contrast to multicellular plants, *F. solaris*, which has two distinct subgenomes denoted Fso_h and Fso_l 11, is the sole example of a unicellular microalgal allopolyploid reported thus far. We previously analyzed the differences in gene expression patterns of the two subgenomes, and discussed their roles in lipid metabolism ^11^, whereas we did not discover any particular differences in genomic structures between the subgenomes, potentially due to both its complex allopolyploid genomic structure and the limitation of the short-read pyrosequencing assembly.

To address this issue, in this study we re-sequenced the *F. solaris* genome using an Oxford Nanopore Technology (ONT) sequencer, MinION. Nanopore sequencing is an emerging technology allowing particularly long-read sequencing, consequently leading to high contiguity. Re-sequencing efforts using MinION have been recently reported to resolve structural features of the genomes of model organisms including a nematode *Caenorhabditis elegans* ^16^, a seed-plant *Arabidopsis thaliana* ^21^, and the diatoms *Phaeodactylum tricornutum* and *Thalassiosira pseudonana* ^22^. These studies resolved unmapped sequences and unanchored or misassembled contigs, guiding us to re-examine the complex allopolyploid genome of *F. solaris* using MinION.

Here, we extracted high molecular weight (HMW) genomic DNA from *F. solaris* and subjected it to nanopore sequencing. The generated contigs were assessed by bioinformatics analyses to resolve a chromosome scale assembly, in which sequence features of the telomeres and centromeres were detected. The consensus motifs were discovered in the *F. solaris* centromeres potentially containing autonomous replication sequences (ARSs), which have never been reported previously in diatoms. Replication capacities of the identified ARSs were assessed by constructing the episomal vectors containing each of them towards the design of artificial chromosomes. The profiling of the read depth along the chromosomes suggests the potential existence of aneuploidy. Moreover, we found tandemly repeated genes which could be directly involved in the biosynthesis of fatty acids and glycerolipids. These finding were solely attributed to the chromosome scale assembly generated by MinION.

## Results

### Nanopore sequencing and assembly of Fistulifera solaris genome

A single run of nanopore sequencing of HMW genomic DNA of *F. solaris* (Supplementary Fig. S1) generated approximately 4.1×10^5^ passed reads (~4.8 Gbp). Because our previous study using pyrosequencing suggested that the size of *F. solaris* nuclear genome was 49.7 Mbp, the coverage depth was estimated to be 97 times, which was comparable with the nanopore sequencing of *C. elegans* genome (the coverage depth was 103 times) ^16^. Distribution of read length and quality Q-score of the passed reads is shown in Supplementary Fig. S2. The average and maximal read lengths were 11.6 and 115.8 kbp, respectively (Table 1). We found 5 and 6,634 reads longer than 100 and 50 kbp, respectively. The average quality Q-score of the passed reads was 10.3, indicating a read error rate of approximately 9.3%.

**Table 1.**
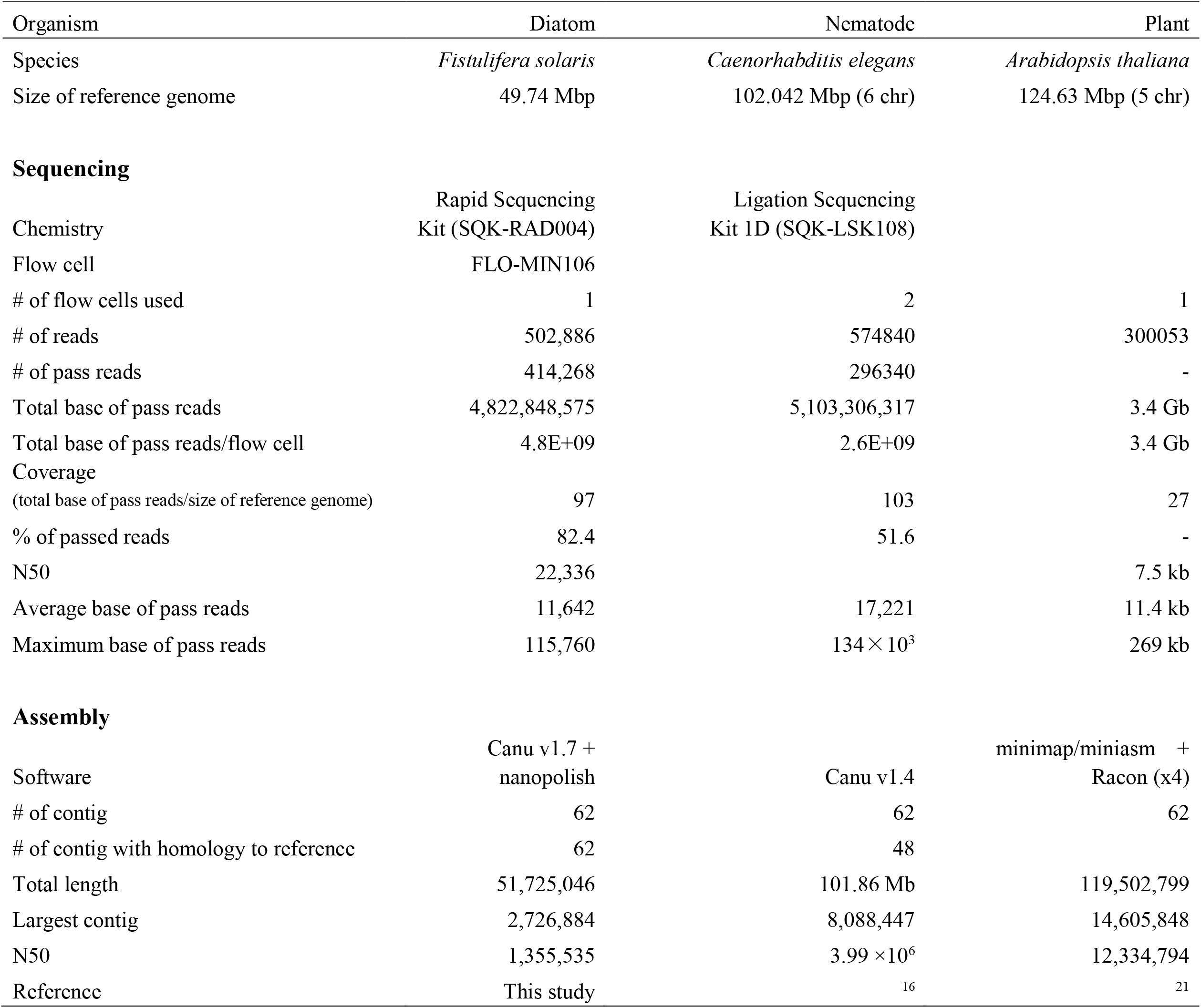
Summary of the nanopore sequencing of the genome of *Fistulifera solaris*, and comparison to other model organisms

| Organism | Diatom | Nematode | Plant |
| --- | --- | --- | --- |
| Species | *Fistulifera solaris* | *Caenorhabditis elegans* | *Arabidopsis thaliana* |
| Size of reference genome | 49.74 Mbp | 102.042 Mbp (6 chr) | 124.63 Mbp (5 chr) |
| **Sequencing** |  |  |  |
|  | Rapid Sequencing | Ligation Sequencing |  |
| Chemistry | Kit (SQK-RAD004) | Kit 1D (SQK-LSK108) |  |
| Flow cell | FLO-MIN106 |  |  |
| # of flow cells used | 1 | 2 | 1 |
| # of reads | 502,886 | 574840 | 300053 |
| # of pass reads | 414,268 | 296340 | - |
| Total base of pass reads | 4,822,848,575 | 5,103,306,317 | 3.4 Gb |
| Total base of pass reads/flow cell | 4.8E+09 | 2.6E+09 | 3.4 Gb |
| Coverage |  |  |  |
| (total base of pass reads/size of reference genome) | 97 | 103 | 27 |
| % of passed reads | 82.4 | 51.6 | - |
| N50 | 22,336 |  | 7.5 kb |
| Average base of pass reads | 11,642 | 17,221 | 11.4 kb |
| Maximum base of pass reads | 115,760 | 134 × 10<sup>3</sup> | 269 kb |
| **Assembly** |  |  |  |
|  | Canu v1.7 + |  | minimap/miniasm + |
| Software | nanopolish | Canu v1.4 | Racon (x4) |
| # of contig | 62 | 62 | 62 |
| # of contig with homology to reference | 62 | 48 |  |
| Total length | 51,725,046 | 101.86 Mb | 119,502,799 |
| Largest contig | 2,726,884 | 8,088,447 | 14,605,848 |
| N50 | 1,355,535 | 3.99 × 10<sup>6</sup> | 12,334,794 |
| Reference | This study | 16 | 21 |

Assembly of these reads was conducted by Canu ^23^, followed by sequence polishing using nanopolish ^24^. As a result, a total of 62 contigs (~51.7 Mbp, Table 1) ranging from 3.4 kbp to 2.7 Mbp with N50 of 1.4 Mbp were generated (Supplementary Fig. S3 (A)). The N50 value obtained in this study was the same in the order of magnitude as that of *C. elegans* ^16^.

Among the 62 contigs, 2 contigs (contigs 139 and 145) showed high sequence identity with the chloroplast genome, and their GC% (approximately 32%, Supplementary Fig. S3 (B)) were matched with the chloroplast genome ^25^. We found that another contig (the smallest contig 208, GC%=54%) showed relatively low sequence identity to the chloroplast genome (sequence identity: 77.3%), while it showed similarities (approximately 90%) to partial sequences of *Sphingomonas* spp. genomes, implying potential contamination. Three contigs (contigs 1, 4, and 6) showed high sequence identities to the mitochondrial genome, and their GC% (approximately 28%, Supplementary Fig. S3 (B)) were matched with that of the mitochondrial genome ^26^. Other than these 6 contigs, the remaining 56 contigs showed high sequence identity with the nuclear genome of *F. solaris*, and have a GC% (approximately 46%, Supplementary Fig. S3 (B)) consistent with the nuclear genome of *F. solaris* ^12^. Among the 56 contigs corresponding to the nuclear genome, 11 had short lengths ranging from 34 to 60 kbp (Supplementary Fig. S3 (A)), and showed high sequence identity with other longer contigs (Supplementary Table S3). It remained unclear why such redundant contigs were generated during the assembly process. However, the sequences of these short contigs were almost always comprised within the corresponding longer contigs (Supplementary Table S3), and thus we concluded that they can be ignored for further analyses. We defined that the remaining 45 contigs correspond to the nuclear genome of *F. solaris.*

In our previous studies, pyrosequencing of *F. solaris* genome generated 295 scaffolds consisting of 84 types of the hypothetical chromosome structures ^12^, among which 120 and 122 scaffolds were classified into Fso_h and Fso_l subgenomes composed of 42 hypothetical chromosomes, respectively ^11^. The remaining 53 scaffolds were could not be classified into either of the subgenomes because they were not involved in the hypothetical chromosome structures. When we aligned the 295 pyrosequencing scaffolds and the 45 MinION contigs, 115 of the 120 scaffolds belonging to the Fso_h subgenome were aligned to 23 MinION contigs, and 120 of the 121 scaffolds belonging to the Fso_l subgenome were aligned to 22 MinION contigs (Fig. 1). Therefore, we defined these sets of MinION contigs as Fso_h and Fso_l subgenomes predicted by nanopore sequencing, respectively. Twenty-three MinION contigs classified as Fso_h subgenome also included one scaffold formerly belonging to the Fso_h subgenome and 25 scaffolds formerly not classified into either subgenome. Twenty-two MinION contigs classified as Fso_l subgenome also showed sequence similarity with 5 scaffolds formerly belonging to the Fso_h subgenome and 28 scaffolds formerly not classified into either subgenome. When we compared the sequences of 23 and 22 MinION contigs classified as Fso_h and Fso_l subgenomes, these sets of MinION contigs showed one-to-one homoeologous correspondence, with an exceptional case where contig 1973 and 1980 (Fso_h) together corresponded to the homoeologous counterpart of contig 76 (Fso_l), and thus we considered that these two contigs were together parts of a single DNA molecule. The length and sequence between contig 1973 and 1980 remain to be determined. We found putative 5S rRNA genes at the termini of contig 1973 and 1980 (Supplementary Fig. S4), suggesting that repeat structures related to 5S rRNA genes, which are frequently found in eukaryotes ^27^, might prohibit the assembly process by Canu.

**Fig. 1.**
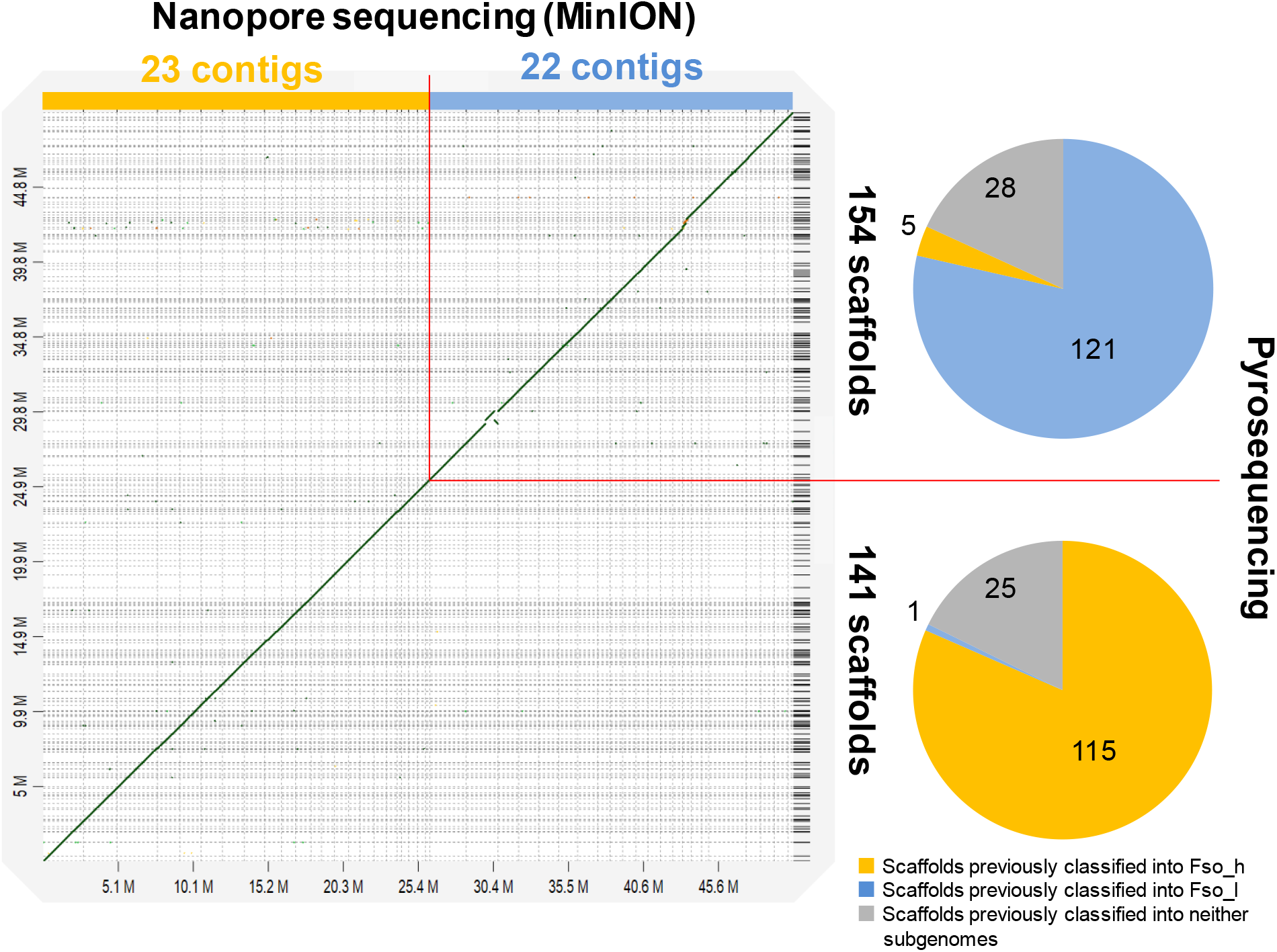
Dot plot analysis by aligning the MinION contigs obtained in this study with previously obtained pyrosequencing scaffolds. Among 25 MinION contigs, 23 and 22 contigs corresponding to the nuclear genome were classified into Fso_h and Fso_l subgenomes of the allopolyploid genome of *F. solaris*, respectively. Among 195 scaffolds previously obtained by pyrosequencing, 141 and 154 scaffolds (both of which included the scaffolds previously classified into Fso_h, Fso_l, or neither subgenomes) showed sequence similarity to the MinION contigs in Fso_h and Fso_l subgenomes, respectively.

Figure 2 shows a total of 22 pairs of these contigs as the homoeologous chromosomes predicted in this study (second outer circle), along with 84 types of the hypothetical chromosomes previously predicted (first outer circle) ^12^. Several hypothetical chromosomes predicted by pyrosequencing were included in single MinION contigs, suggesting the high contiguity of MinION contigs obtained in this study.

**Fig. 2.**
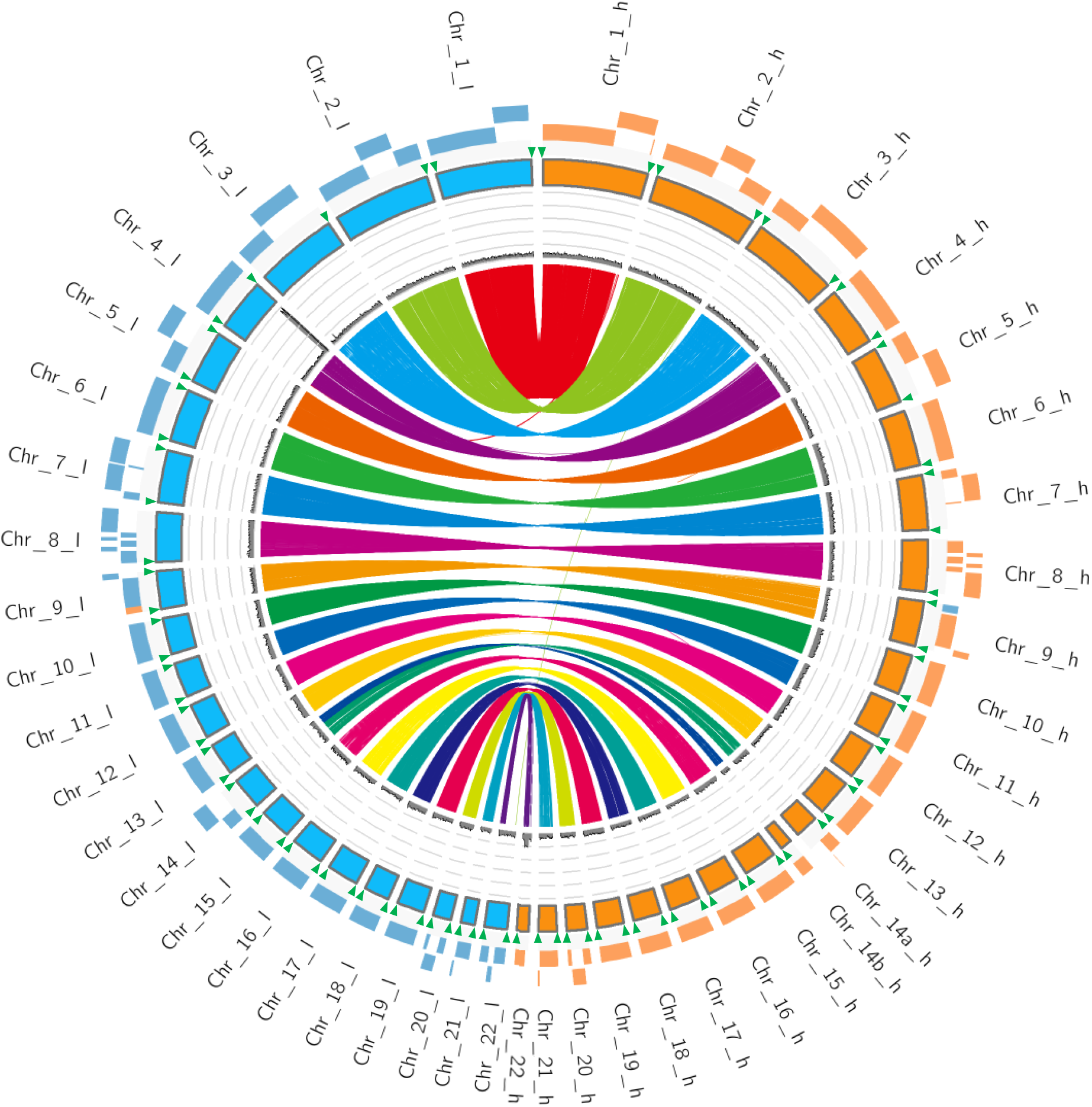
Circular view of the landscape of the allopolyploid genome of *Fistulifera solaris* as revealed by MinION sequencing. The outermost circle (orange and blue tiles without outlines) are pyrosequencing scaffolds previously obtained. The second outer most circle are MinION contigs obtained in this study (orange and blue tiles with outlines). Telomeric repeats found in the MinION contigs are shown by green triangles. The chromosome numbers in two subgenomes (Fso_h and Fso_l) are marked on these circles. The third circle represents the read depth along the MinION contigs. The lines in the innermost circle represents the positions of homoeologous gene pairs on the MinION contigs.

### Telomeres and centromeres found in MinION contigs

Telomeres are fundamental structures of chromosomes, and thus these structures can be benchmarks to assess the contiguity and completeness of the assembled contigs. Telomeric repeats (CCCTAA) ^28^ were found at 82 positions out of 88 termini (~93%) of the 44 chromosomes (green triangles on the second outer circle of Fig. 2). All chromosomes have telomeric repeats at either or both termini. In the hypothetical chromosomes previously predicted, telomeric repeats were found at only 54 of the 168 termini (~32%) ^12^. These results suggested the high contiguity and completeness of the contigs obtained in this study. We propose that these sets of MinION contigs represent the chromosome structure of the allopolyploid genome of *F. solaris* (Fig. 2).

Although centromeres are also basic landmarks of chromosomes, sequence features of diatom centromeres are largely unknown. Recently, diatom centromeres were predicted as low-GC content regions in the genome of the model diatom *P. tricornutum* ^29^. The predicted regions were supported by a chromatin immunoprecipitation experiment combining with illumina sequencing (ChIP-seq). We applied the same prediction method for the chromosomes of *F. solaris*. In the employed method, the number of 100 bp windows less than or equal to 32% GC within a 3 kbp window (N_box32_) were counted, and plotted along the contigs. In the *P. tricornutm* genome, clear peaks, which were often found once per chromosome, were observed in almost all chromosomes. However, in the proposed chromosomes of *F. solaris*, a limited number of them had clear peaks, and 8 homoeologous chromosome pairs (chromosomes 1, 3, 10, 11, 13, 14, 20, and 22) possessed the peaks at similar positions (Supplementary Fig. S5). We hypothesized that the position of these N_box32_ peaks indicated the centromeres. Homoeologous chromosome pairs were presumably derived from distinct parental species which would be phylogeneticaly closely related, and thus we assumed that the positional relationships of the hypothetical centromeres and neighboring genes should resemble between homoeologous chromosome pairs. To assess this assumption, we confirmed the positional relationships, and found that the proposed centromeres on each homoeologous chromosome pair and the neighboring genes were exactly conserved in 7 of 8 chromosome pairs, except for chromosome 13. Although a positional shift of the centromeres was found in chromosome 13, a similar array of genes surrounds the proposed centromeres. Furthermore, putative centromere regions were narrowed by removing the regions generating transcription signals based on our previous transcriptome data ^12^ because, in general, eukaryotic centromeres are non-transcribed regions ^30^ (Supplementary Fig. S6). Eventually, we tentatively determined the 16 sequences with low-GC contents (36~42%) in the 8 chromosome pairs as putative centromeres in the *F. solaris* genome (Supplementary data 1).

### Functional assessment of the centromeres with autonomous replication sequences

The centromeres of Chr 1_l, 13_l, and 22_h contain small-scale repeated elements (Supplementary Fig. S7), but none of the centromeres appear to contain highly repeated elements that are typical of other eukaryotic genomes ^31^. A dot-plot analysis to assess the sequence similarities between the putative centromeres indicates that some centromeres showed sequence-similarities to their homoeologous pairs (i.e., Chr 1, 3, 14, Supplementary Fig. S7), whereas, other than these, the large-scale conserved sequences were not found within the proposed centromeres in *F. solaris*. Nonetheless, we further investigated the centromere sequences using MEME SUiTE to find other small-scale common motifs. As a result, three consensus motifs ( WTTTATTCCTAATTTCCTAAAGTYAGAATGYAATTTTGACATTCGACTG, TCCWTCWYTWSGATKHYRMHRAMKCAWRSWAACHGAMCRGWSCARRKAAA, and AWATGHAAAHRMAMAAWGGAAAATTCAGTCGAATRTCAADA) were discovered from 11 out of 16 centromere sequences (Fig. 3). Subsequently, we explored the discovered motifs in the entire sequences of the chromosomes (Supplementary Fig. S5). Forty-three out of 44 chromosomes contained at least one motif within their sequences, except for Chr. 17_h. As overall tendencies, multiple types of motifs were frequently found within narrow regions (41/44 chromosomes). Phylogenetic analysis of the sequences of the discovered motifs demonstrated that segregation tendency of the sequences derived from each subgenome (Supplementary Fig. S8). The region covering the N_box32_ peaks tended to contain the multiply stacked motifs. However, high N_box32_ values were not an essential condition to find the motifs because the regions with low N_box32_ also contain multiple motifs (Supplementary Fig. S5). These motifs were localized at certain intervals along chromosomes. This localization feature is similar to those of the autonomous replication sequences (ARS) found in other eukaryotic genomes ^32^. These findings guided us to investigate whether the putatively identified centromeres, some of which contain the conserved motifs whereas others do not, are responsible for chromosome replication in *F. solaris*.

**Fig. 3.**
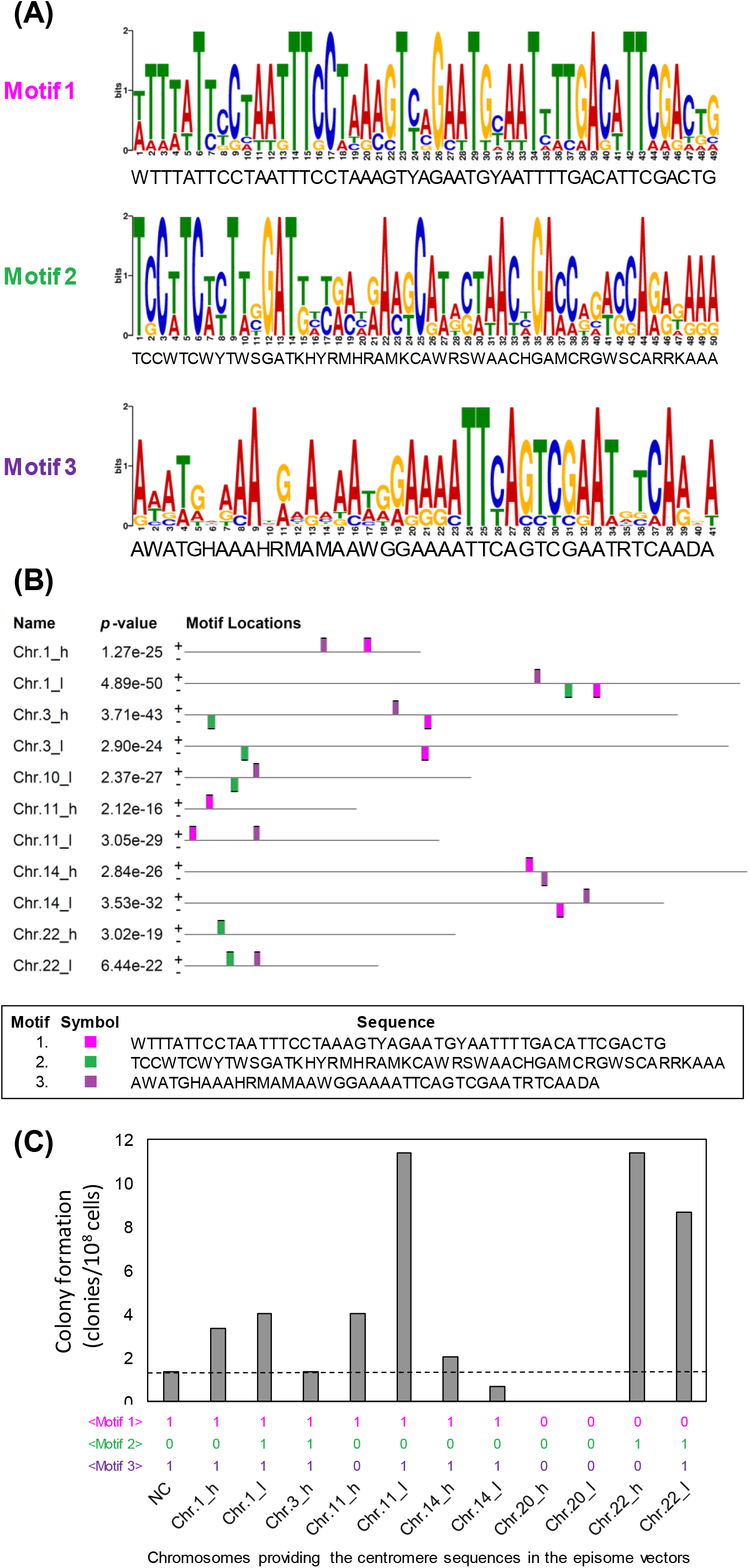
Sequence features of the centromeres putatively identified from the MinION contigs of *F. solaris*. (A) The 3 consensus motifs discovered in the centromeres potentially containing autonomous replication sequences. (B) Distributions of the consensus motifs in the centromeres. No motif was found from those of chromosome 10_h, 13_h, 13_l, 20_h, and 20_l. (C) Transformation efficiency of *F. solaris* by bacterial conjugation of the episomal vectors containing the centromeres. Negative control (NC) represents the data for the vector not containing the centromere sequences

We constructed 11 circular vectors containing a drug-resistant gene and one of the centromeres found from Chr 1_h, 1_l, 3_h, 11_h, 11_l, 14_h, 14_l, 20_h, 20_l, 22_h, and 22_l, and introduced them into *F. solaris* by bacterial conjugation ^33^. A negative control vector pFs_H4_nptII_NC was also introduced to estimate background colony formation. As a result, seven vectors generated greater numbers of drug-resistant clones than the negative control (Fig. 3 (C)). The vector with the centromeres of Chr 20_h and 20_l did not result in any transformant colonies. This preliminary data suggests that, at least, the centromeres in the above mentioned chromosomes (other than Chr 20_h and 20_l) could contribute to replication of the vectors in *F. solaris*. In particular, those with the centromeres of Chr 11_l (GC content: 36%, length: 1834 bp), 22_h (GC content: 37%, length: 1952 bp), and 22_l (GC content: 36%, length: 1396 bp) showed relatively high colony formation performance. These centromeres are relatively short and consist of low GC% sequences, consistent with the previous study investigating the maintenance of extrachromosomal artificial chromosomes in *P. tricornutum* ^29^. The functional defect of the predicted centromeres of Chr. 20_h and Chr. 20_l could be explained by the absence of the conserved motifs in these sequences. This result further supports our hypothesis that the discovered motifs could be involved in the ARS function. To the best of our knowledge, this is the first study to find consensus motifs of ARSs in diatom centromeres.

### Potential aneuploidy in the genome of Fistulifera solaris

Next, we assessed the ploidy distribution in *F. solaris*. We examined the coverage depth along the chromosomes by aligning the MinION reads toward the assembled contigs (third outer circle in Fig. 2). Overall, all chromosomes showed similar depths (average 84, Supplementary Fig. S9), with exception of chromosome 4, suggesting that the paired chromosomes derived from each parental species are contained in the *F. solaris* genome at a 1:1 ratio.

Exceptionally, the contig 49 corresponding to chromosome 4 in subgenome Fso_l has a significantly high sequencing depth. We plotted the depth values at each base along the chromosomal positions, and found a region (~135 kbp) with remarkably high depth at a terminal region (Supplementary Fig. S10). The average depth of this high depth region was 759, while that of other ordinal regions in the same chromosome is 80, comparable to other chromosomes (Supplementary Fig. S10 (A)). The high depth region was not found on the homoeologous chromosome 4 in subgenome Fso_h. To examine the reason for generation of the high depth region, we analyzed the sequence and mapping data at the boundary of the ordinal depth region and high depth region on the assembled contig. First we assessed the sequence at the boundary, and found a telomeric repeat-like sequence in the middle of the assembled contig (Supplementary Fig. S10 (C)). Subsequently, we focused on the mapping data of the MinION reads towards the assembled sequence, and found that the MinION reads mapped to the boundary could be classified into two types of reads; the reads with and without telomeric repeats (Supplementary Fig. S11). The reads without telomeric repeats were mapped to both ordinal and high depth regions, and a lot of sequence variations from the assembled sequence. By contrast, mapping position of the reads with telomeric repeats started from the boundary of the high depth region.

A possible explanation for these analytical results is the existence of independent mini-chromosomes with high sequence identity to the high depth region on chromosome 4 in the Fso_l subgenome (Supplementary Fig. S12 (A)). This hypothetical mini-chromosome (~135 kbp, containing 47 genes, Supplementary Table S4) showed approximately 8-fold copy number as compared to chromosome 4. The presence of an abnormal number of chromosomes suggests aneuploidy in the *F. solaris* genome. The existence of ARS motifs within this high read depth region (Supplementary Fig. S5) supports the existence of the hypothetical mini-chromosomes because ARSs could be required for their independent replication. Aneuploidy in diatom genomes was previously suggested in the genome of *Thalassiosira weissflogii* ^28^. Nonetheless, this is the first report of aneuploidy consisting of a partial region of a diatom chromosome.

### Organelle genome structures revealed by MinION

Long read sequencing revealed the unique features of *F. solaris* organelle genomes. In particular, the entire structure of the repeat regions in the mitochondrial genome was unveiled, which had not been fully elucidated by previous pyrosequencing ^12^. Among the 3 MinION contigs (contigs 1, 4, and 6, Supplementary Fig. S13 (A)) corresponding to the mitochondrial genome, two (contigs 4 and 6) were similar in size (~42.9 and ~43.0 kbp), while the other (contig 1) was almost double in size (~85 kbp), and repeated almost identical sequences. This was perhaps caused by mis-assembly. Contigs 4 and 6 showed high sequence identity (~99.2%), but distinguishable with several alignment gaps. This result might suggest the existence of two different types of mitochondrial genomes. Contig 6 is more similar to the mitochondrial genome sequence read by pyrosequencing (KT363689) than contig 4, and thus contig 6 was analyzed in more detail. MinION contig 6 was longer than the mitochondrial genome sequence read by pyrosequencing (~39.5 kbp). The extended regions account for a part of the repeat regions (Supplementary Fig. 13 (A)), locating between ORF251 and tRNA (UUG)-encoding gene fragment (Supplementary Fig. 13 (B)). It is widely known that diatom mitochondrial genomes contain characteristic repeat regions ^34, 35^. The common feature of these repetitive regions is that they are segregated from protein- and tRNA-coding regions, and are highly species-specific (Supplementary Fig. 13 (C)). The repeat region found in the mitochondrial genome of *Synedra acus* was relatively simple, containing two imperfect direct repeats (1,450 and 1316 bp) and short repeats less than 50 nucleotides ^34^. The *T. pseudonana* repeat region (~4.8 kbp) consists of an array of almost identical double-hairpin elements (75 bp) and its truncated forms (35 bp) are presumably generated by slipped stranded mispairing. The repeat region of *P. tricornutum* (~35 kbp) is much longer than that of *T. pseudonana*, and accounts for approximately half of the entire mitochondrial genome (~77 kbp). The *P. tricornutum* repeat region does not contain double-hairpin elements but consists of the complex arrays of 5 types of basic elements (Elements A~E whose size range from 33 to 404 bp). The *F. solaris* repeat region consists of eight relatively large repeating blocks ranging from 345 to 943 bp, four of which are tandemly repeated in the same direction whereas the other four are arrayed in a complementary orientation (Supplementary Fig. 13 (D)). The sequences of eight repeating blocks can be aligned with some gaps (Supplementary Fig. 13 (E)). The repeat region containing complementary blocks has not been discovered in other diatom mitochondrial genomes reported so far. Although the function of the discovered complementarily repeated blocks remains unknown, its structure being obviously different from other diatoms could suggest distinct functions. Accumulation of full length sequences of diatom mitochondrial genomes including the repeated regions will assist the elucidation of their functions in the future.

### Tandemly repeated genes related to lipid metabolism

Long read nanopore sequencing allowed us to find previously undetected genomic landscapes including tandemly repeated genes. Mapping of the previously identified *F. solaris* genes to MinION contigs revealed some tandemly repeated genes. A particularly striking feature which can be related to the oleaginous phenotype is a sequence of five tandem repeat of 1-acyl-sn-glycerol-3-phosphate lysophosphatidic acid acyltransferase genes (*LPAAT*, fso:g9516) (green arrows in Fig. 4 (A)), which encodes an enzyme catalyzing the transfer of an acyl chain to lysophosphatidic acid, and is essential for assembling glycerolipids including TAGs. The tandemly repeated *LPAAT* genes are located between the genes encoding bromodomain-containing protein (fso:g9515) and aarF domain-containing kinase (fso:g12324) in chromosome 9 belonging to subgenome Fso_h. Eight read sequences covered the entire region containing these tandem arrayed genes (Fig. 4 (B)). In addition, PCR targeting the tandemly repeated *LPAAT* genes and their neighboring regions generated the amplification patterns which were consistent with the MinION assembly (Supplementary Fig. S14). These results indicate that this *bona fide* repeat feature is not the result of misassembly.

**Fig. 4.**
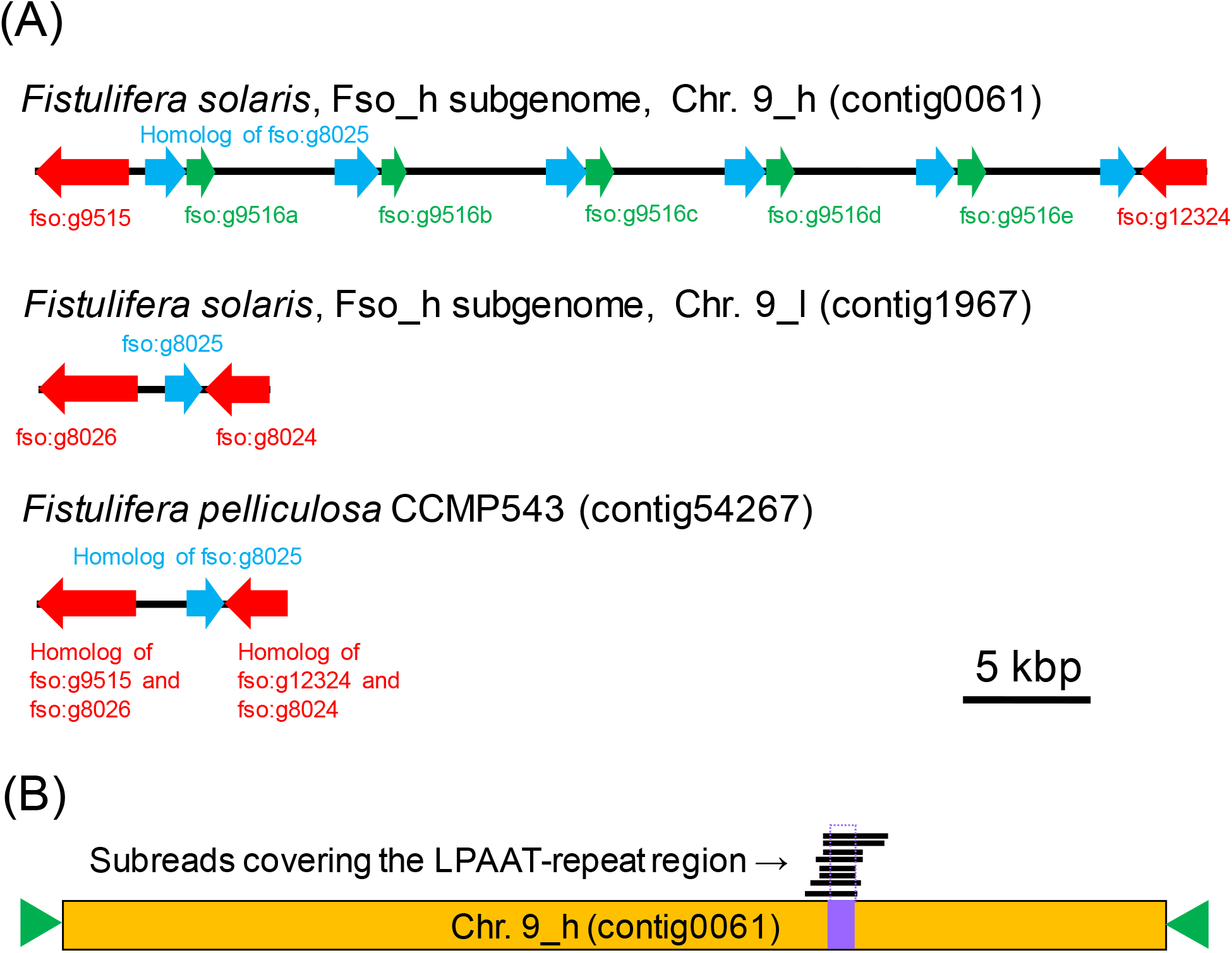
Tandemly arrayed genes of 1-acyl-sn-glycerol-3-phosphate lysophosphatidic acid acyltransferase (LPAAT) found on chromosome 9 in the Fso_h subgenome (Chr. 9_h) of *Fistulifera solaris*. (A) Arrangements of the genes of the tandemly arrayed LPAAT genes and neighboring genes in Chr. 9_h. The tandemly arrayed *LPAAT* genes were not found in the homoeologous chromosome Chr. 9_l, and the corresponding contigs of *Fistulifera pelliculosa* CCMP543, a closely-related species of *F. solaris*. (B) Eight sub-reads obtained by MinION sequencing covered the region of the tandemly arrayed LPAAT genes (shown in purple) in Chr. 9_h.

The five arrayed genes (designated as fso:g9516a~e) have two nucleotide polymorphisms (Supplementary Fig. S15 (A), fso:g9516a, b and e are identical, and other fso:g9516c and d are identical), and consequently encode two types of proteins with a single amino acid substitution (Supplementary Fig. S15 (B)). Phylogenetic analysis suggests the generation of the clade separated from other *LPAAT* genes (Supplementary Fig. S15 (C)). Repeated genes encoding transmembrane proteins with no functional annotation and homology with the protein encoded by fso:g8025 (light blue arrows in Fig. 4 (A)) were associated with these *LPAAT* repeats. Intergenic regions between the repeated *LPAAT* contain highly repeated elements (Supplementary Fig. S16) with no homology to any previously reported sequences. The functions of the repeat elements thus remain unclear.

*F. solaris* has nine genes putatively annotated as LPAAT-encoding genes. Phylogenetic analysis revealed that LPAATs encoded by the tandemly repeated fso:g9516 genes were closer to chloroplast-localized LPAATs of diatoms ^36^ and chloroplast-localized *A. thaliana* seed gene 2 (ATS2) rather than ER-localized LPAATs (Supplementary Fig. S17), whereas they do not have chloroplast-targeting signal sequences. LPAATs encoded by fso:g9516 genes possess the conserved motifs (i.e., a HX_4_D acyltransferase motif and a LPVIGW-like motif ^36, 37^, Supplementary Fig. 17), and thus they could, at least partially, retain the catalytic function even though they are shorter (96 amino acid residues) than other *F. solaris* LPAATs (ranging from 303 to 1240 residues). Furthermore, our transcriptome data obtained previously supports the expression of the tandemly repeated *LPAAT* genes during the entire period of oil accumulation, although the expression level of fso:g9516 was not the highest among the putative *LPAAT* genes (Supplementary Fig. S18). This result suggests that the tandemly repeated *LPAAT* genes could potentially contribute to glycerolipid synthesis.

In addition to the tandemly repeated *LPAAT*, we discovered tandemly repeated genes of enoyl-[acyl-carrier-protein (ACP)] reductase (EAR) as an additional feature potentially related to the oleaginous phenotype. EAR (EC1.3.1.9) functions in the fatty acid elongation process by reducing trans-2-enoyl-ACP to generate acyl-ACP, in which an NADPH is consumed. EAR genes were not fully identified in *F. solaris* genome in our previous study ^12^, and thus this new discovery now completes the fatty acid generation pathway in this organism.

## Discussion

Nanopore sequencing of the *F. solaris* genome using a MinION generated the assembled contigs the high contiguity and completeness. This chromosome-scale assembly allowed us to perform the centromere prediction based on the assumption that a single chromosome of diatoms has one centromere ^29^. We putatively identified 16 centromere sequences, within which the conserved motifs were discovered. The bacterial conjugation experiment indicates that the conserved motifs could serve as ARS in the diatom genome. The conserved motifs in the diatom centromeres have been explored using model diatom species^29^, whereas no successful study have been reported. Therefore, this is the first study to propose the functionally active centromere motifs in diatom genomes. Chromosomes belonging in the subgenomes Fso_h and Fso_l could possess ARSs with different sequence features (Supplementary Fig. S8), while those from both subgenomes can work in the same cells. Insights into functional centromeres in diatoms will be useful not only for fundamental biology studies dealing with diatom evolution, but also for biotechnological applications, in particular for the rational design of artificial chromosomes for efficient genetic engineering of diatoms.

Furthermore, read depth profiling along the chromosome suggests the feature of aneuploidy of *F. solaris* genome. Ratios of homoeologous chromosomes are not necessarily 1:1 in allopolyploid organisms due to complicated hybrid events. For example, Oku et al. revealed that larger brewing yeast *Saccharomyces pastorianus* strains (allopolyploid hybrid between *S. cerevisiae* and *S. eubayanus*) possess one set of *S. cerevisiae*-type chromosomes (haploid) and two or three sets of *S. eubayanus*-type chromosomes (diploid or triploid) ^28^. Uneven sequence depths in the middle of chromosomes were also found in *S. pastorianus*, whereas the underling mechanism was not discussed ^38^. An unusual depth was found in *F. solaris* genome not at the middle region but at the terminal region of Chr. 4_l, suggesting the existence of ~8 copy mini-chromosomes. A total of 47 genes consists of the putative mini-chromosomes (Supplementary Table S4). These genes encode some characteristic proteins which might be related to robust characteristics of *F. solaris*. For example, chloride channel 7 containing voltage activated chloride channel CLC7 type domain (GAX11033.1) is a voltage-dependent chloride channel that fine-tunes photosynthesis in plants 39. A hypothetical protein FisN_2Lh576T contains a tratricopeptide-like helical domain, which were found in the proteins involved in biogenesis of the photosynthetic apparatus ^40^. Besides these photosynthesis-related proteins, several ribosome- and translation-related proteins such as large subunit ribosomal protein L3 (GAX22749.1), 21S rRNA (GM2251-2’-O)-methyltransferase (GAX11026.1), rRNA adenine demethylase (GAX11006.1 and GAX11068.1) could be noted. Multiplication of these gene caused by anueploidization might contribute to reinforcement of photosynthesis and protein production leading to robust growth of this diatom. Further investigations will be performed to obtain more evidence to prove this aneuploidy in future studies.

In addition, tandemly repeated genes of LPAAT involved in glycerolipid synthesis and EAR involved in fatty acid synthesis were discovered from Fso_h subgenome and both subgenomes, respectively. As *F. solaris* is a allopolyploid organism, we searched a homeologous *LPAAT* gene of fso:g9516. However, no homoeologous gene was found in the Fso_l subgenome (Fig. 4 (A)). We found the homoeologous genes (fso:g8026 and fso:g8024 in Fso_l) of the neighboring genes (fso:g9515 and fso:g1232 in Fso_h), whereas only fso:g8025 exists between fso:g8026 and fso:g8024. These data suggested two possibilities; (1) the Fso_l subgenome has lost the homoeologous gene after the hybridization event through diploidization, and (2) the ancestor species providing Fso_h subgenome obtained these tandemly repeated genes prior to the hybridization event. To distinguish between these possibilities, we sequenced a closely related diatom *F. pelliculosa* CCMP543 using MinION (Supplementary Table S5), and found that the contigs contain the homologs of the neighboring genes (i.e., fso g9515 and fso:g12324). However, a homolog of fso:g9516 was not found between the homologues of neighboring genes (Fig. 4 (A) and Supplementary Fig. S19), suggesting that the tandemly repeated *LPAAT* genes might be a specific feature of the Fso_h subgenome derived from the ancestor species. By costrast, tandemly repeated EARs were found in both Fso_h (chromosome 2_h, contig 1952) and Fso_l (chromosome 2_l, contig 0014) subgenomes, as well as in the genome of the closely related pennate diatom *F. pelliculosa* (Supplementary Fig. S20 (A)). By contrast, the *EAR* gene of the model pennate diatom *P. tricornutum* is not tandemly repeated, although the arrangement of the neighboring genes is well conserved (Supplementary Fig. S20 (A)). These data suggest that tandemly repeated *EAR* genes are not common in pennate diatoms, but might be conserved in the genus *Fistulifera.* Alignment of a total of 4 EAR proteins from *F. solaris,* among which EAR1 (fso:g16257) and EAR3, and EAR 2 (fso:g16258) and EAR4 are homoeologous pairs, revealed identical amino acid sequences except for the extended C-terminal regions (Supplementary Fig. S20 (B)). DNA sequence identities between EAR1 and EAR3, and EAR2 and EAR4 are 95% and 94%, respectively (Supplementary Fig. S20 (C)), suggesting that the *EAR* homoeologous pairs contain relatively few sequence divergences as compared to other homoeologous pairs (median of identities of global homoeologous pairs is 91%). In general, allopolyploid organisms take advantage of the functional plasticity of their genomes which express similar but not identical proteins from homoeologous genes derived from distinct parental species. As opposed to this general notion, *F. solaris* might conserve the *EAR* genes to enhance fatty acid synthesis metabolism which supports the abundant production of lipids in *F. solaris.*

In conclusion, a single run of nanopore sequencer MinION revealed the previously unsolved chromosome structures of *F. solaris* genome, including centromeres, ARS, aneuploidy, and tandem repeats of lipogenesis genes. These new findings together improve our understanding of the molecular mechanisms underlying superior oil accumulation in *F. solaris*.

## Methods

### Strains and culture conditions

The marine diatom *F. solaris* JPCC DA0580 was cultured in half strength Guillard’s f medium ^41^. Cultures were grown for 5 days at 25 °C under 130 μmol photons/m^2^/sec (400 ~ 700 nm, luminometer HD2302.01 with a probe LP471PAR, Delta OHM S.r.l, Caselle di Selvazzano, Italy) of continuous illumination with 0.8 l/l/min airflow containing 2% CO_2_. Prior to and during cultivation, Hoechst 33342 (Thermo Fisher Scientific, CA, USA) was used to confirm that there was no contamination of cultures by bacteria using fluorescence microscopy.

### Genomic DNA extraction

Cells were centrifuged at 8,500 *g* for 10 min to obtain wet microalgal cells (approximately 490 mg). Hexadecyltrimethylammonium bromide (CTAB) method was used for DNA extraction. The wet microalgal cells frozen in liquid nitrogen were ruptured with a mortar and pestle, and suspended in a mixture of 9.5 ml of 10 mM Tris-HCl (pH 8.0), 0.5 ml of 10% sodium dodecyl sulfate (SDS), and 50 μl of 20 mg/ml proteinase K. After incubation for 1 hour at 37 °C, 100 μl of 10 mg/ml RNase A (RNase Cocktail Enzyme Mix, 120 U/mg, Applied Biosystems, Thermo Fisher Scientific, CA, USA) was added to the suspension, and incubated for 30 min at 37 °C. 1.8 ml of 5 M NaCl and 1.5 ml of a mixture of 10% CTAB/0.7 M NaCl were added to the suspension, and the mixture was incubated for 20 min at 65 °C. Subsequently, an equal volume phenol/chloroform/isoamyl alcohol (25:24:1) was added, and the suspension was centrifuged at 13,000 *g* for 15 min. After that, the aqueous layer was transferred to a new tube, and an equal volume of chloroform (100%) was added. The suspension was vortexed, and centrifuged at 13,000 *g* for 10 min. A tenth amount of 3 M sodium acetate and a triple amount of 100% ethanol (room temperature) were added to the aqueous layer, and the mixture was incubated at room temperature for 20 min. After centrifugation at 13,000 *g* for 30 min, the supernatant was removed, and 15 ml of 70% ethanol (room temperature) was added to the pellet. The pellet was washed and centrifuged at 13,000 *g* for 5 min. The pellet was dissolved in 200 μl of nuclease free water (New England BioLabs, Massachusetts, USA) over 2 days. The extracted genomic DNA was subjected to electrophoresis using a 1% agarose gel. Prior to library preparation, genomic DNA was purified with Ampure XP beads (Beckman Coulter).

### Nanopore sequencing

The DNA libraries for nanopore sequencing were prepared following the manufacture’s guidance for the rapid sequencing kit (SQK-RAD004) which attached adapter sequences with transposase activity (Oxford Nanopore Technologies (ONT), Oxford, UK) for *F. solaris*, and the genomic sequencing kit SQK-LSK308 for *F. pelliculosa*. The sequence library was prepared using

Sequencing was performed with nanopore sequencer MinION and MinION R9.4 flow cell (FLO-MIN106, ONT, Oxford, UK) for 48 h under the control of MinKNOW software. Resulting FAST5 files were base-called using the Albacore with parameters for FLO-MIN106. The qualities of the passed reads were analyzed and visualized using NanoStat and NanoPlot ^42^. The passed reads (Q score ≧ 7) were assembled using Canu (9). The assembled reads were polished using Nanopolish (10).

### Analysis of the assembled contigs

Lengths and GC contents of the assembled contigs were measured using fx2tab function of seqkit ^43^. Minimap2 ^44^ was used to align the MinION read sequences to the gene sequences derived from Illumina sequences obtained in our previous transcriptome analyses ^12^ to the contigs. The alignment results were visualized using integrative genomics viewer (IGV) ^45^. The read depth of each contig was analyzed using SAMtools ^46^. The contigs were classified into subgenomes and organelle genomes by comparison to those analyzed in our previous studies ^11, 12^ using D-GENIES ^47^. The circular illustration highlighting the landscapes of nuclear and organelle genomes was prepared using ClicO FS 48 and GeSeq 49, respectively. GC contents within 100-bp windows were obtained using sliding and fx2tab functions of seqkit ^43^. Sequence similarities between the predicted centromeres or within the region containing the tandemly repeated genes were assessed using BLASTn, Clustal Omega ^50^ and Dotlet ^51^. Consensus motifs in the centromeres were searched using MEME SUITE ^52^ with default setting, where the motifs with E-values smaller than 10^−10^ were employed in this study. The discovered motifs from the centromeres were searched from the MinION contigs using the motif database scanning algorithms FIMO equipped in MEME SUITE, and the motifs with p-values smaller than 10^−10^ were plotted along the contigs. Phylogenetic analyses were performed using MEGA X with the maximum likelihood method ^53^.

### Bacterial conjugation with vectors containing the predicted centromere sequences

pPtPuc3 ^33^ was obtained from Addgene (#62863). pPtPuc3_NC was constructed by removing the CEN/ARS/HIS sequence from pPtPuc3 using the primer set pPt/pFs_NC_F and pPt/pFs_NC_R (Supplementary Table S1) with PrimeSTAR Mutagenesis Basal Kit. The fragmented pPtPuc3-NC without the *sh*Ble-expression cassette and the *npt*II-expression cassette containing histone H4 gene promoter derived from *F. solaris* and *fcpA* terminator derived from *P. tricornutum* were amplified by PCR was amplified by PCR using PrimeSTAR Max DNA polymerase (TaKaRa Bio Inc., Shiga, Japan) and the primer (pFs_nptII_1_F, pFs_nptII_1_R, pFs_nptII_2_F, pFs_nptII_2_R, pFs_nptII_3_F, pFs_nptII_3_R, pFs_nptII_4_F, pFs_nptII_4_R, pFs_nptII_5_F, pFs_nptII_5_R). The amplified fragments were separated by agarose-gel electrophoresis and purified using QIAquick Gel Extraction Kit (QIAGEN, Limburg, Neferland). DNA concentrations were measured using a droplet-based absorptiometer, e-Spect (Malcom Co. Ltd, Tokyo, Japan). The purified fragments were assembled using NEBuilder HiFi DNA Assembly Master Mix (New England Biolabs Japan, Tokyo) at 50 °C (3 h). The assembled samples were introduced into *E. coli* strain TOP10 with a heat shock method (42 °C, 45 sec). The colonies were cultured in LB medium, and the resulting plasmids were extracted and subjected to Sanger sequencing to confirm the assembly. The resulting plasmid was designated as pFs_H4_nptII_NC (a negative control vector lacking centromere sequences). The linearized pFs_H4_nptII_NC was amplified by PCR using primer set, pFS_H4_nptII_golden_F and pFS_H4_nptII_golden_R. Subsequently, The DNA fragments of the putatively identified 11 centromeres were separately amplified by PCR using the *F. solaris* genomic DNA as a template and the primers listed in Supplementary Table S1, followed by separation by agarose-gel electrophoresis and purification. The linearized pFs_H4_nptII_NC (100 ng), each centromere fragment (equimolar to pPtPuc3), *Bsa*I (10 U, New England Biolabs Japan, Tokyo) and T4 DNA ligase (200 U, New England Biolabs Japan, Tokyo) were mixed, and subjected to the thermal cycles (50 cycles of 37 °C (5 min) for digestion and 16 °C (5 min) for ligation, followed by 55 °C (15 min) and 85 °C (20 min)). Each assembled sample was introduced into *E. coli* strain TOP10 with a heat shock method (42 °C, 45 sec). The colonies were cultivated in LB medium, and the resulting plasmids were extracted and subjected to Sanger sequencing to confirm the assemblies. Descriptions of the 12 vectors introduced into the diatom cells are summarized in Supplementary Table S2.

Bacterial conjugation experiments were performed by the method previously described ^33^ with some modifications. *E. coli* strain S17-1 harboring the constructed vectors was cultivated in the 5 ml of LB medium containing 50 μg/ml kanamycin at 37 °C until the optical density at 600 nm reached 0.8~1.0. The cultivated cells were harvested by centrifugation. After the supernatant was discarded, the cells were suspended in 250 μl of SOC medium, followed by cell counting. The microalgal cells were cultured in f/2 medium, and harvested by centrifugation. The harvested microalgal cells (5.0×10^7^ cells) were spread on f/2 medium plates prepared with 1wt% agar and 5vol% LB medium). The cell suspension of *E. coli* harboring each episomal vector (8.4×10^9^ cells) was overlaid on the microalgal cells, followed by air drying. The as-prepared plates were incubated at 30 °C for 90 min in the dark, and subsequently at 25 °C for 48 h under light conditions (130 μmol photons/m^2^/sec). Afterwards, the cells were retrieved from the plates, and cultivated on f/2 medium agar plates containing 500 μg/ml G-418 at 25 °C under continuous light conditions (130 μmol photons/m^2^/sec) for 4~5 weeks.

### PCR amplification of lipogenesis-related genes with a tandemly repeated structure

The presence of a lipogenesis-related gene (gene ID: fso:g9516, having a tandemly repeated structure) was confirmed using PCR. Primers were prepared based on the sequence of a fso:g9516 and synthesized by Invitrogen (Massachusetts, USA) (Table 1). As the DNA polymerase for PCR, PrimeSTAR GXL DNA Polymerase (Takara Bio Inc., Shiga, Japan) was used.

### Data access

Data of nanopore sequencing of *F. solaris* and *F. pelliculosa* can be found in the DDBJ Sequence Read Archive. Accession numbers of the Bioproject for *F. solaris* and *F. pelliculosa* are PRJDB11829 and PRJDB12112, respectively.

## Acknowledgements

This study was supported by JSPS KAKENHI Grant-in-Aid for Scientific Research C [grant number 18K04847 and 21K04784] (granted to Y. M.), and a project commissioned by the New Energy and Industrial Technology Development Organization (NEDO) [grant number 20350498] (granted to T. T.). The Global Innovation Research Organization (GIR) at TUAT also supports this international joint research.

## Refereances

1. Maddison D, Rehdanz K. The impact of climate on life satisfaction. Ecol Econ 70, 2437–2445 (2011).

2. Lemoine D, Kapnick S. A top-down approach to projecting market impacts of climate change. Nat Clim Chang 6, 51–55 (2016).

3. Maeda Y, Yoshino T, Matsunaga T, Matsumoto M, Tanaka T. Marine microalgae for production of biofuels and chemicals. Cur Opin Biotechnol 50, 111–120 (2018).

4. Fortier M-OP, Roberts GW, Stagg-Williams SM, Sturm BS. Life cycle assessment of bio-jet fuel from hydrothermal liquefaction of microalgae. Appl Ener 122, 73–82 (2014).

5. Wei H, Liu W, Chen X, Yang Q, Li J, Chen H. Renewable bio-jet fuel production for aviation: A review. Fuel 254, 115599 (2019).

6. Bwapwa JK, Anandraj A, Trois C. Possibilities for conversion of microalgae oil into aviation fuel: a review. Renew Sustain Energy Rev 80, 1345–1354 (2017).

7. Radakovits R, et al. Draft genome sequence and genetic transformation of the oleaginous alga *Nannochloropis gaditana*. Nat Commun 3, 686 (2012).

8. Cecchin M, et al. *Chlorella vulgaris* genome assembly and annotation reveals the molecular basis for metabolic acclimation to high light conditions. Plant J 100, 1289–1305 (2019).

9. Ge F, et al. Methylcrotonyl-CoA carboxylase regulates triacylglycerol accumulation in the model diatom *Phaeodactylum tricornutum*. Plant Cell 26, 1681–1697 (2014).

10. Maeda Y, Nojima D, Yoshino T, Tanaka T. Structure and properties of oil bodies in diatoms. Philos Trans R Soc Lond B Biol Sci 372, 20160408 (2017).

11. Nomaguchi T, et al. Homoeolog expression bias in allopolyploid oleaginous marine diatom *Fistulifera solaris*. BMC Genomics 19, 330 (2018).

12. Tanaka T, et al. Oil accumulation by the oleaginous diatom *Fistulifera solaris* as revealed by the genome and transcriptome. Plant Cell 27, 162–176 (2015).

13. Nojima D, Yoshino T, Maeda Y, Tanaka M, Nemoto M, Tanaka T. Proteomics analysis of oil body-associated proteins in the oleaginous diatom. J Proteome Res 12, 5293–5301 (2013).

14. Nonoyama T, et al. Proteomics analysis of lipid droplets indicates involvement of membrane trafficking proteins in lipid droplet breakdown in the oleaginous diatom *Fistulifera solaris*. Algal Res 44, 101660 (2019).

15. Liang Y, Osada K, Sunaga Y, Yoshino T, Bowler C, Tanaka T. Dynamic oil body generation in the marine oleaginous diatom *Fistulifera solaris* in response to nutrient limitation as revealed by morphological and lipidomic analysis. Algal Res 12, 359–367 (2015).

16. Tyson JR, O’Neil NJ, Jain M, Olsen HE, Hieter P, Snutch TP. MinION-based long-read sequencing and assembly extends the *Caenorhabditis elegans* reference genome. Genome Res 28, 266–274 (2018).

17. (IWGSC) IWGSC. A chromosome-based draft sequence of the hexaploid bread wheat (*Triticum aestivum*) genome. Science 345, 1251788 (2014).

18. Zhang T, et al. Sequencing of allotetraploid cotton (*Gossypium hirsutum* L. acc. TM-1) provides a resource for fiber improvement. Nat Biotechnol 33, 531–537 (2015).

19. Wang X, et al. The genome of the mesopolyploid crop species *Brassica rapa*. Nat Genet 43, 1035–1039 (2011).

20. Jin J, et al. Transcriptome and functional analysis reveals hybrid vigor for oil biosynthesis in oil palm. Sci Rep 7, 439 (2017).

21. Michael TP, et al. High contiguity *Arabidopsis thaliana* genome assembly with a single nanopore flow cell. Nat Commun 9, 541 (2018).

22. Filloramo GV, Curtis BA, Blanche E, Archibald JM. Re-examination of two diatom reference genomes using long-read sequencing. BMC Genomics 22, 379 (2021).

23. Koren S, Walenz BP, Berlin K, Miller JR, Bergman NH, Phillippy AM. Canu: scalable and accurate long-read assembly via adaptive k-mer weighting and repeat separation. Genome Res 27, 722–736 (2017).

24. Loman NJ, Quick J, Simpson JT. A complete bacterial genome assembled de novo using only nanopore sequencing data. Nat Methods 12, 733–735 (2015).

25. Tanaka T, et al. High-throughput pyrosequencing of the chloroplast genome of a highly neutral-lipid-producing marine pennate diatom, *Fistulifera* sp. strain JPCC DA0580. Photosynthesis Res 109, 223–229 (2011).

26. Tang X, Bi G. Complete mitochondrial genome of *Fistulifera solaris* (Bacillariophycidae). Mitochondrial DNA A DNA Mapp Seq Anal 27, 4405–4406 (2016).

27. Long EO, Dawid IB. Repeated genes in eukaryotes. Annu Rev Biochem 49, 727–764 (1980).

28. Bowler C, et al. The *Phaeodactylum* genome reveals the evolutionary history of diatom genomes. Nature 456, 239–244 (2008).

29. Diner RE, et al. Diatom centromeres suggest a mechanism for nuclear DNA acquisition. Proc Natl Acad Sci U S A 114, E6015–e6024 (2017).

30. Nakamura Y, et al. A stable, autonomously replicating plasmid vector containing *Pichia pastoris* centromeric DNA. Appl Environ Microbiol 84, e02882 (2018).

31. Willard HF, Waye JS. Hierarchical order in chromosome-specific human alpha satellite DNA. Trends in Genetics 3, 192–198 (1987).

32. Liachko I, Bhaskar A, Lee C, Chung SC, Tye BK, Keich U. A comprehensive genome-wide map of autonomously replicating sequences in a naive genome. PLoS Genet 6, e1000946 (2010).

33. Karas BJ, et al. Designer diatom episomes delivered by bacterial conjugation. Nat Commun 6, 6925 (2015).

34. Ravin NV, et al. Complete sequence of the mitochondrial genome of a diatom alga *Synedra acus* and comparative analysis of diatom mitochondrial genomes. Cur Genet 56, 215–223 (2010).

35. Oudot-Le Secq MP, Green BR. Complex repeat structures and novel features in the mitochondrial genomes of the diatoms *Phaeodactylum tricornutum* and *Thalassiosira pseudonana*. Gene 476, 20–26 (2011).

36. Misra N, Panda PK, Parida BK. Genome-wide identification and evolutionary analysis of algal LPAT genes involved in TAG biosynthesis using bioinformatic approaches. Mol Biol Rep 41, 8319–8332 (2014).

37. Ghosh AK, Chauhan N, Rajakumari S, Daum G, Rajasekharan R. At4g24160, a soluble acyl-coenzyme A-dependent lysophosphatidic acid acyltransferase. Plant Physiol 151, 869–881 (2009).

38. Okuno M, Kajitani R, Ryusui R, Morimoto H, Kodama Y, Itoh T. Next-generation sequencing analysis of lager brewing yeast strains reveals the evolutionary history of interspecies hybridization. DNA research : an international journal for rapid publication of reports on genes and genomes 23, 67–80 (2016).

39. Herdean A, et al. A voltage-dependent chloride channel fine-tunes photosynthesis in plants. Nat Commun 7, 11654 (2016).

40. Bohne AV, Schwenkert S, Grimm B, Nickelsen J. Roles of tetratricopeptide repeat proteins in biogenesis of the photosynthetic apparatus. Int Rev Cell Mol Biol 324, 187–227 (2016).

41. Guillard RR, Ryther JH. Studies of marine planktonic diatoms: I. *Cyclotella nana* Hustedt, and *Detonula confervacea* (Cleve) Gran. Can J Microbiol 8, 229–239 (1962).

42. De Coster W, D’Hert S, Schultz DT, Cruts M, Van Broeckhoven C. NanoPack: visualizing and processing long-read sequencing data. Bioinformatics 34, 2666–2669 (2018).

43. Shen W, Le S, Li Y, Hu F. SeqKit: A cross-platform and ultrafast toolkit for FASTA/Q file manipulation. PloS One 11, e0163962 (2016).

44. Li H. Minimap2: pairwise alignment for nucleotide sequences. Bioinformatics 34, 3094–3100 (2018).

45. Robinson JT, et al. Integrative genomics viewer. Nat Biotechnol 29, 24–26 (2011).

46. Li H, et al. The sequence alignment/map format and SAMtools. Bioinformatics 25, 2078–2079 (2009).

47. Cabanettes F, Klopp C. D-GENIES: dot plot large genomes in an interactive, efficient and simple way. PeerJ 6, e4958 (2018).

48. Cheong WH, Tan YC, Yap SJ, Ng KP. ClicO FS: an interactive web-based service of Circos. Bioinformatics 31, 3685–3687 (2015).

49. Tillich M, et al. GeSeq - versatile and accurate annotation of organelle genomes. Nuc Acids Res 45, W6–w11 (2017).

50. Sievers F, et al. Fast, scalable generation of high-quality protein multiple sequence alignments using Clustal Omega. Mol Syst Biol 7, 539 (2011).

51. Junier T, Pagni M. Dotlet: diagonal plots in a web browser. Bioinformatics 16, 178–179 (2000).

52. Bailey TL, et al. MEME SUITE: tools for motif discovery and searching. Nuc Acids Res 37, W202–208 (2009).

53. Kumar S, Stecher G, Li M, Knyaz C, Tamura K. MEGA X: Molecular evolutionary genetics analysis across computing platforms. Mol Biol Evol 35, 1547–1549 (2018).

