## Supplementary for "Chromosome scale assembly of allopolyploid genome of the diatom *Fistulifera solaris*"

Running title: Nanopore sequencing of *Fistulifera solaris* genome

Yoshiaki Maeda[1,*], Kahori Watanabe[1], Ryosuke Kobayashi[1], Tomoko Yoshino[1], Chris Bowler[2], Mitsufumi Matsumoto[3], Tsuyoshi Tanaka[1]

[1] Division of Biotechnology and Life Science, Institute of Engineering, Tokyo University of Agriculture and Technology, 2-24-16, Koganei, Tokyo, 184-8588, Japan

[2] Institut de Biologie de l'Ecole Normale Supérieure (IBENS), Ecole Normale Supérieure, CNRS, INSERM, Université PSL, 75005 Paris, France

[3]Biotechnology Laboratory, Electric Power Development Co., Ltd, 1, Yanagisaki-machi, Wakamatsu-ku, Kitakyusyu, Fukuoka, 808-0111, Japan

* Corresponding Author: Yoshiaki Maeda

Table S1 The primers used in this study

| Primer name | Sequence (5' → 3') |
|---|---|
| **Construction of episomal vectors** | |
| pPt/pFs_NC_F | CAGCGAGCTTTTACACAGGAGTCTGGACTTGAC |
| pPt/pFs_NC_R | GTGTAAAAGCTCGCTGAAGACGAGCTAGTGTTA |
| pFs_nptII_1_F | tgttcttatgtagTTTTACACAGGAGTCTGG |
| pFs_nptII_1_R | ttgaataaatcgaCTCACCAGTCACAGAAAAG |
| pFs_nptII_2_F | gtgactggtgagTCGATTTATTCAACAAAGCCAC |
| pFs_nptII_2_R | gacttggttgagTGCCAGTGTTACAACCAATTAAC |
| pFs_nptII_3_F | tgtaacactggcaCTCAACCAAGTCATTCTG |
| pFs_nptII_3_R | tggtcggcgtcggTCTAGATCAGAAGAACTCG |
| pFs_nptII_4_F | ttctgatctagaCCGACGCCGACCAACACC |
| pFs_nptII_4_R | tcctgtgtaaaaCTACATAAGAACACCTTTGGTGGAGGG |
| pFs_nptII_5_F | gagcaggactgaCCGACGCCGACCAACACC |
| pFs_nptII_5_R | tcctgtgtaaaaCTACATAAGAACACCTTTGGTGGAGGG |
| **Amplification of the region of the tandemly repeated LPAAT genes** | |
| 9515_F | TTTTTCGCTGGAGGAGGACCCGATTG |
| 9516_R | TTAAAGGGCACGACTCTTCTTG |
| 9516_F | ATGCTTGCGGCGAATCACAGCTC |
| 12324_R | TCTTCCTTTGCAGACTTTTGTTGC |
| 8026_F | ATGCTTGCGGCGAATCACAGCTC |
| 8024_R | GTTTCTTCCTTTTGCAGACTTTCATCGT |
| 9516_R2 | AGTGTCCATCCATGAGCTGTGATTC |
| 9516_F2 | TAGCAAAAATGAAGTGCAGCGCGTC |
| 12324_R | TCTTCCTTTGCAGACTTTTGTTGC |

Table S2 The vectors introduced into *Fistulifera solaris* by bacterial conjugation in this study

| Plasmid names | ARS | Antibiotics-resistance |
|---|---|---|
| pPtPuc3_NC | - | *sh*Ble/Kanamycin$^r$ |
| pFs_H4_nptII_NC | - | *npt*II/Kanamycin$^r$ |
| pFs_H4_nptII_cen1H | Centromere in Chr. 1_h | *npt*II/Kanamycin$^r$ |
| pFs_H4_nptII_cen1L | Centromere in Chr. 1_l | *npt*II/Kanamycin$^r$ |
| pFs_H4_nptII_cen3H | Centromere in Chr. 3_h | *npt*II/Kanamycin$^r$ |
| pFs_H4_nptII_cen11H | Centromere in Chr. 11_h | *npt*II/Kanamycin$^r$ |
| pFs_H4_nptII_cen11L | Centromere in Chr. 11_l | *npt*II/Kanamycin$^r$ |
| pFs_H4_nptII_cen14H | Centromere in Chr. 14_h | *npt*II/Kanamycin$^r$ |
| pFs_H4_nptII_cen14L | Centromere in Chr. 14_l | *npt*II/Kanamycin$^r$ |
| pFs_H4_nptII_cen20H | Centromere in Chr. 20_h | *npt*II/Kanamycin$^r$ |
| pFs_H4_nptII_cen20L | Centromere in Chr. 20_l | *npt*II/Kanamycin$^r$ |
| pFs_H4_nptII_cen22H | Centromere in Chr. 22_h | *npt*II/Kanamycin$^r$ |
| pFs_H4_nptII_cen22L | Centromere in Chr. 22_l | *npt*II/Kanamycin$^r$ |

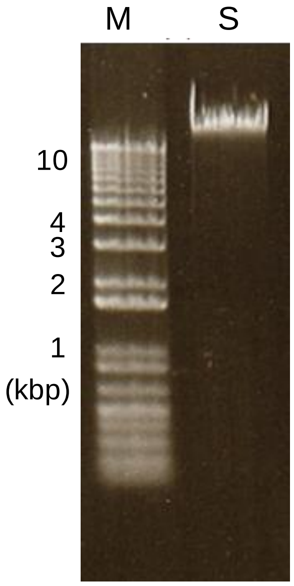

Fig. S1 Agarose gel electrophoresis of the DNA sample extracted from *Fistulifera solaris* with CTAB method, and used for nanopore sequencing. Lane M and S represent the molecular marker and genomic DNA sample.

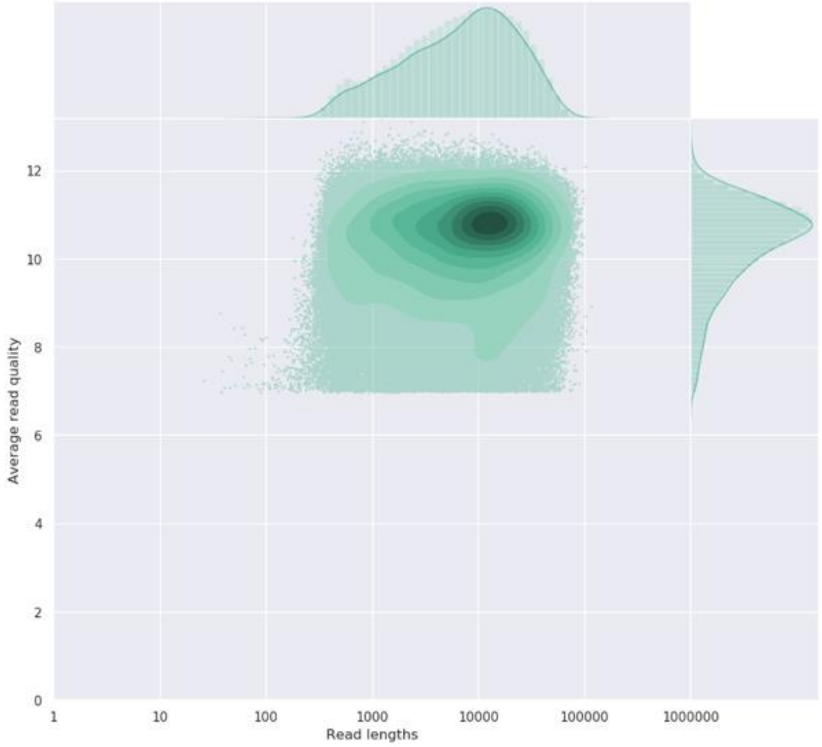

Fig. S2 Distribution of the read lengths and quality Q-scores of the passed reads obtained by nanopore sequencing of *Fistulifer solaris* genome.

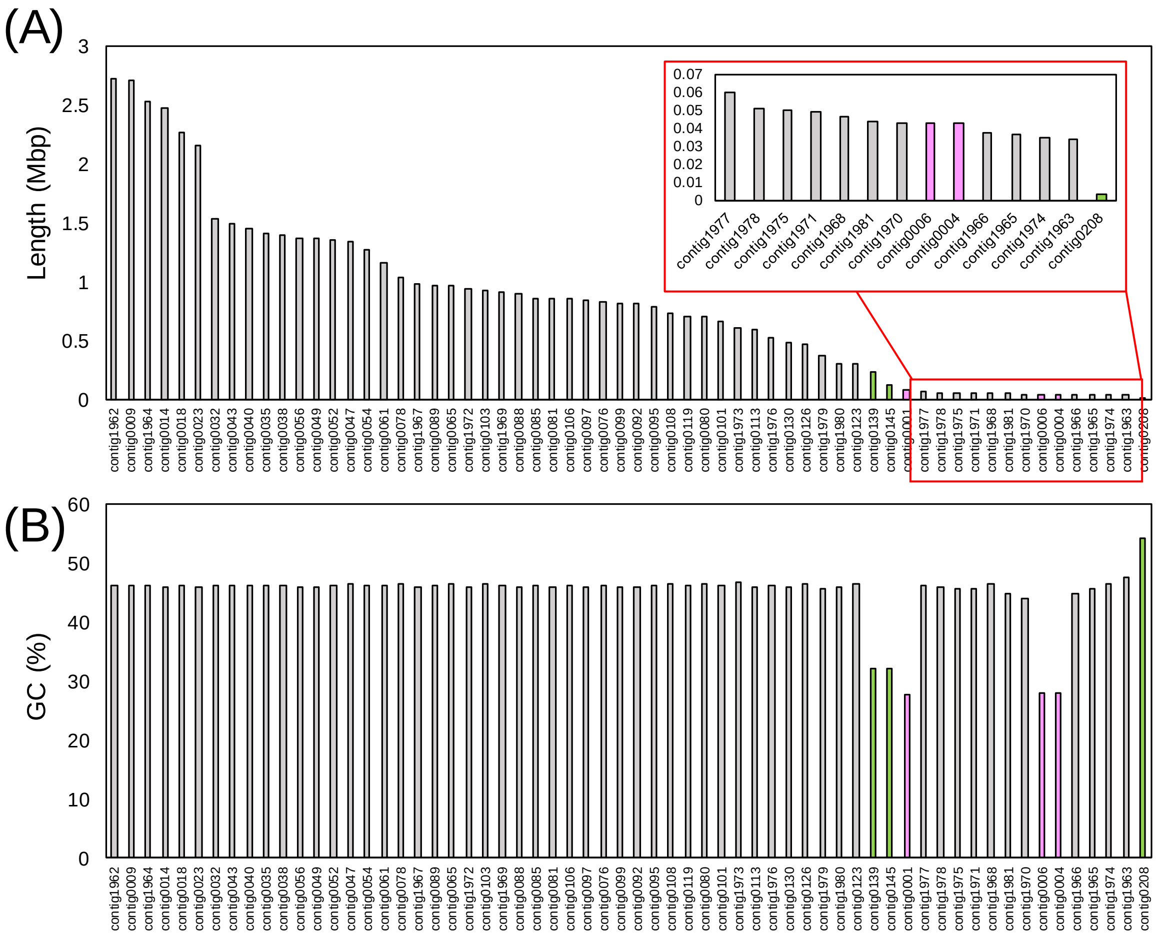

Fig. S3 Length (A) and GC% (B) of the MinION contigs of the *Fistulifera solaris* genome. Assembly of the reads were implemented by Canu. The contigs corresponding to mitochondrial and chloroplast genomes were highlighted in green and pink, respectively.

Table S3 Summary of short MinION contigs of *Fistulifera solaris* genome

| Contigs | Length (bp) | GC% | Reagion showin sequence similarity | Sequence identity |
|---|---|---|---|---|
| contig1977 | 60337 | 46.1 | 3' terminal region of contig1976 | 99.30% |
| contig1978 | 51503 | 46.0 | 3' terminal region of contig1979 | 98.42% |
| contig1975 | 50458 | 45.6 | 5' terminal region of contig1976 | 99.07% |
| contig1971 | 49208 | 45.7 | 5' terminal region of contig1972 | 98.86% |
| contig1968 | 46932 | 46.6 | 5' terminal region of contig1969 | 98.74% |
| contig1981 | 44189 | 44.9 | 3' terminal region of contig1980 | 99.35% |
| contig1970 | 43202 | 44.0 | 3' terminal region of contig1969 | 98.42% |
| contig1966 | 37290 | 45.0 | 5' terminal region of contig1967 | 98.77% |
| contig1965 | 36987 | 45.6 | 3' terminal region of contig1964 | 98.44% |
| contig1974 | 34886 | 46.6 | 3' terminal region of contig1973 | 99.00% |
| contig1963 | 34274 | 47.6 | 3' terminal region of contig1962 | 98.26% |

(A)

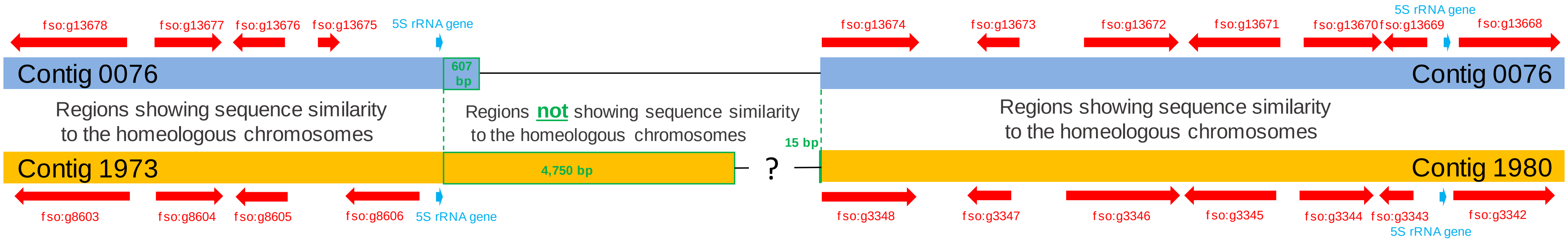

(B)

AGGAACGACCATACTACGATGACAGCACCGCTTCCCGTCTGCTCAGCGAAGTTAAGCATCGTCAGGCCCGGTTAGTACTACGGTGGGGACCACGTTGGAATCCC GGGTGTTGTTCTT

(95% sequence identity with *Diatoma tenue* gene for 5S ribosomal RNA (NCBI accession number: D00058.1))

Fig. S4 Schematic diagram of the alignment of chromosome 14_I (contig 76) and chromosome 14_h (contig 1973 and 1980) (A). The sequence between contig 1973 and 1980 (shown as question mark) remain to be determined. 5S rRNA genes (B) (light blue arrows in (A)) were found close to the gap between contig 1973 and 1980.

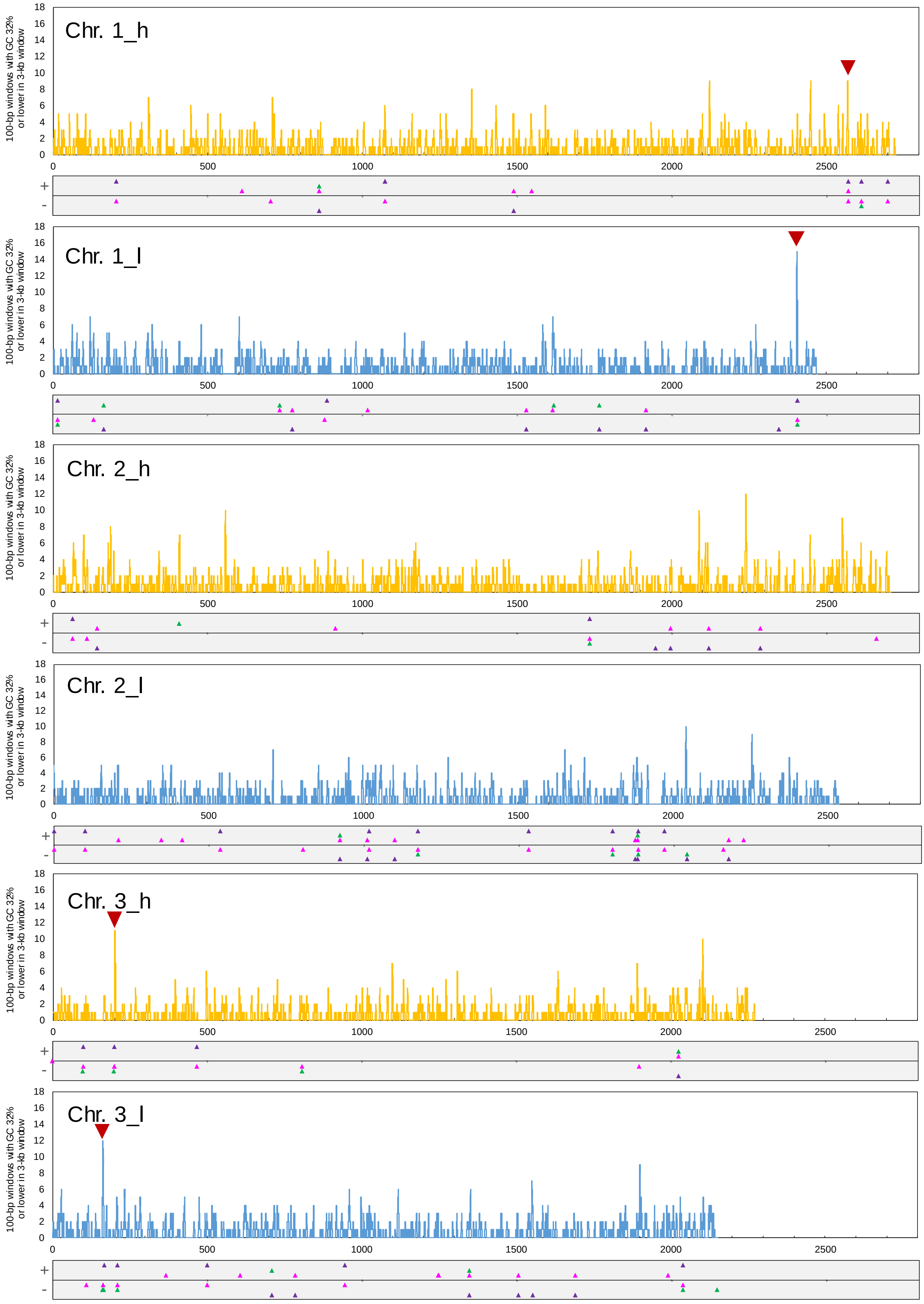
Chr. 1_h
Chr. 1_l
Chr. 2_h
Chr. 2_l
Chr. 3_h
Chr. 3_l
100-bp windows with GC 32% or lower in 3-kb window
0
2
4
6
8
10
12
14
16
18
0
500
1000
1500
2000
2500
+
-

(continued)

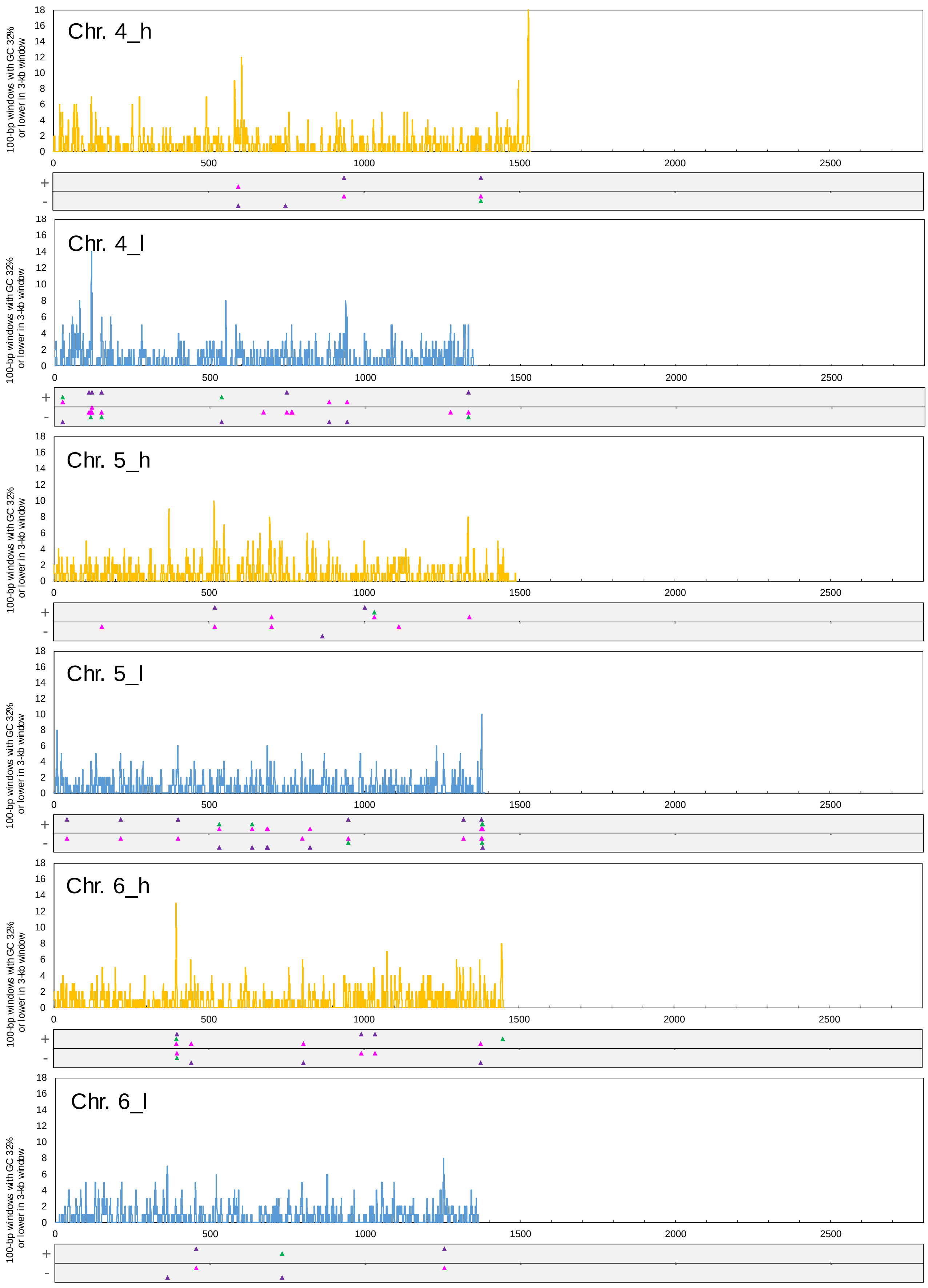
Chr. 4_h
Chr. 4_l
Chr. 5_h
Chr. 5_l
Chr. 6_h
Chr. 6_l
100-bp windows with GC 32% or lower in 3-kb window
0
500
1000
1500
2000
2500
0
2
4
6
8
10
12
14
16
18
+
-

(continued)

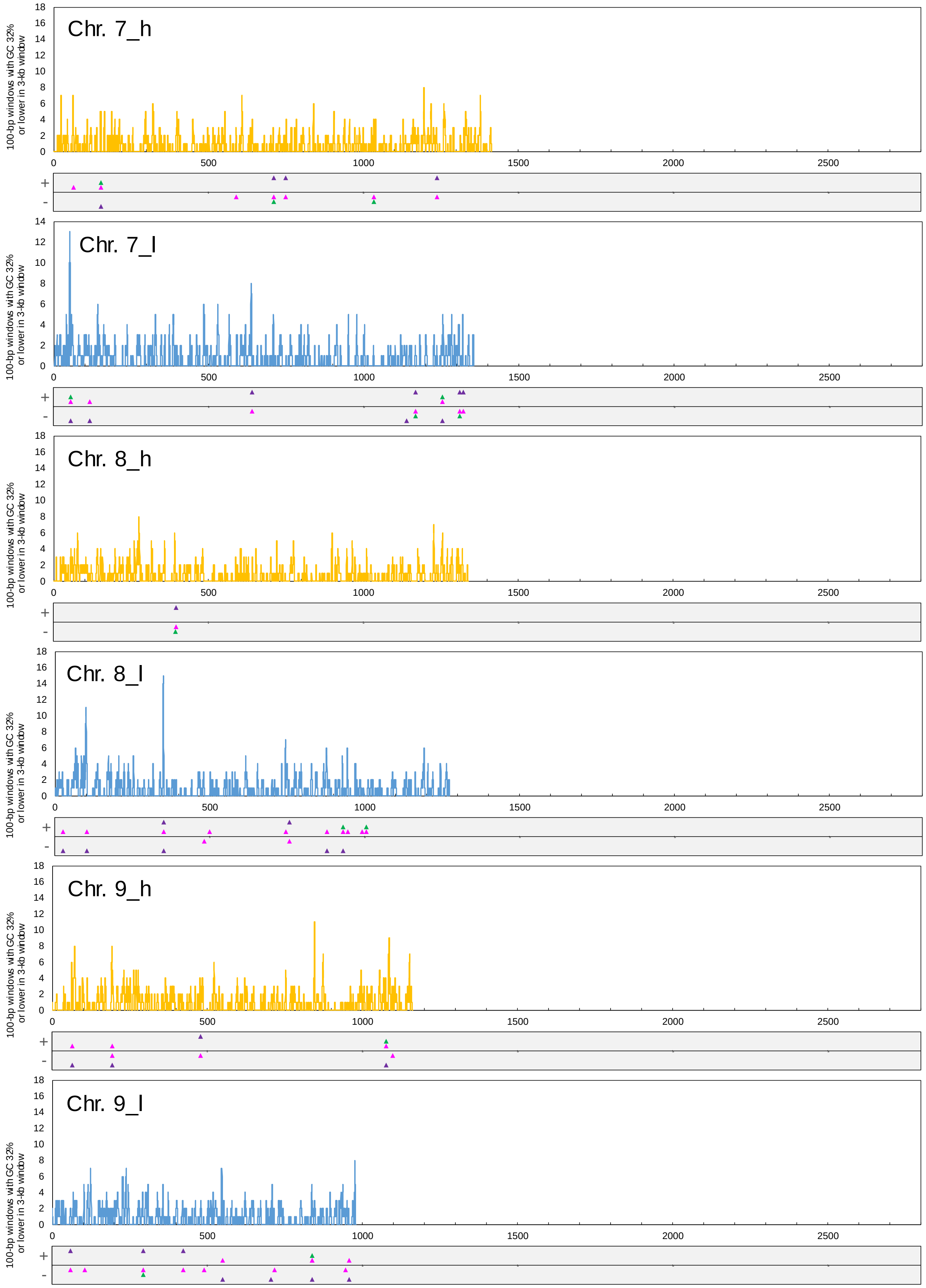

Chr. 7_h
Chr. 7_l
Chr. 8_h
Chr. 8_l
Chr. 9_h
Chr. 9_l
100-bp windows with GC 32% or lower in 3-kb window
0
500
1000
1500
2000
2500
+
-

(continued)

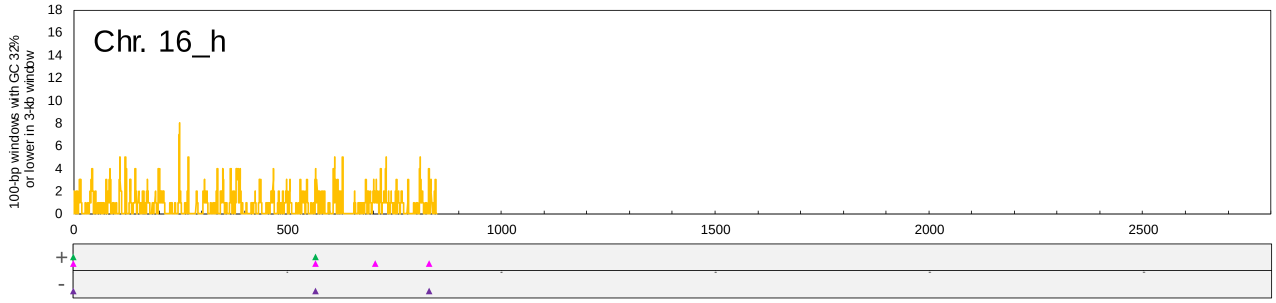
Chr. 16_h
100-bp windows with GC 32% or lower in 3-kb window
0 2 4 6 8 10 12 14 16 18
0 500 1000 1500 2000 2500
+
-

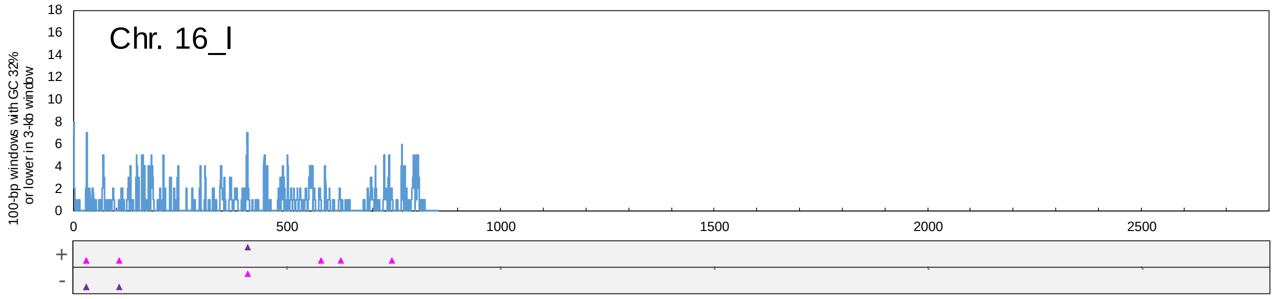
Chr. 16_l
100-bp windows with GC 32% or lower in 3-kb window
0 2 4 6 8 10 12 14 16 18
0 500 1000 1500 2000 2500
+
-

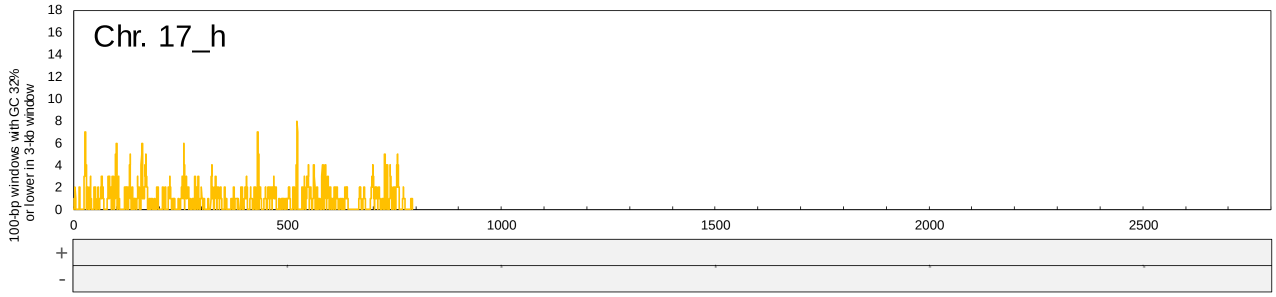
Chr. 17_h
100-bp windows with GC 32% or lower in 3-kb window
0 2 4 6 8 10 12 14 16 18
0 500 1000 1500 2000 2500
+
-

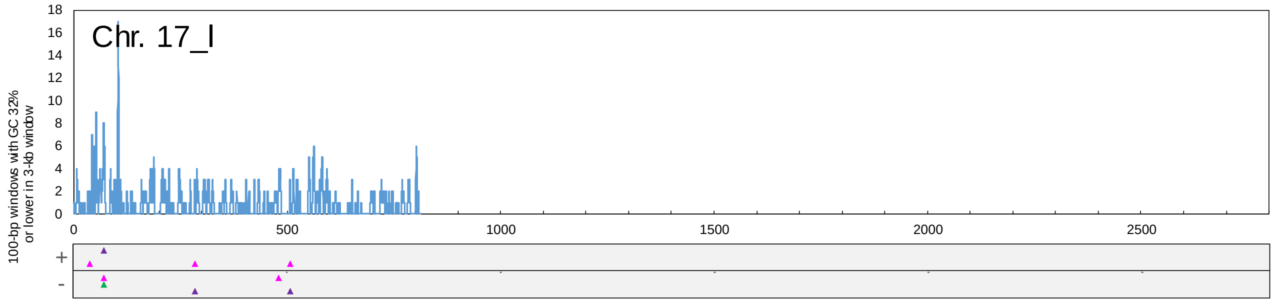
Chr. 17_l
100-bp windows with GC 32% or lower in 3-kb window
0 2 4 6 8 10 12 14 16 18
0 500 1000 1500 2000 2500
+
-

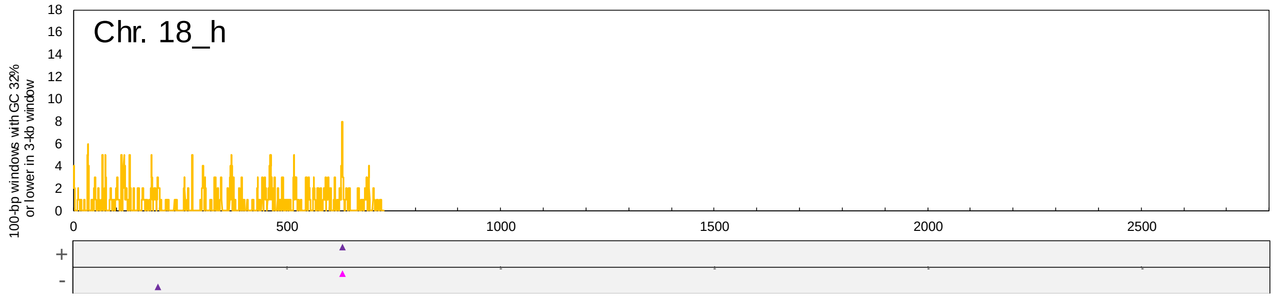
Chr. 18_h
100-bp windows with GC 32% or lower in 3-kb window
0 2 4 6 8 10 12 14 16 18
0 500 1000 1500 2000 2500
+
-

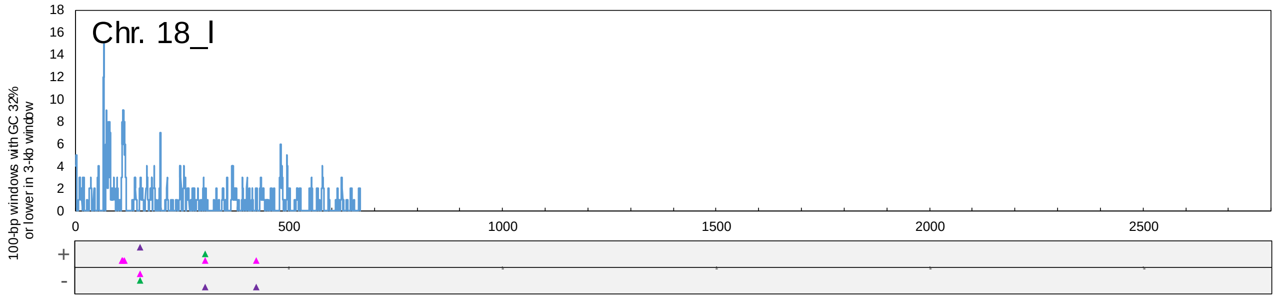
Chr. 18_l
100-bp windows with GC 32% or lower in 3-kb window
0 2 4 6 8 10 12 14 16 18
0 500 1000 1500 2000 2500
+
-

(continued)

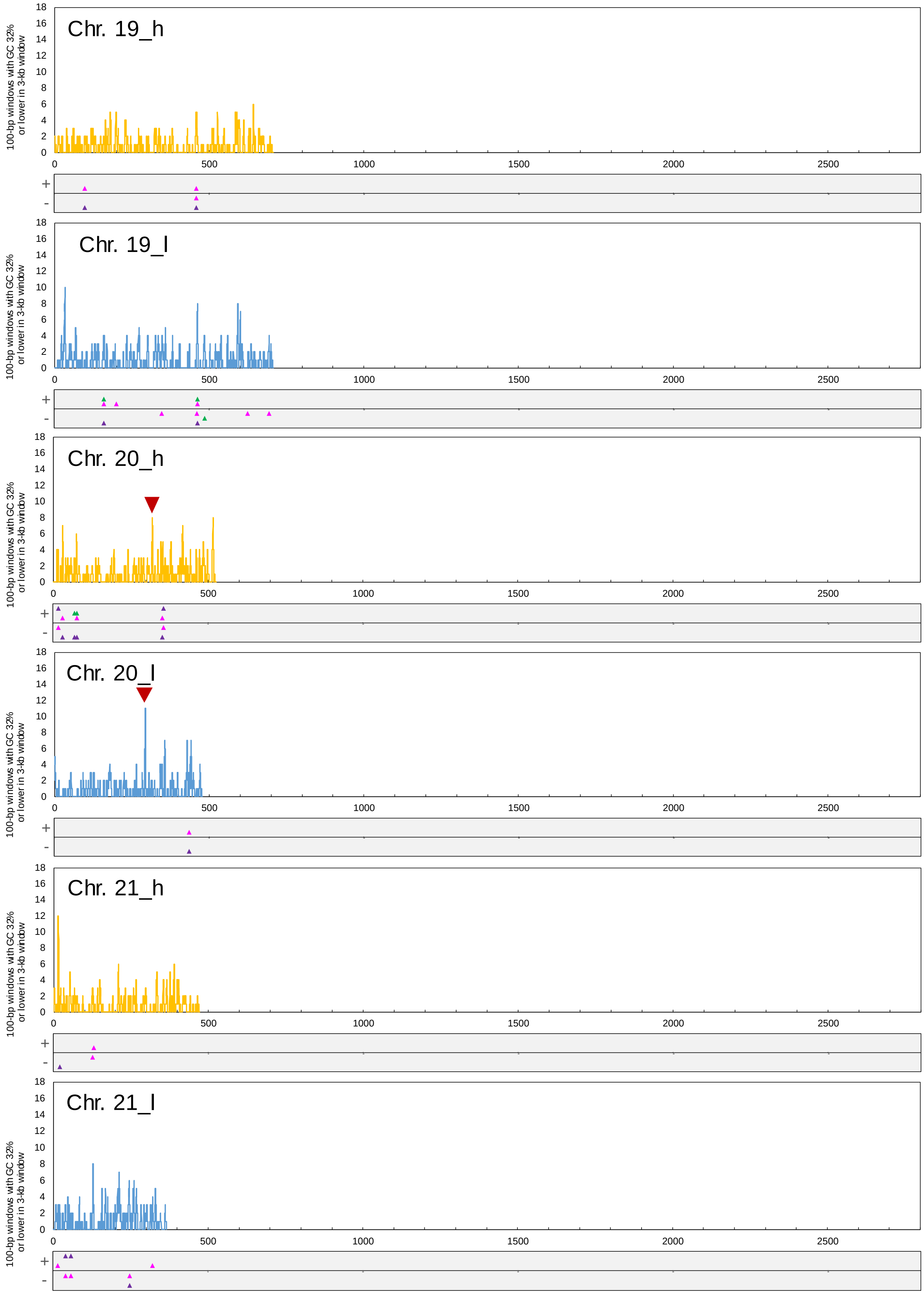
Chr. 19_h
Chr. 19_l
Chr. 20_h
Chr. 20_l
Chr. 21_h
Chr. 21_l
100-bp windows with GC 32% or lower in 3-kb window
0
2
4
6
8
10
12
14
16
18
0
500
1000
1500
2000
2500
+
-

(continued)

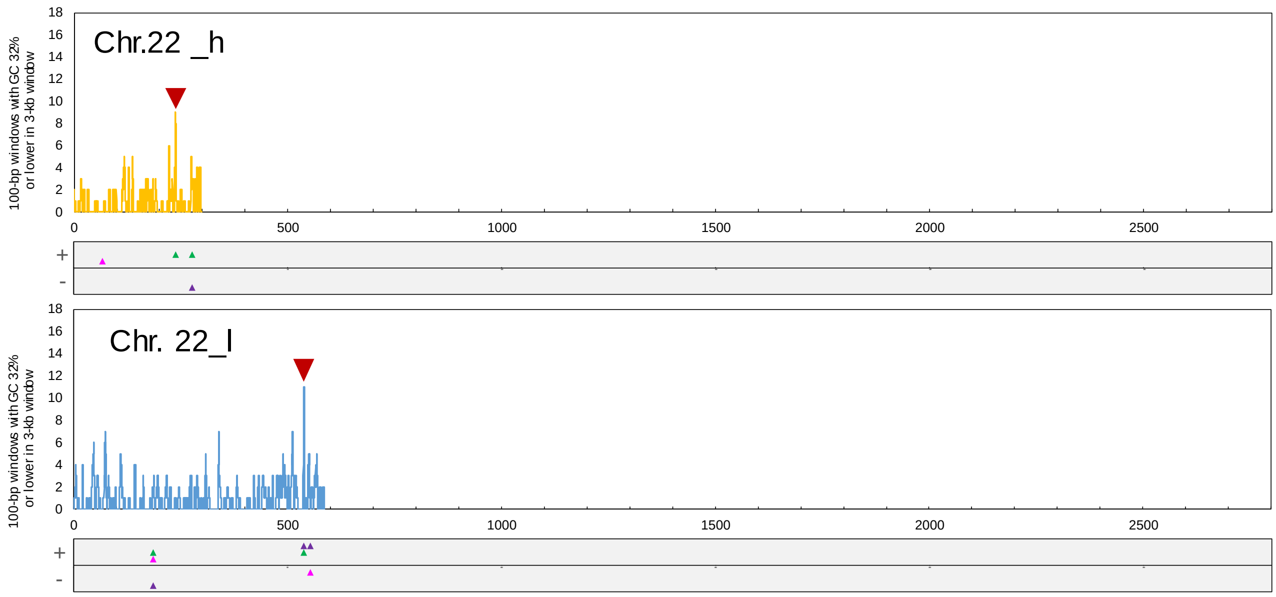

Fig. S5 Putatively identified centromere regions in the *F. solaris* chromosomes. GC content was calculated for the chromosomes in 100-bp sliding windows that overlapped by 50 bp. Numbers of 100-bp windows with GC content of 32% or lower within a larger sliding 3-kb window are plotted along the chromosomes. The putatively identified centromeres are shown by red triangles. The distributions of the motifs 1, 2, and 3 shown in Figure 3 in the main text along the positive (+) and negative (-) strands of the chromosomes are shown by pink, green and purple triangles.

Chr. 1 Fso_h

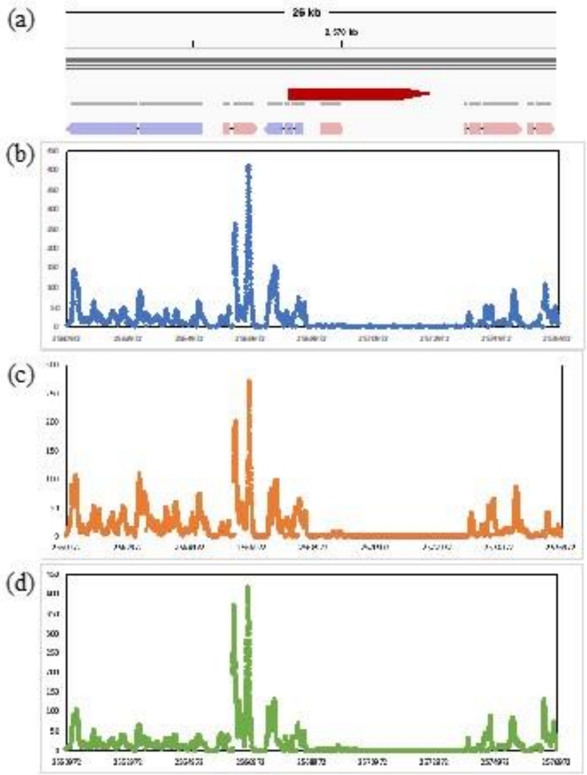

Chr. 1 Fso_l

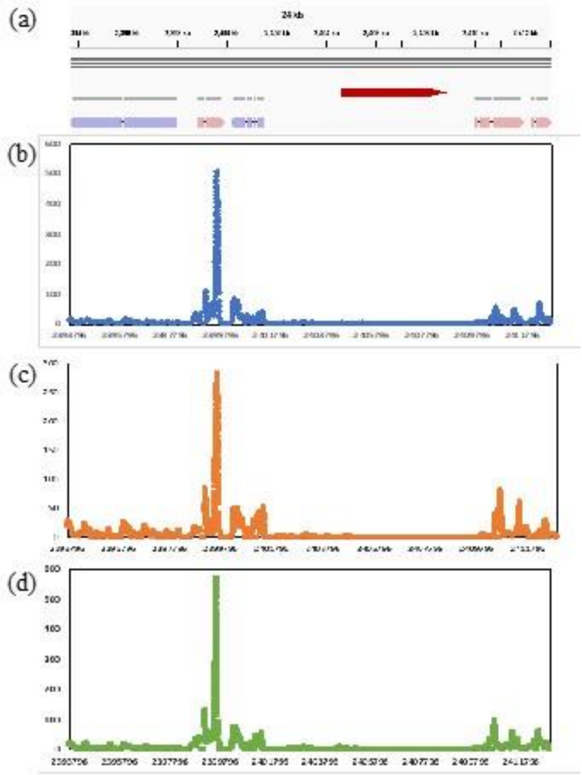

Chr. 3 Fso_h

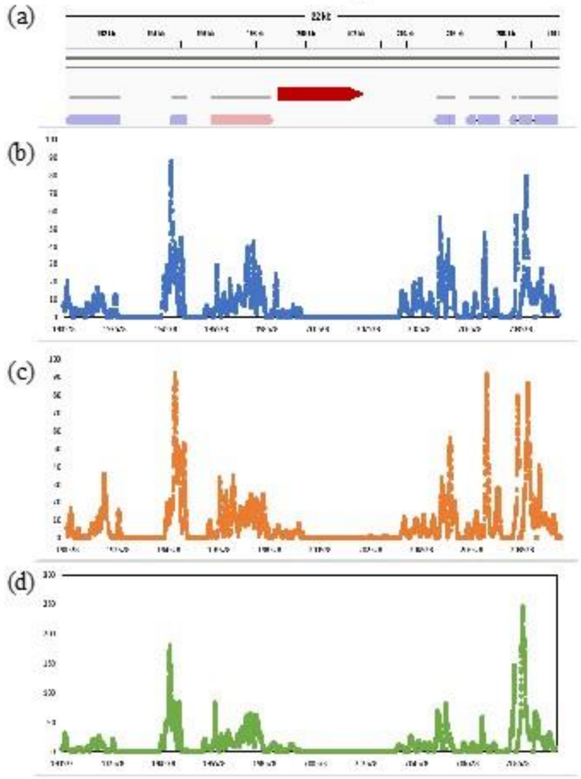

Chr. 3 Fso_l

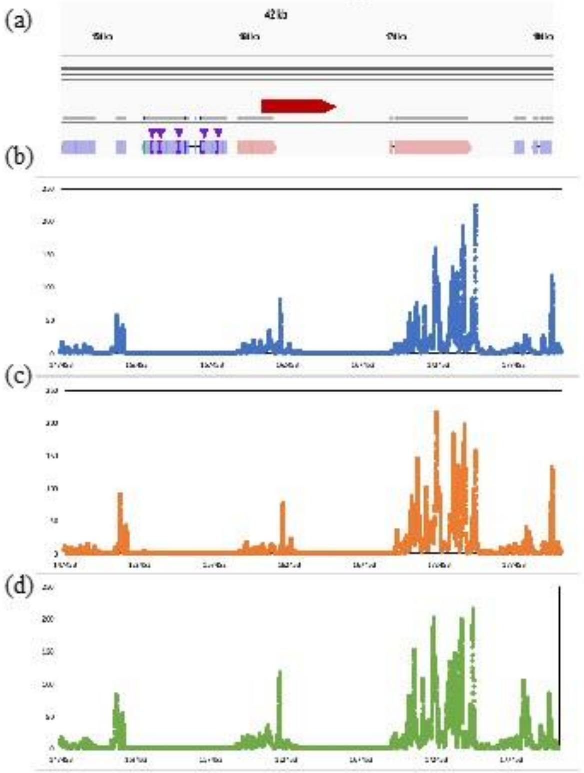

(continued)

#### Chr. 10 Fso_h

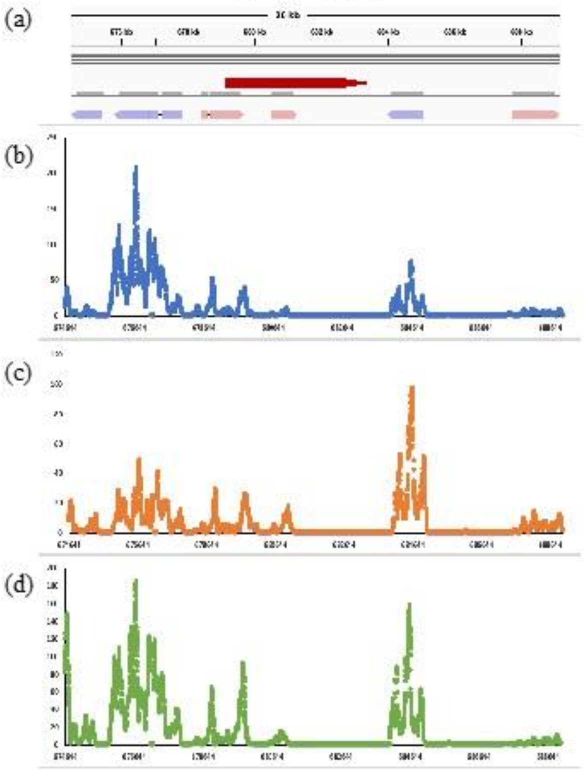

#### Chr. 10 Fso_l

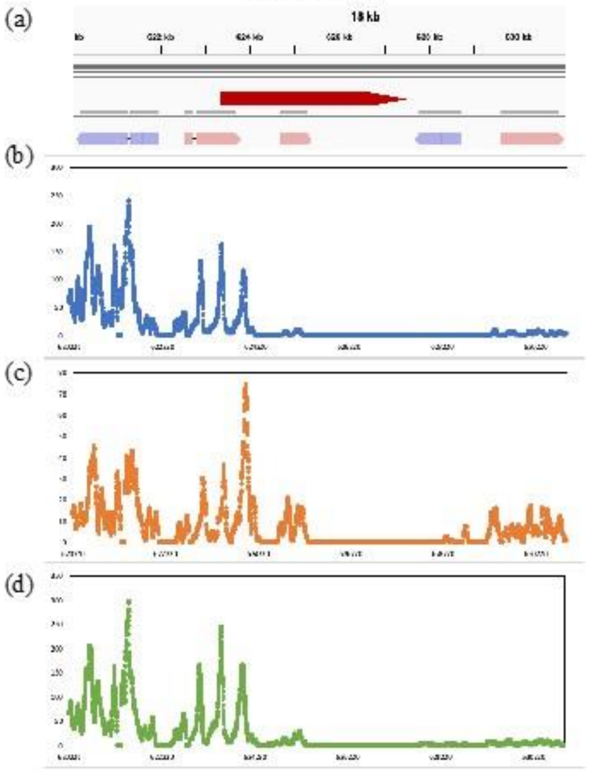

#### Chr. 11 Fso_h

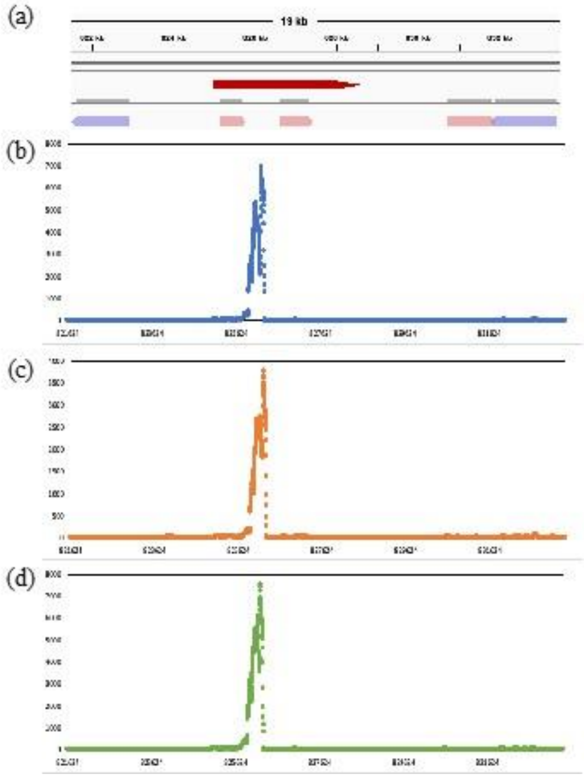

#### Chr. 11 Fso_l

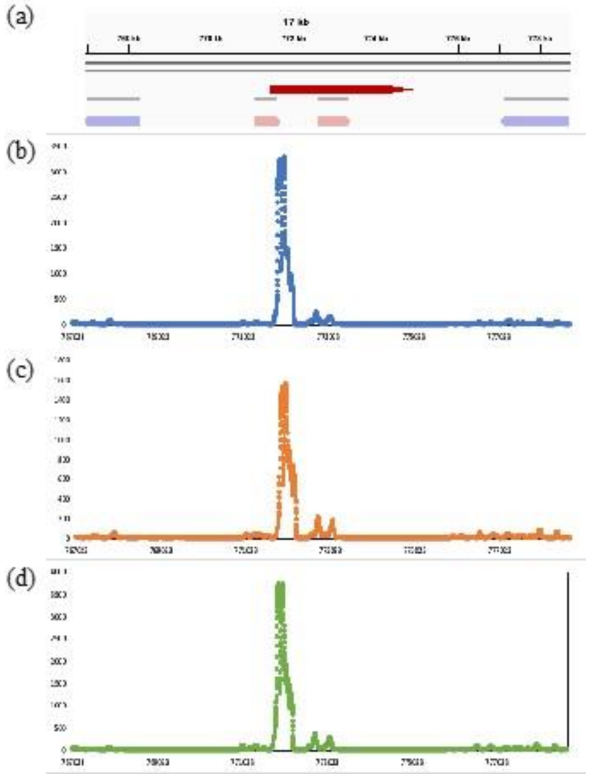

(continued)

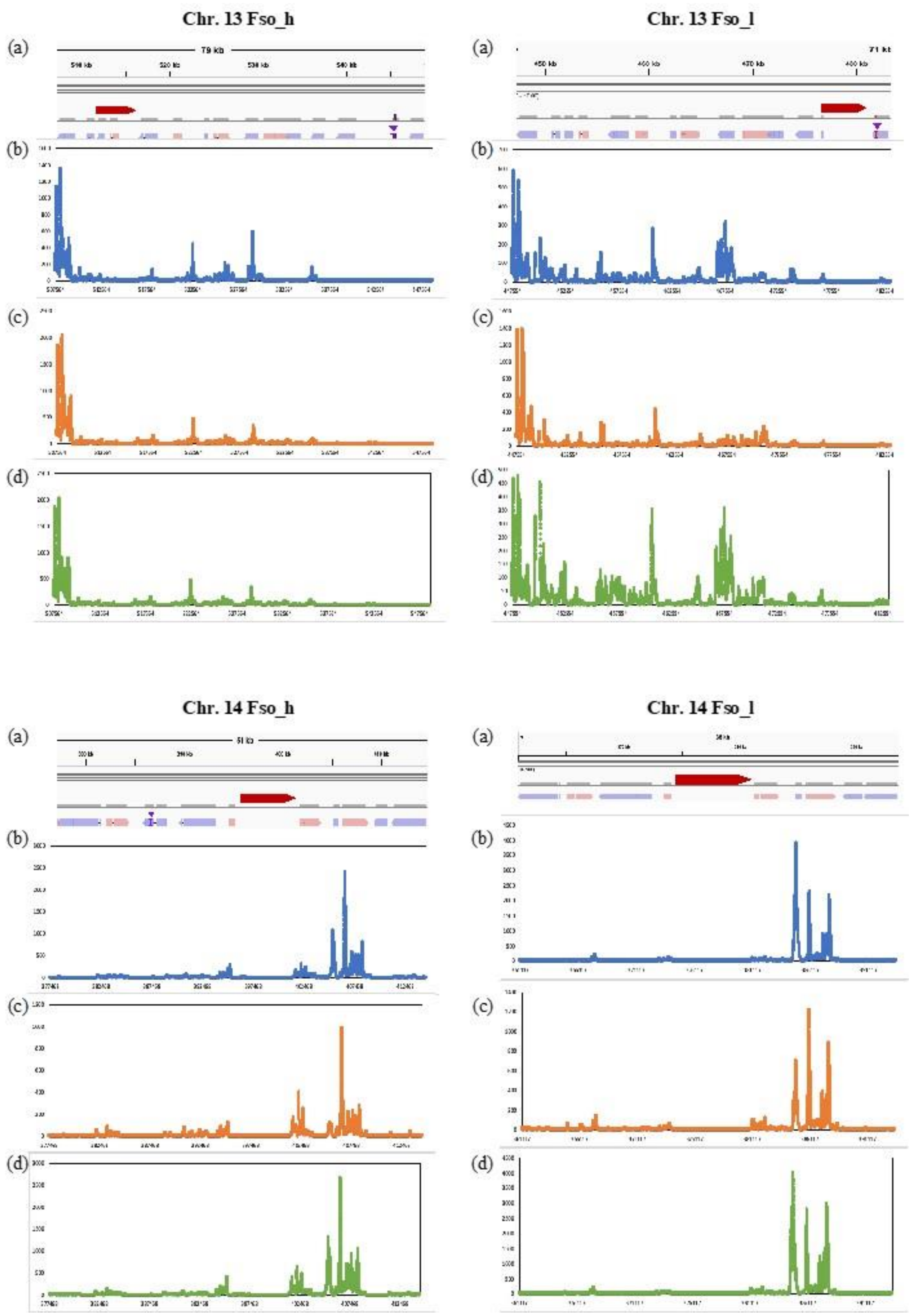

(continued)

**Chr. 20 Fso_h**

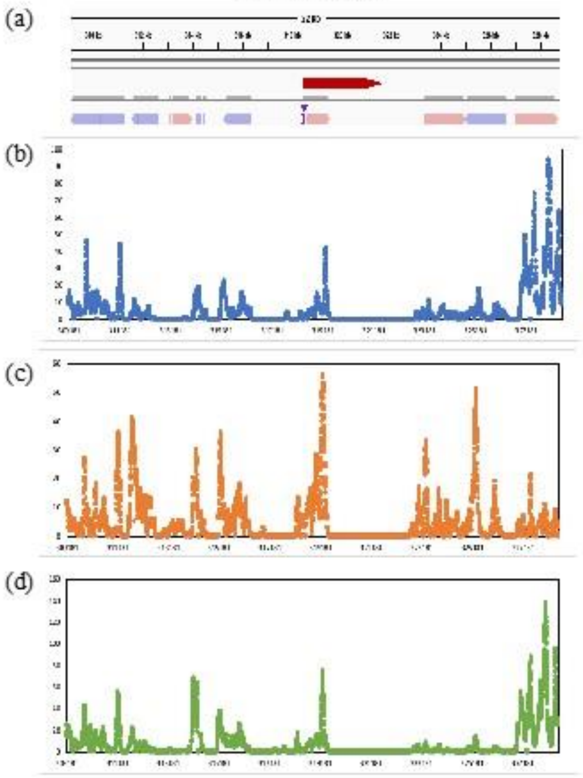

**Chr. 20 Fso_l**

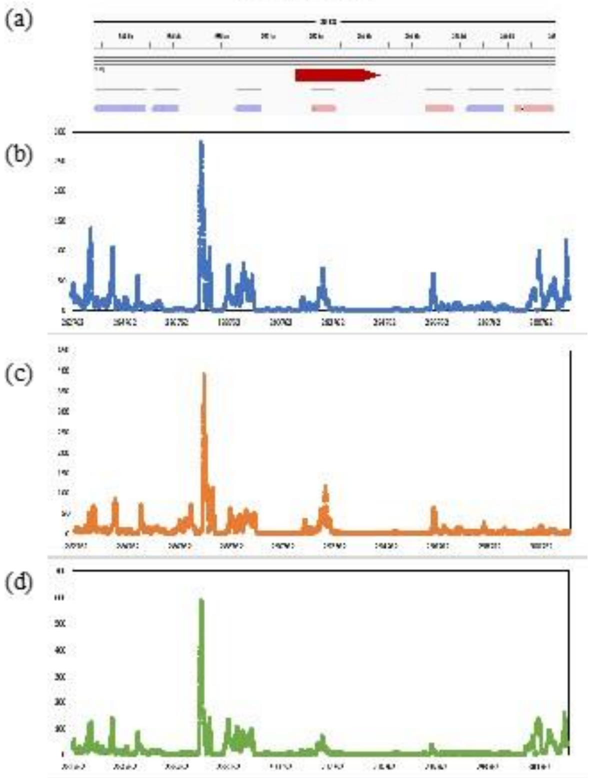

**Chr. 22 Fso_h**

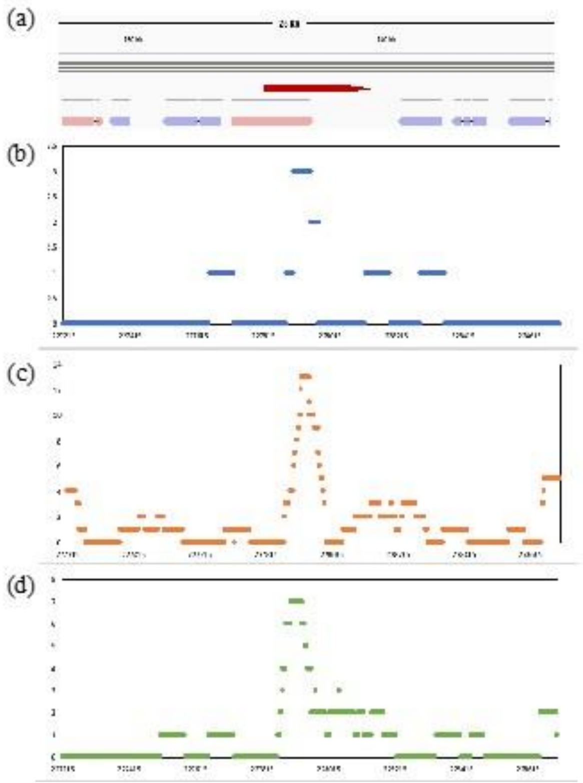

**Chr. 22 Fso_l**

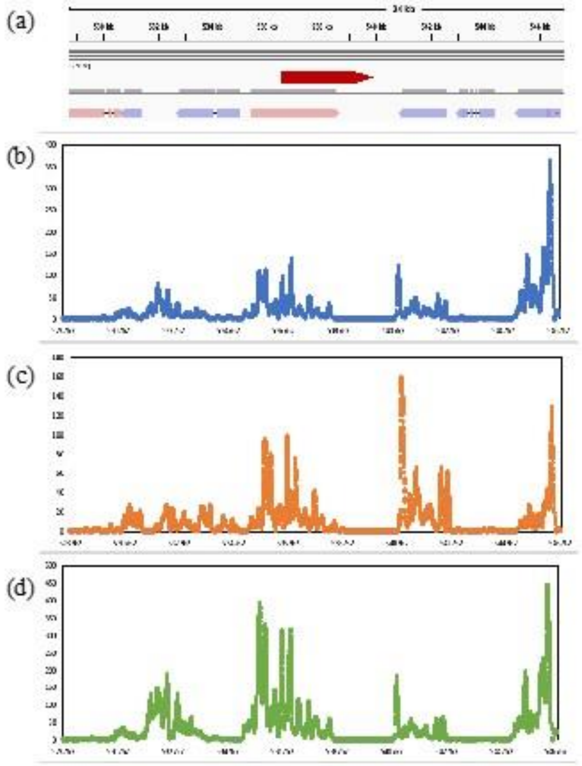

(continued)

Fig. S6 Positional relationships between the gene regions (blue and pink arrows) and the regions containing the putative centromeres (red arrows) in the MinION contigs (a). These regions show high numbers of 100-bp windows with GC content of 32% or lower within a larger sliding 3-kb window. (b, c, d) Mapping depths of the transcriptome reads obtained from *Fistulifera solaris* cultivated for 48, 96, and 144 h in f/2 medium in our previous study. The putative centromeres determined by excluding the transcribed regions from the red arrows.

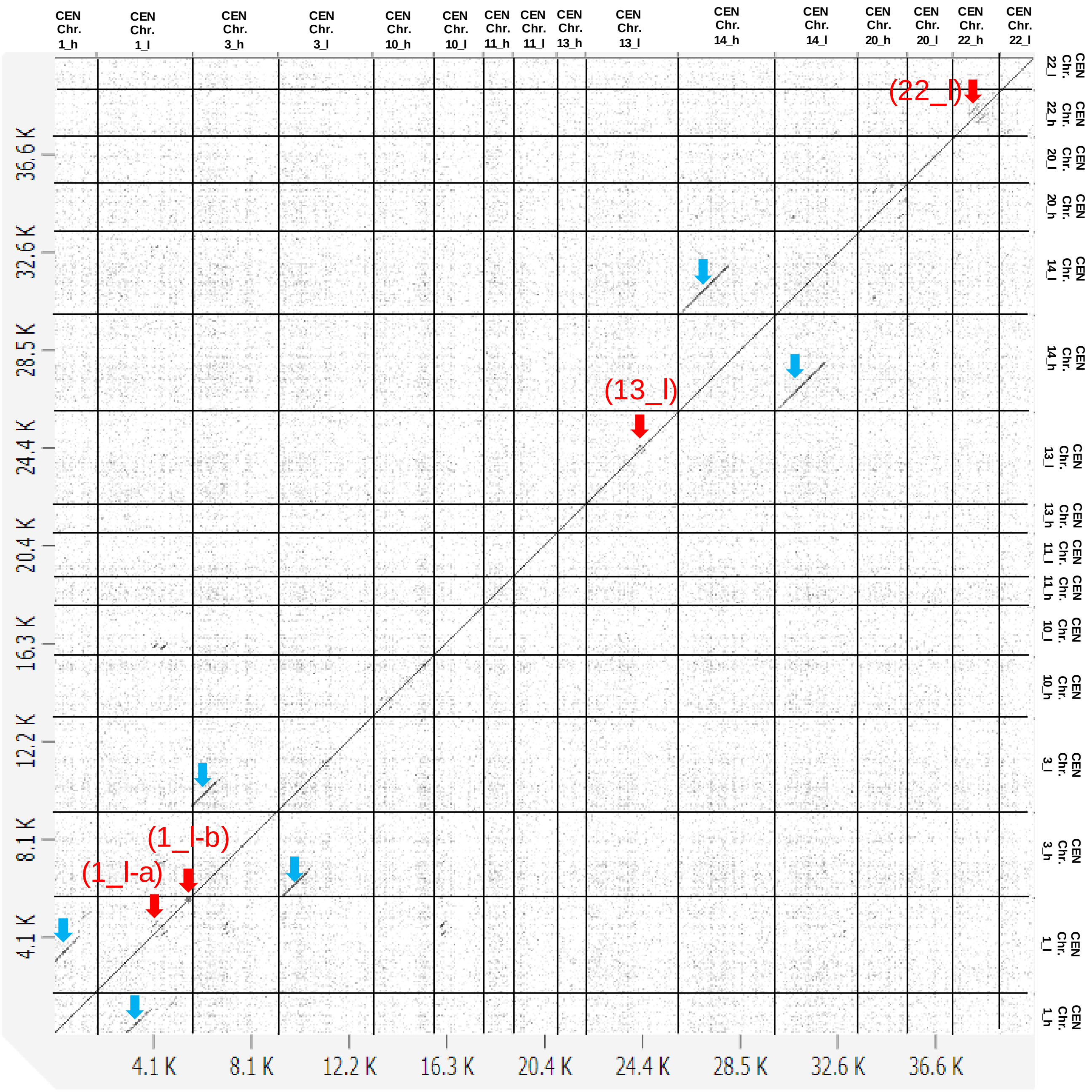

Fig. S7 Dot plot analysis by self-alignment of the putatively identified centromere sequences. Red arrows indicate the small-scale repeating regions. Light blue arrows indicate the regions showing sequence similarity to their homoeologous counterparts.

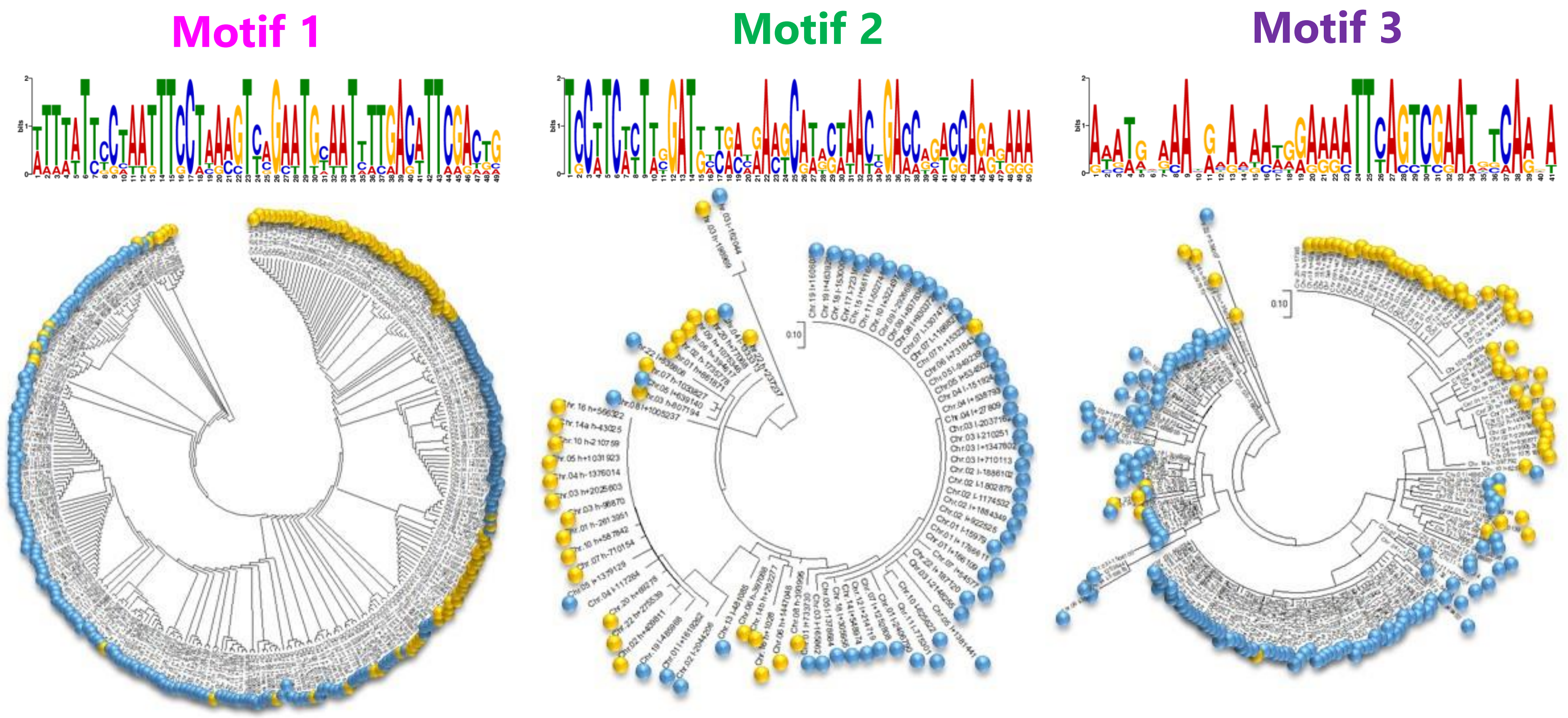

Fig. S8 Phylogenetic analysis of the conserved motifs of the potential autonomous replication sequences in *Fistulifera solaris* genome. Conserved motifs 1, 2, and 3 found from the centromere regions were explored from the entire genome sequence as shown in Supplementary Figure S5. The found sequences of each motif were subjected to the phylogenetic analysis using MEGA-X with the neighbor-joining method under the default setting. The sequences found from the subgenomes Fso_h and Fso_l are shown in orange and blue circles.

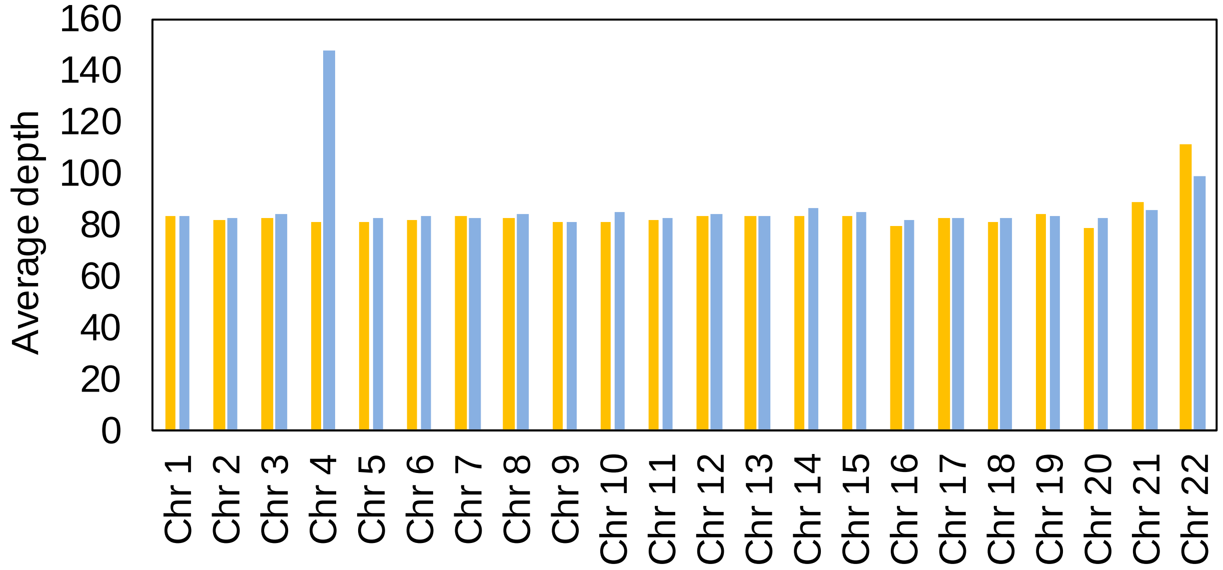

Fig. S9 Average read depth of the MinION contigs of the *Fistulifera solaris* allopolyploid genome. Orange and blue bards represent the contigs belonging to Fso_h and Fso_l subgenomes, respectively

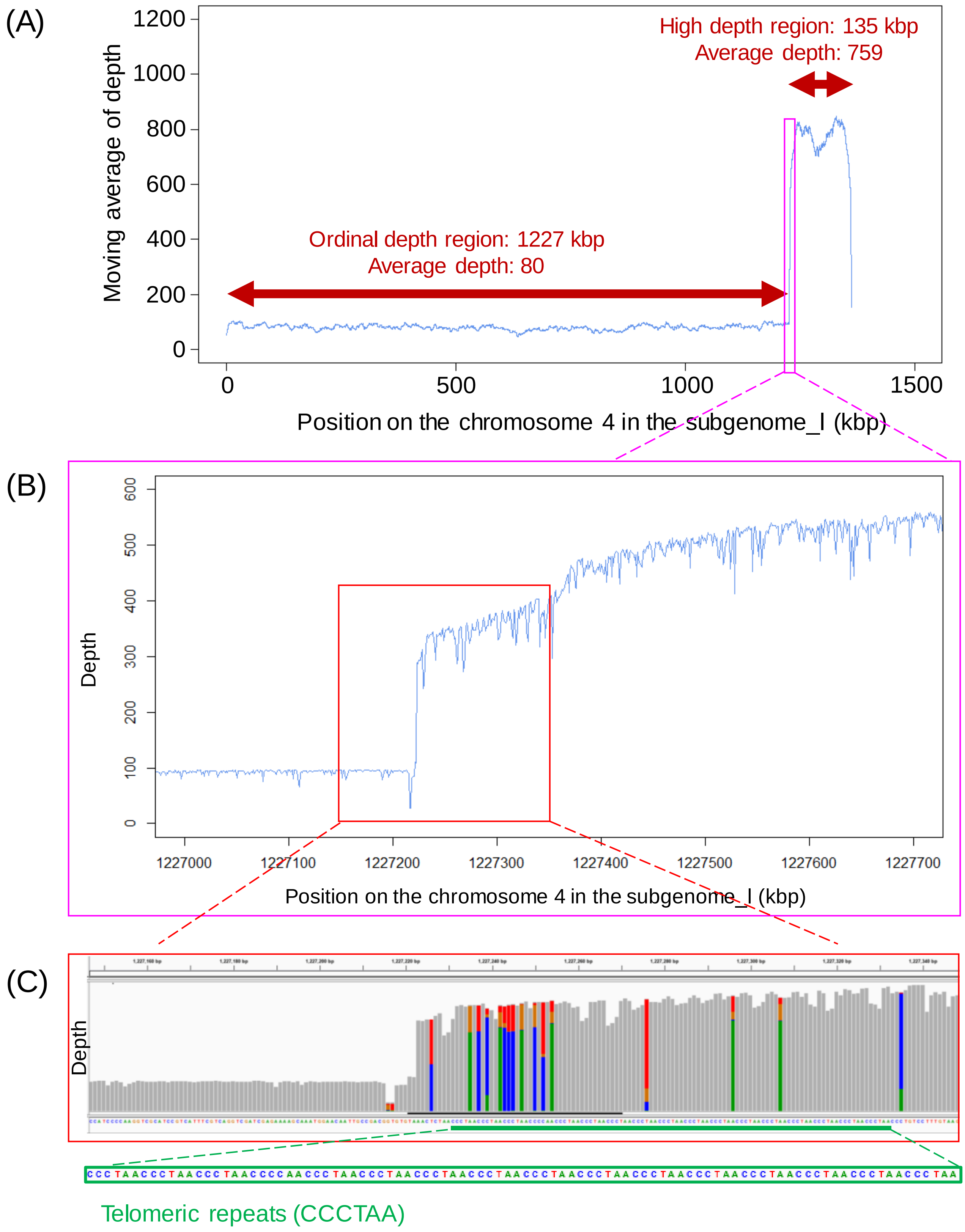

Fig. S10 Read depths along the chromosome 4 in the subgenome Fso_I (A). The magnified image of the pink region in (A) indicates the sudden increase in the read depths (B). The magnified image of the red region indicates the co-existence of two types of subreads one of which contains telomeric repeats (CCCTAA) (C).

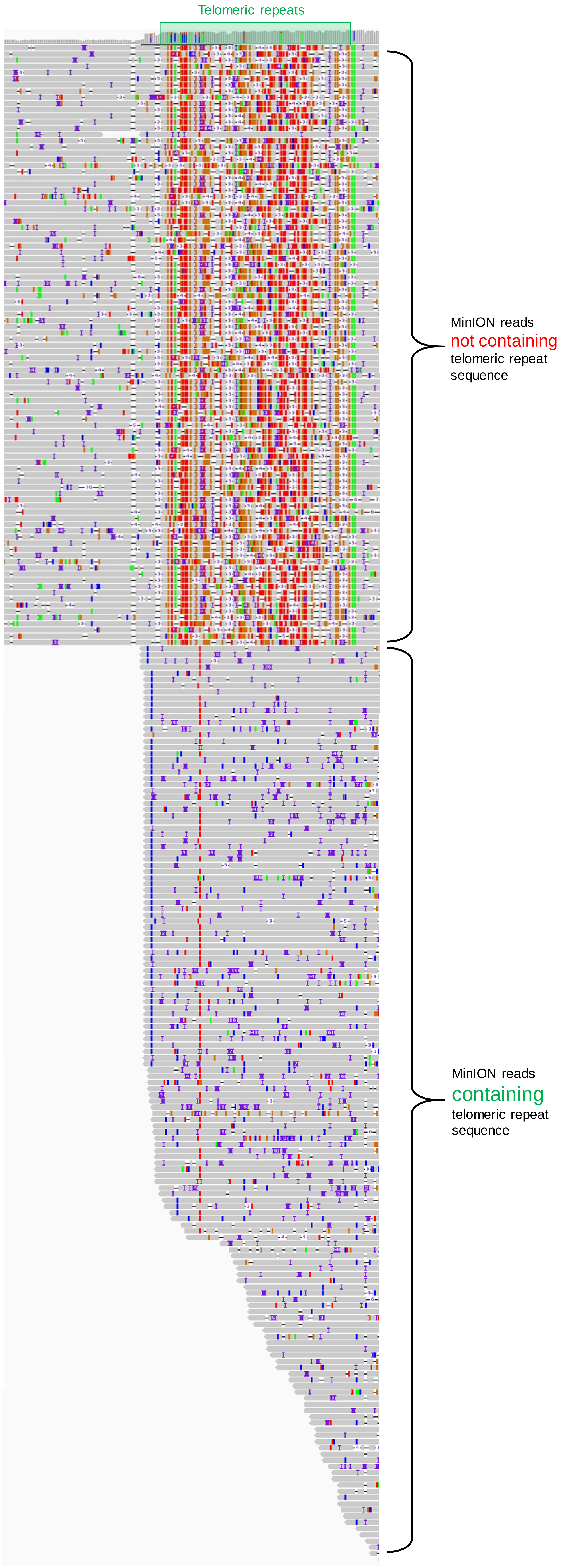

Fig. S11 An alignment result of the MinION reads onto the contig (corresponding to chromosome 4 in the subgenome Fso_I) generated by the assembly of the MinION reads. The MinION reads can be classified into two classes, one of which contain telemetric repeats sequence and others do not contain them.

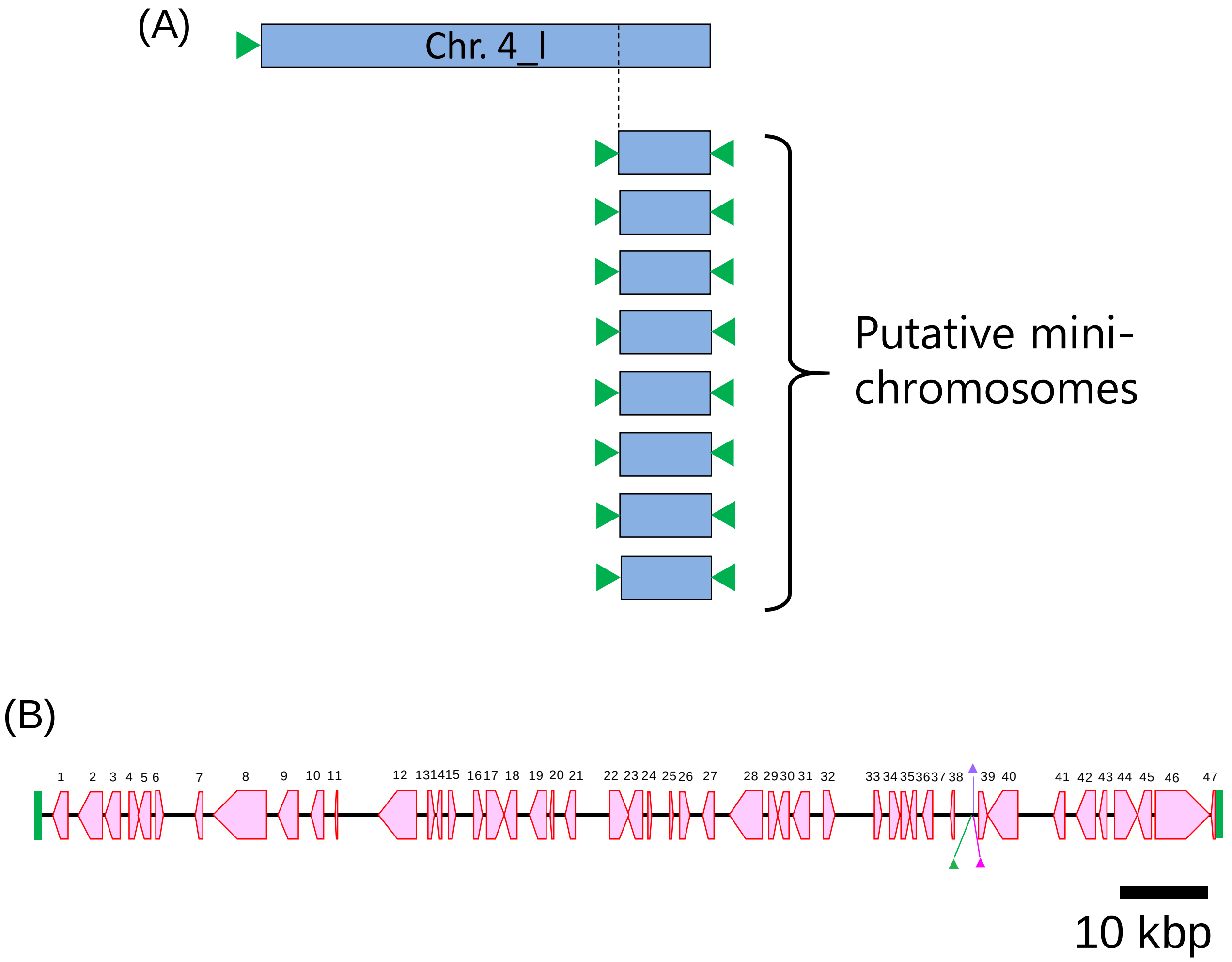

Fig. S12 Schematic diagrams of the chromosomes 4 belonging to the subgenome Fso_I (Chr. 4_I), and mini-chromosomes (A), which contain 47 genes (pink arrows) between the telomere sequences (B). Green, purple, and pink triangles in panel (B) represent the autonomous replication sequences shown in Supplementary Figure S5. Numbers above the genes correspond to the gene order shown in Supplementary Table S4.

### Table S4 The genes on the mini-chromosomes in chromosome 4_I.

| Order on Mini-Chr. 4_I | Gene ID | Protein ID | Protein function | Predicted domain | Protein size (AA) |
|---|---|---|---|---|---|
| 1 | fso:g1828 | GAX11032.1 | 6-phosphofructokinase 1 | pyrophosphate-dependent phosphofructose kinase | 545 |
| 2 | fso:g1829 | GAX11033.1 | chloride channel 7 | voltage activated chloride channel CLC7 type | 863 |
| 3 | fso:g1830 | GAX11034.1 | cytochrome P450 family 26 subfamily A | cytochrome P450 | 525 |
| 4 | fso:g1831 | GAX11035.1 | hypothetical protein FisN_2Lh564 | ---NA--- | 353 |
| 5 | fso:g1832 | GAX11036.1 | creatine kinase | arginine kinase | 448 |
| 6 | fso:g1833 | GAX11037.1 | hypothetical protein FisN_2Lh566 | ---NA--- | 299 |
| 7 | fso:g1834 | GAX11038.1 | hypothetical protein FisN_2Lu567 | structural maintenance of chromosomes 3 | 288 |
| 8 | fso:g1835 | GAX11039.1 | erbb2-interacting protein | L domain | 2073 |
| 9 | fso:g1836 | GAX11040.1 | hypothetical protein FisN_2Lh569 | predicted protein | 467 |
| 10 | fso:g1837 | GAX11041.1 | hypothetical protein FisN_2Lh570 | BTB POZ domain-containing | 450 |
| 11 | fso:g1838 | GAX11042.1 | hypothetical protein FisN_2Lu571 | hypothetical protein FRACYDRAFT:268161 | 103 |
| 12 | fso:g1351 | GAX10557.1 | hypothetical protein FisN_2Lh591 | ---NA--- | 237 |
| 13 | fso:g1352 | GAX10558.1 | hypothetical protein FisN_2Lh590 | hypothetical protein FRACYDRAFT:185991 | 192 |
| 14 | fso:g1353 | GAX10559.1 | hypothetical protein FisN_2Lu589 | Transmembrane 19 | 292 |
| 15 | fso:g1354 | GAX10560.1 | protein MAK16 | RNA-binding nuclear (MAK16) containing a distinct C4 Zn-finger | 312 |
| 16 | fso:g1355 | GAX10561.1 | hypothetical protein FisN_2Lh587 | nucleotide transporter 3 | 624 |
| 17 | fso:g1356 | GAX10562.1 | hypothetical protein FisN_2Lh586 | ATPase AFG2 | 516 |
| 18 | fso:g1357 | GAX10563.1 | hypothetical protein FisN_2Lh585 | ---NA--- | 572 |
| 19 | fso:g1358 | GAX10564.1 | hypothetical protein FisN_2Lu584 | ---NA--- | 117 |
| 20 | fso:g1359 | GAX10565.1 | hypothetical protein FisN_2Lh583 | S-adenosyl-L-methionine-dependent methyltransferase | 318 |
| 21 | fso:g13571 | GAX22741.1 | hypothetical protein FisN_2Lh582 | beta-glucan synthase | 684 |
| 22 | fso:g13572 | GAX22742.1 | hypothetical protein FisN_2Lh581 | nucleotide transporter 3 | 547 |
| 23 | fso:g13573 | GAX22743.1 | hypothetical protein FisN_2Lh580 | ---NA--- | 195 |
| 24 | fso:g1822 | GAX11026.1 | 21S rRNA (GM2251-2'-O)-methyltransferase | ribose methyltransferase | 422 |
| 25 | fso:g13574 | GAX22744.1 | hypothetical protein FisN_2Lh579 | ---NA--- | 168 |
| 26 | fso:g13575 | GAX22745.1 | hypothetical protein FisN_2Lh578 | predicted protein | 355 |
| 27 | fso:g13576 | GAX22746.1 | hypothetical protein FisN_2Lh577 | Dynein heavy chain axonemal | 398 |
| 28 | fso:g13577 | GAX22747.1 | hypothetical protein FisN_2Lh576 | Tetratricopeptide-like helical | 1193 |
| 29 | fso:g13578 | GAX22748.1 | hypothetical protein FisN_2Lh575 | predicted protein | 367 |
| 30 | fso:g13579 | GAX22749.1 | large subunit ribosomal protein L3 | 50S ribosomal L3 | 378 |
| 31 | fso:g13580 | GAX22750.1 | hypothetical protein FisN_2Lh573 | ---NA--- | 592 |
| 32 | fso:g13581 | GAX22751.1 | hypothetical protein FisN_2Lh572 | ---NA--- | 414 |
| 33 | fso:g1859 | GAX11063.1 | hypothetical protein FisN_2Lh606 | ---NA--- | 290 |
| 34 | fso:g1860 | GAX11064.1 | hypothetical protein FisN_2Lh605 | ---NA--- | 367 |
| 35 | fso:g1861 | GAX11065.1 | tRNA acetyltransferase TAN1 | thump domain-containing 1-like | 317 |
| 36 | fso:g1862 | GAX11066.1 | hypothetical protein FisN_2Lh603 | Ankyrin repeat-containing domain | 213 |
| 37 | fso:g1863 | GAX11067.1 | hypothetical protein FisN_2Lh602 | hypothetical protein THAOC:34853, partial | 379 |
| 38 | fso:g1864 | GAX11068.1 | hypothetical protein FisN_2Lu601 | rRNA adenine dimethylase | 176 |
| 39 | fso:g1865 | GAX11069.1 | transcription elongation factor S-II | transcription elongation factor S-II | 309 |
| 40 | fso:g1866 | GAX11070.1 | hypothetical protein FisN_2Lh599 | ---NA--- | 1162 |
| 41 | fso:g1867 | GAX11071.1 | hypothetical protein FisN_2Lh598 | ---NA--- | 372 |
| 42 | fso:g1868 | GAX11072.1 | hypothetical protein FisN_2Lh597 | DNA repair family | 714 |
| 43 | fso:g1869 | GAX11073.1 | hypothetical protein FisN_2Lh596 | predicted protein | 290 |
| 44 | fso:g1870 | GAX11074.1 | ribonucleoside-diphosphate reductase subunit M1 | ribonucleoside-diphosphate reductase large subunit | 800 |
| 45 | fso:g1871 | GAX11075.1 | hypothetical protein FisN_2Lh594 | predicted protein | 459 |
| 46 | fso:g1872 | GAX11076.1 | hypothetical protein FisN_2Lh593 | D -box ATP-dependent RNA helicase D 14 | 2051 |
| 47 | fso:g1873 | GAX11077.1 | peroxiredoxin Q/BCP | peroxiredoxin Q | 189 |

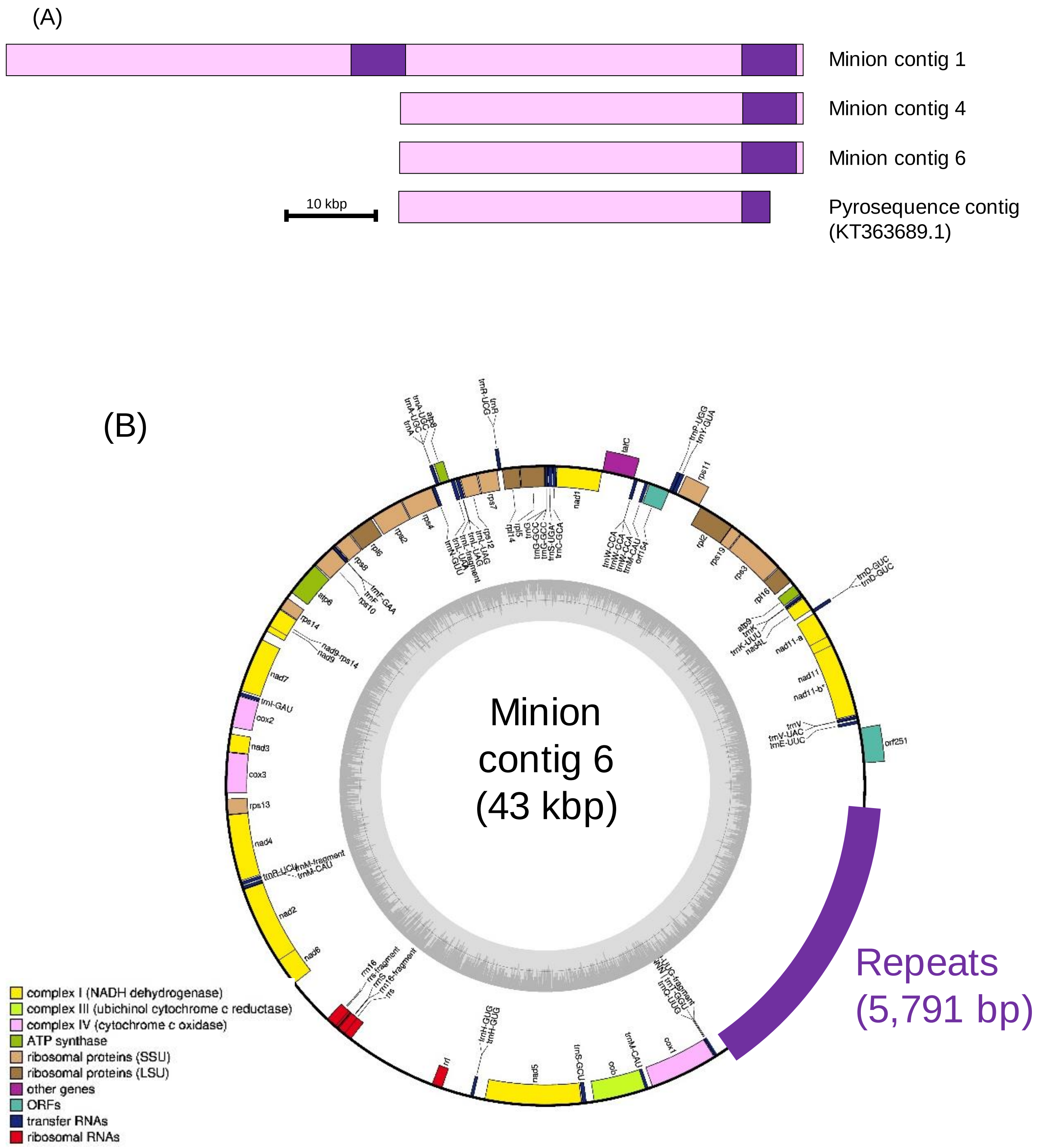

(continued)

(continued)

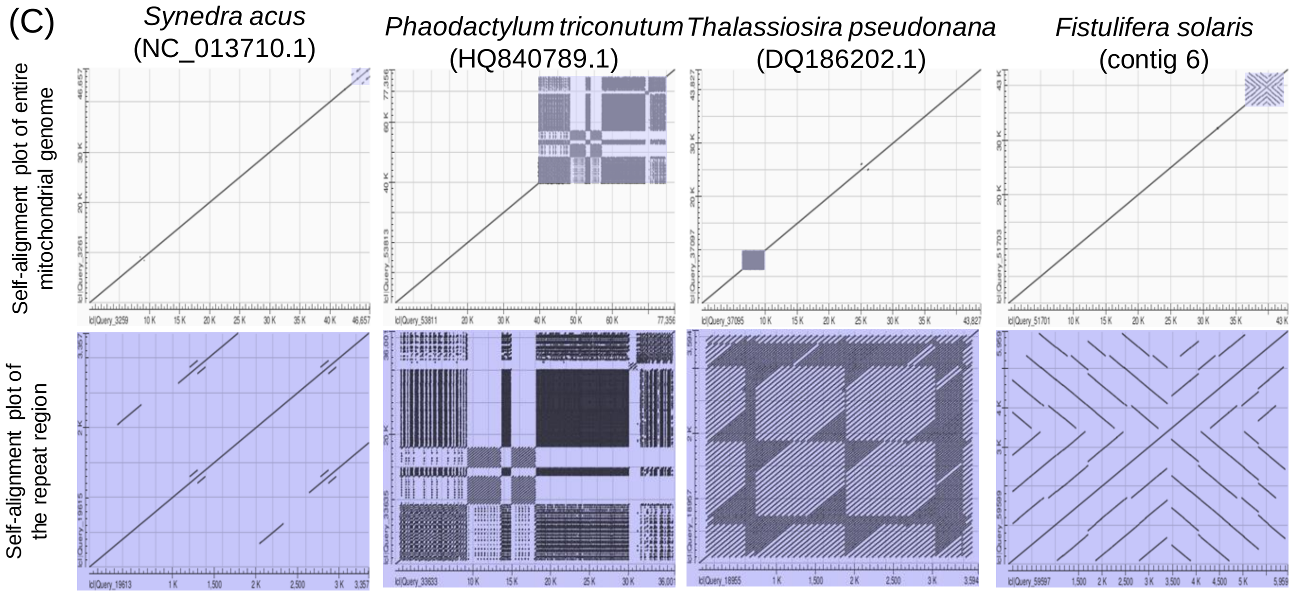

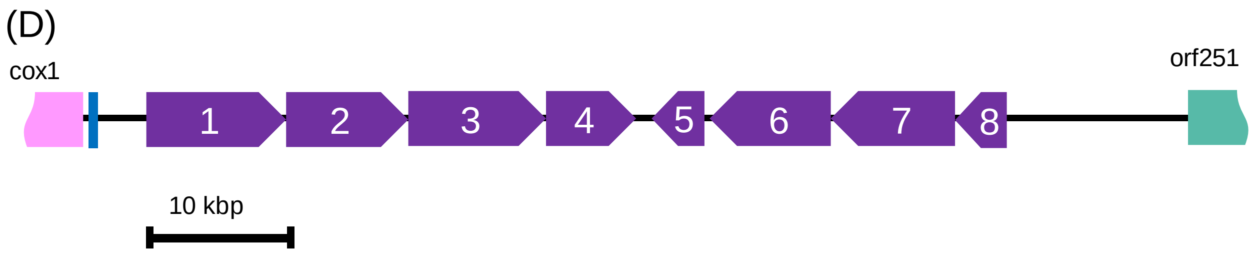

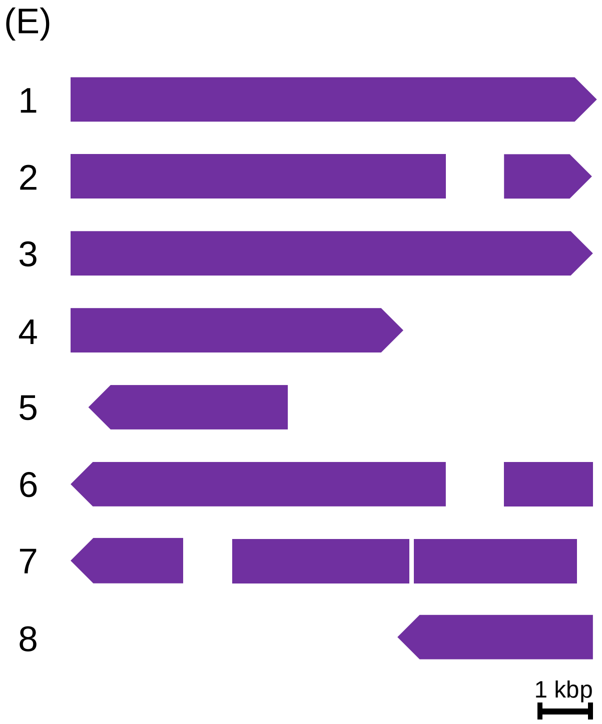

Fig. S13 The contigs corresponding to the mitochondrial genomes of *F. solaris* revealed by nanopore sequencing. (A) Length of 3 MinION contigs and a pyrosequencing scaffold corresponding to the mitochondrial genomes. Purple regions represent the repeat regions. (B) The gene map of the contig 6 generated using GeSeq (Tillich et al. 2017), (C) Dot plot analyses of the repeated regions in the mitochondrial genomes of diatoms. The light purple regions corresponding to the repeated region are shown in the magnified images. The length and arrangements of the repeated elements in the region were shown in (D) and (E).

### PCR tests across and within the repeated regions

PCR amplifications targeting the tandemly repeated *LPAAT* genes (fso:g9516a~e), their neighboring regions, and their homoeologous regions (Supplementary Fig. S13) were performed using the genomic DNA of *F. solaris* as a template and PrimeSTAR GXL DNA polymerase (Takara Bio Inc., Shiga, Japan) with the primers summarized in Supplementary Table S3.

Six target regions (region 1~6 shown in Supplementary Fig. S13) and primer sets used for amplification of each region are shown below.

Region 1: 9515_F and 9516_R
Region 2: 9516_F and 12324_R
Region 3: 8026_F and 8024_R
Region 4: 9515_F and 9516_R2
Region 5: 9516_F2 and 9516_R2
Region 6: 9516_F2 and 12324_R

Primers, 9515_F, 12324_R, 9516_F2 and 9516_R2 were designed to hybridize with the 5' region of fso:g9515, 3' region of fso:g12324, 5' and 3' regions of fso:g8026, respectively, and not to hybridize with their homoeologous genes.

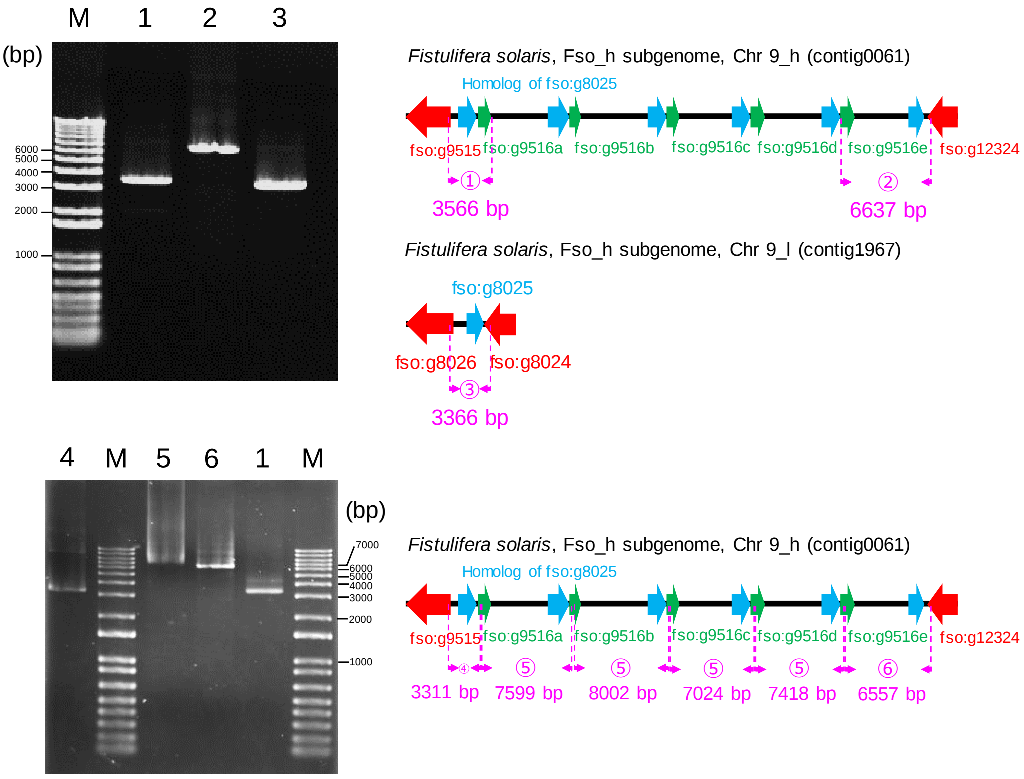

Fig. S14 Agarose gel electrophoresis of the PCR fragments targeting the tandemly repeated LPAAT genes (fso:g9516a~e) and their neighboring regions on the chromosome 9_h (belonging to Fso_h subgenome), and their homoeologous regions on the chromosome 9_l (belonging to Fso_l subgenome). fso:g9515 and fso:g8026, and fso:g12324 and fso:g8024 are homoeologous gene pairs. Lane numbers on the agarose gels and region numbers shown in the schematic diagrams are corresponded. Primer sequences targeting each region are shown in Supplementary Table S3.

(A)

```
fso:g9516a      ATGCTTGCGGCGAATCACAGCTCATGGATGGACACTTTCTATTTGGGTGCGACCGTCGGC    60
fso:g9516b      ATGCTTGCGGCGAATCACAGCTCATGGATGGACACTTTCTATTTGGGTGCGACCGTCGGC    60
fso:g9516e      ATGCTTGCGGCGAATCACAGCTCATGGATGGACACTTTCTATTTGGGTGCGACCGTCGGC    60
fso:g9516c      ATGCTTGTGGCGAATCACAGCTCATGGATGGACACTTTCTATTTGGGTGCGACCGTCGGC    60
fso:g9516d      ATGCTTGTGGCGAATCACAGCTCATGGATGGACACTTTCTATTTGGGTGCGACCGTCGGC    60
                ******* ****************************************************

fso:g9516a      TGGAGAAATTTTAGCCTCATTAGCAAAAATGAAGTGCAGCGCGTCCCTATTCTCGGTAAG    120
fso:g9516b      TGGAGAAATTTTAGCCTCATTAGCAAAAATGAAGTGCAGCGCGTCCCTATTCTCGGTAAG    120
fso:g9516e      TGGAGAAATTTTAGCCTCATTAGCAAAAATGAAGTGCAGCGCGTCCCTATTCTCGGTAAG    120
fso:g9516c      TGGAGAAATTTTAGCCTCATTAGCAAAAATGAAGTGCAGCGCGTCCCCATTCTCGGTAAG    120
fso:g9516d      TGGAGAAATTTTAGCCTCATTAGCAAAAATGAAGTGCAGCGCGTCCCCATTCTCGGTAAG    120
                *********************************************** ************

fso:g9516a      GCCATCGAGGTTGGCGGCAATATTCTACCTGGTAGTAGATCGACAAGATCGCAAGTCGCA    180
fso:g9516b      GCCATCGAGGTTGGCGGCAATATTCTACCTGGTAGTAGATCGACAAGATCGCAAGTCGCA    180
fso:g9516e      GCCATCGAGGTTGGCGGCAATATTCTACCTGGTAGTAGATCGACAAGATCGCAAGTCGCA    180
fso:g9516c      GCCATCGAGGTTGGCGGCAATATTCTACCTGGTAGTAGATCGACAAGATCGCAAGTCGCA    180
fso:g9516d      GCCATCGAGGTTGGCGGCAATATTCTACCTGGTAGTAGATCGACAAGATCGCAAGTCGCA    180
                ************************************************************

fso:g9516a      GCTCATGACGTTGAAAAAGGGAATTCAATACCAGTACTTACAAGACTTTGTACATTTGTG    240
fso:g9516b      GCTCATGACGTTGAAAAAGGGAATTCAATACCAGTACTTACAAGACTTTGTACATTTGTG    240
fso:g9516e      GCTCATGACGTTGAAAAAGGGAATTCAATACCAGTACTTACAAGACTTTGTACATTTGTG    240
fso:g9516c      GCTCATGACGTTGAAAAAGGGAATTCAATACCAGTACTTACAAGACTTTGTACATTTGTG    240
fso:g9516d      GCTCATGACGTTGAAAAAGGGAATTCAATACCAGTACTTACAAGACTTTGTACATTTGTG    240
                ************************************************************

fso:g9516a      AAGGTATCCCGAGGGAACACGTGCGCATACAAGAAGAGTCGTGCCCTTTAA    291
fso:g9516b      AAGGTATCCCGAGGGAACACGTGCGCATACAAGAAGAGTCGTGCCCTTTAA    291
fso:g9516e      AAGGTATCCCGAGGGAACACGTGCGCATACAAGAAGAGTCGTGCCCTTTAA    291
fso:g9516c      AAGGTATCCCGAGGGAACACGTGCGCATACAAGAAGAGTCGTGCCCTTTAA    291
fso:g9516d      AAGGTATCCCGAGGGAACACGTGCGCATACAAGAAGAGTCGTGCCCTTTAA    291
                ***************************************************
```

(B)

```
fso:g9516a、fso:g9516b、fso:g9516e      MLAANHSSWMDTFYLGATVGWRNFSLISKNEVQRVPILGKAIEVGGNILPGSRSTRSQVA    60
fso:g9516c、fso:g9516d                  MLVANHSSWMDTFYLGATVGWRNFSLISKNEVQRVPILGKAIEVGGNILPGSRSTRSQVA    60
                                        **.*********************************************************

fso:g9516a、fso:g9516b、fso:g9516e      AHDVEKGNSIPVLTRLCTFVKVSRGNTCAYKKSRAL    96
fso:g9516c、fso:g9516d                  AHDVEKGNSIPVLTRLCTFVKVSRGNTCAYKKSRAL    96
                                        ************************************
```

(C)

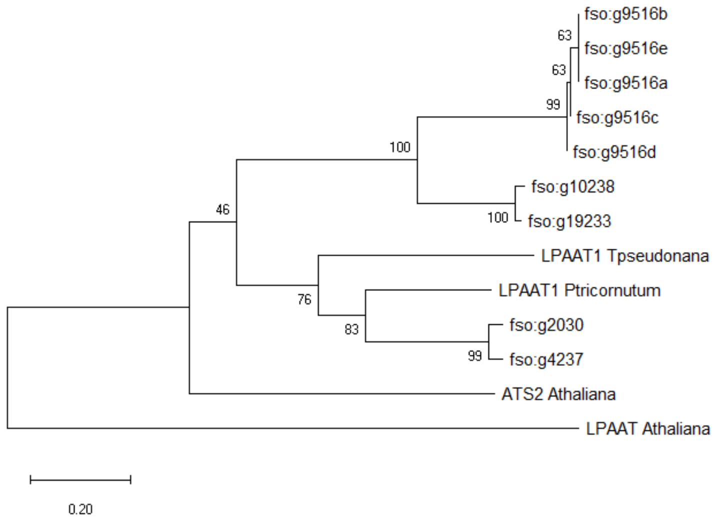

Fig. S15 Sequences of the tandemly repeated *LPAAT* gene (fso:g9516 a~e). Variations of DNA (A) and amino acid (B) sequences of fso:g9516s and their encoding LPAAT. Nucleotide polymorphism at 8th position was supported by the read sequences of the cDNA analyzed by illumine in our previous study (Tanaka et al. 2015). The phylogenetic tree of *LPAAT* genes derived from *Fistuifera solaris*, *Phaeodactylum tricornutum*, *Thalassiosira pseudonana*, and *Arabidopsis thaliana* was constructed with a neighbor joining method using MEGA X.

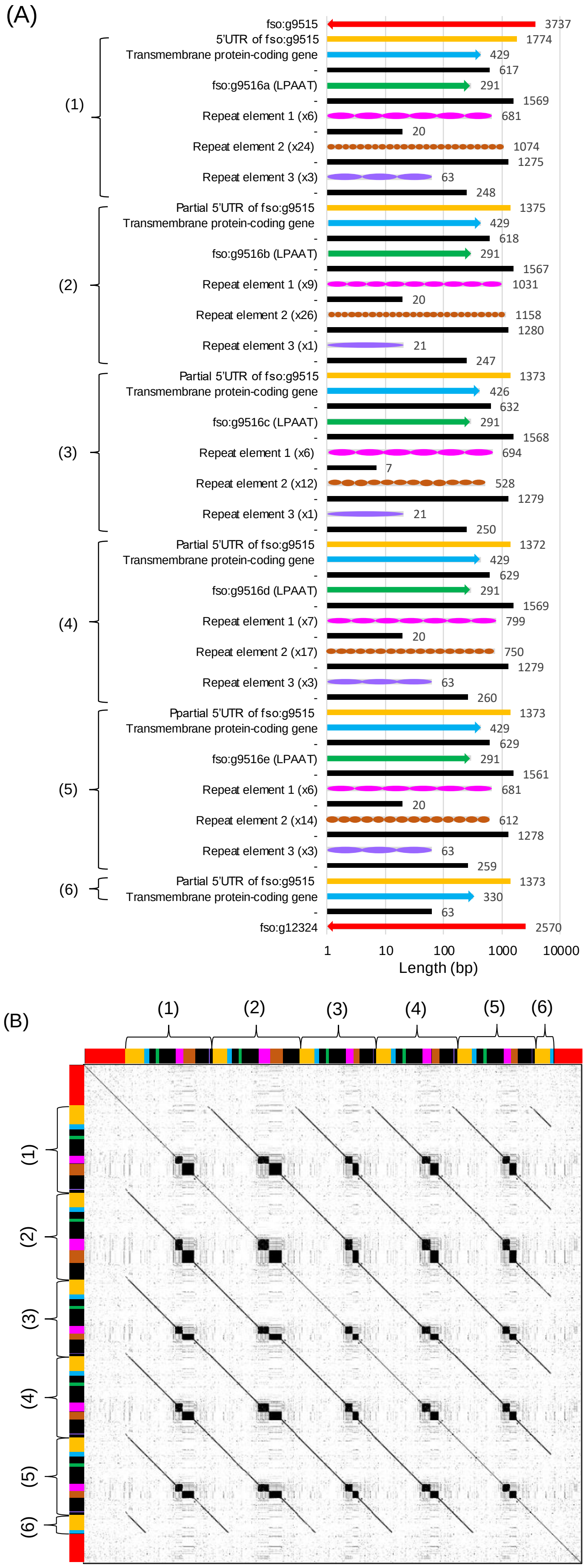

Fig. S16 Length and arrangements of the repeating elements found in the region containing the tandemly repeated *LPAAT* gene (fso:g9516 a~e) (A). Dot plot analysis were performed to visualize the repeated regions (B).

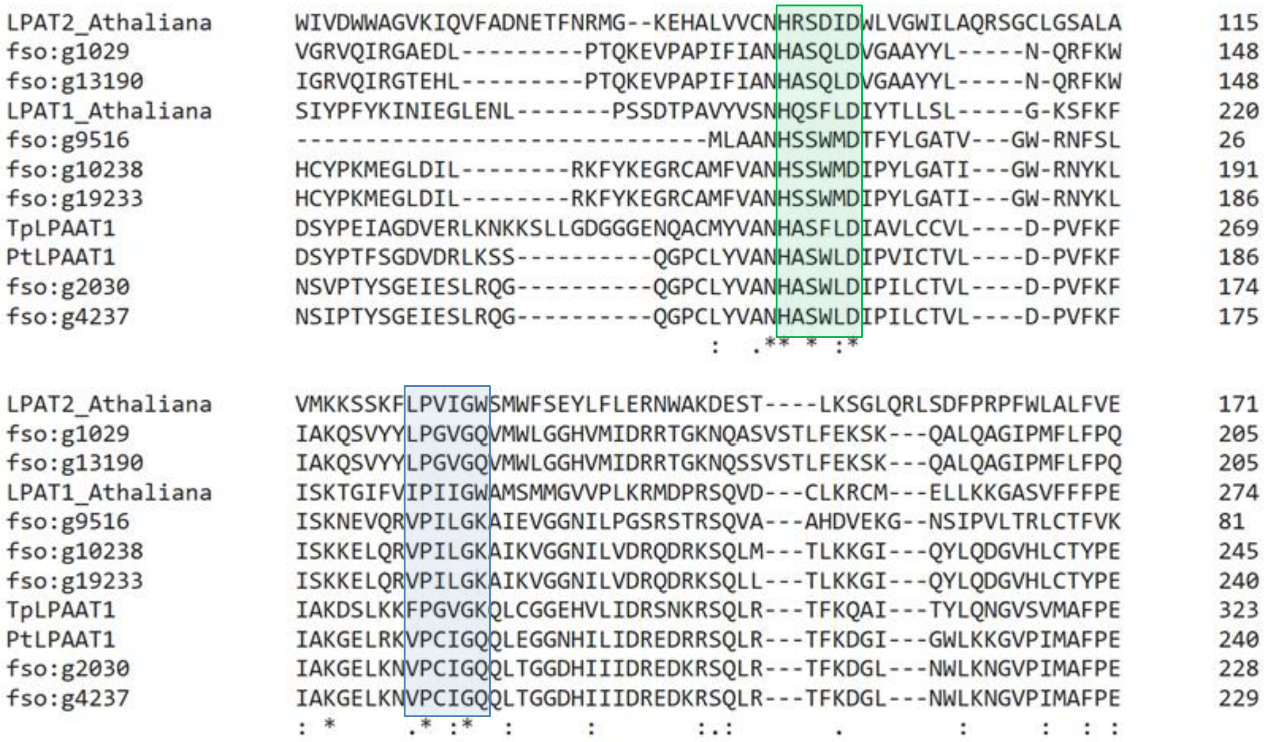

Fig. S17 A partial result of amino acid sequence alignment of LPAATs encoded in the genomes of *Fistulifera solaris*, *Phaeodactylum tricornutum*, *Thalassiosila pseudonana*, and *Arabidopsis thaliana*. Green and blue boxes represent the conserved $HX_4D$ and LPVIGW-like motifs, respectively.

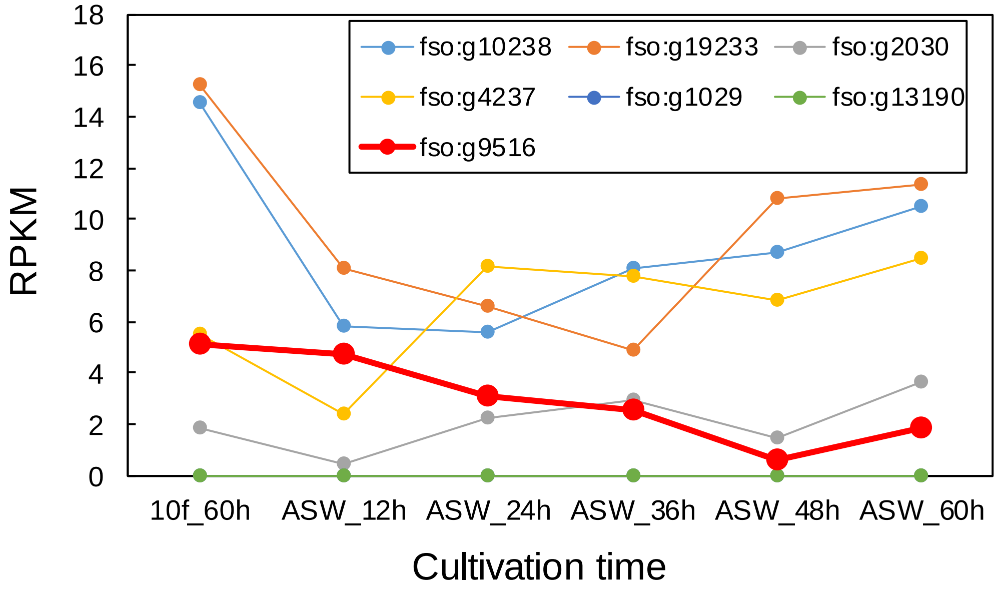

Fig. S18 Changes in reads per kilobase of exon per million mapped reads (RPKM) values of transcriptomes coding putative LPAATs when *Fistulifera solaris* cells were transferred from the nutrient-rich 10f medium to nutrient-free artificial sea water (ASW) (Osada et al. 2017). Prior to transferring the cell from 10f to ASW, the cells were cultured in 10f medium for 60 h. After the cells were transferred to ASW, *F. solaris* immediately accumulated oil in the cells (Osada et al. 2017). Calculation of RPKM values were described previously (Tanaka et al. 2015). which were up-regulated during lipid degradation. The values for the tandemly repeated LPAAT gene (fso:g9516) was shown in red circles, although fso:g9516a~e were not distinguished in the previous transcriptome analysis (Tanaka et al. 2015).

### **Nanopore sequencing of the genome of *Fistulifera pelliculosa* CCMP543**

*Fistulifera pelliculosa* CCMP543 was obtained from The National Center for Marine Algae and Microbiota, U.S.A. We performed MinION sequencing of *F. pelliculosa* genome twice using Ligation Kit 1D2 (SQK-LSK308) and FLO-MIN107. *F. pelliculosa* was cultured for 7 days at 25°C under 130 μmol photons/$m^2$/sec of continuous illumination with 0.8 l/l/min airflow containing 2% $CO_2$. Before cultivation, Hoechest 33342 (Thermo Fisher Scientific, CA, USA) was used to confirm that there is no contamination of culture by bacteria using fluorescence microscopy. For 2 rounds sequencing, the cell cultures reaching approximately 0.7 and 1.1 $\times 10^7$ cells/ml were prepared, respectively. Each culture was observed using fluorescence microscopy to confirm no contamination again, and centrifuged at 8,500*g* for 10 min. Genome extraction was carried out by CTAB method as described in the main text. Agarose gel electrophoresis was performed to confirm no significant fragmentation of the genomic DNA. DNA libraries were prepared using Ligation Kit 1D2 by following manufacture's instruction. We selected ligation kit 1D2 for sequencing *F. pelliculosa* CCMP543 whose genome have not been sequenced yet because this kit tends to provide more precise but shorter sequences than Rapid sequencing kit (SQK-RAD004) used for genome sequencing of *F. solaris* DA0580. Assembly of the MinION reads (>500 bp) by Canu resulted in generation of a total of 573 contigs (1.0 kb~1.9 Mbp) with total length of ~35 Mbp (Table S4). N50 was 176.1 kbp, which was shorter than that of *F. solaris* owing to shorter read lengths.

Homologs of fso:g9516 (LPAAT) was searched in the assembled contigs of *F. pelliculosa*, with local BLAST+, but no homolog was found. By contrast, those of the neighboring fso:g9515 (bromodomain-containing protein) were found from 3 contigs (*i.e.*, contigs 1741, 54267, and 54268 of MinION assembly of *F. pelliculosa* genome). Multiple alignment of another neighboring gene fso:g12324 (aarF domain-containing kinase) using clustalomega allowed us to find the homologs of fso:g12324 in these 3 contigs (Fig. S19). However, as explained in the main text, no homolog of fso:g9516 (*LPAAT* genes) was found in the corresponding sites of the contigs of *F. pelliculosa*.

Table S5 Summary of the nanopore sequencing of the genome of *Fistulifera pelliculosa*

| Organism | Diatom | | |
|---|---|---|---|
| Species | *Fistulifera pelliculosa* | | |
| Size of reference genome | 49.74 Mb | | |
| **Sequencing** | | | |
| Chemistry | Ligation Kit 1D2 (SQK-LSK308) | | |
| Flow cell | FLO-MIN107 | | |
| # of flow cells used | 2 | 1st run | 2nd run |
| # of reads | 5,658,856 | 2,412,559 | 3,246,297 |
| # of pass reads | 284,738 | 61,794 | 222,944 |
| Total base of pass reads | 710,121,755 | 178,281,600 | 531,840,155 |
| Total base of pass reads/flow cell | 3.6E+08 | | |
| Coverage (total base of pass reads/size of reference genome) | 14 | | |
| % of passed reads | 5.0 | | |
| N50 | 4,650 | | |
| Average base of pass reads | 2,494 | | |
| Maximum base of pass reads | 90,906 | | |
| **Assembly** | | | |
| Software | Canu v1.7 + | | |
| # of contig | 573 | | |
| # of contig with homology to reference | 62 | | |
| Total length | 35,793,069 | | |
| Largest contig | 1,938,242 | | |
| N50 | 176,105 | | |
| Reference | This study | | |

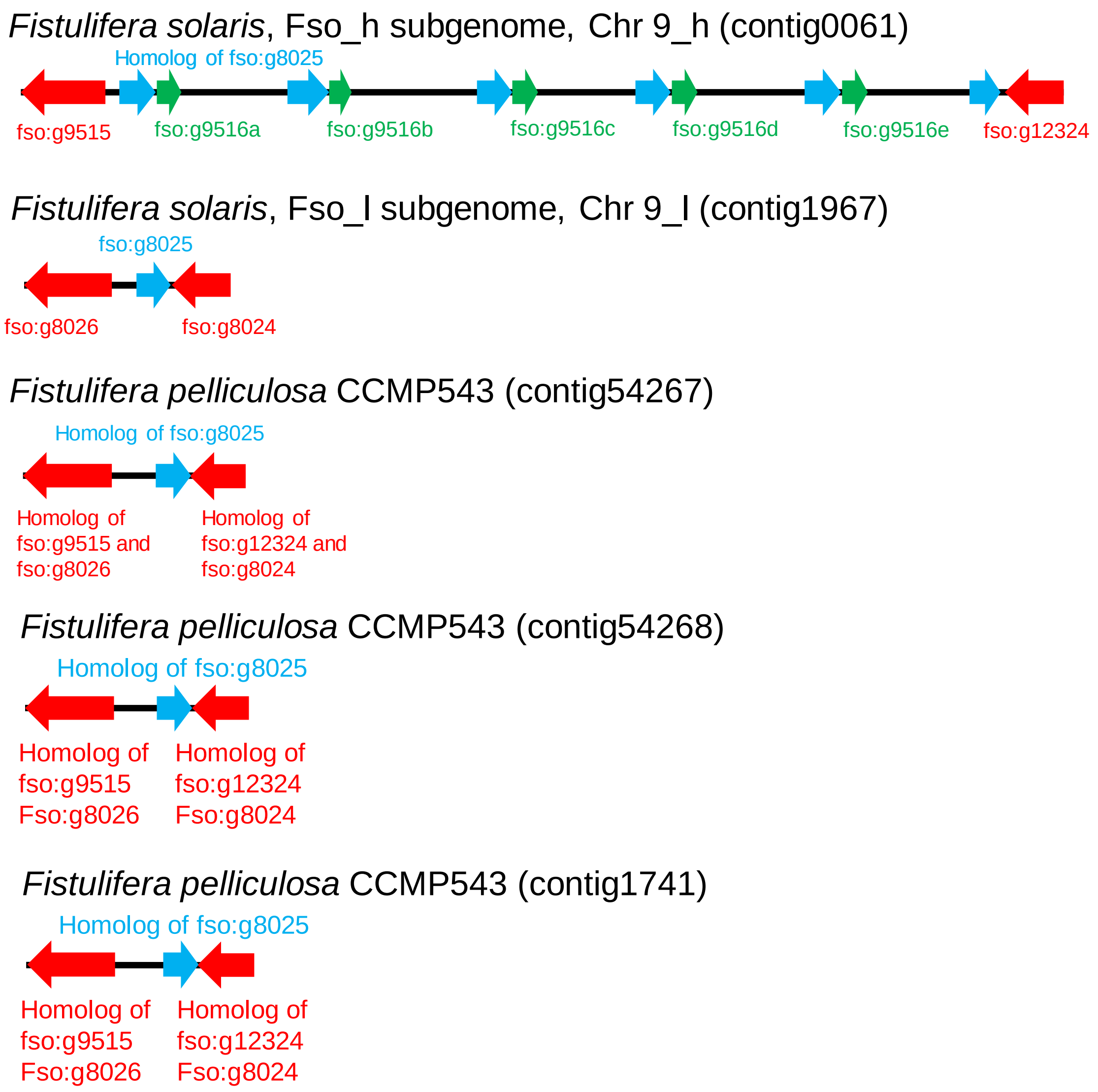

Fig. S19 Arrangements of the genes of the tandemly repeated LPAAT genes and neighboring genes in Chr. 9_h. The tandemly repeated *LPAAT* genes were not found in the homoeologous chromosome Chr. 9_l, and the corresponding 3 contigs of *Fistulifera pelliculosa* CCMP543 which is a closely-related species of *F. solaris*.

(A)

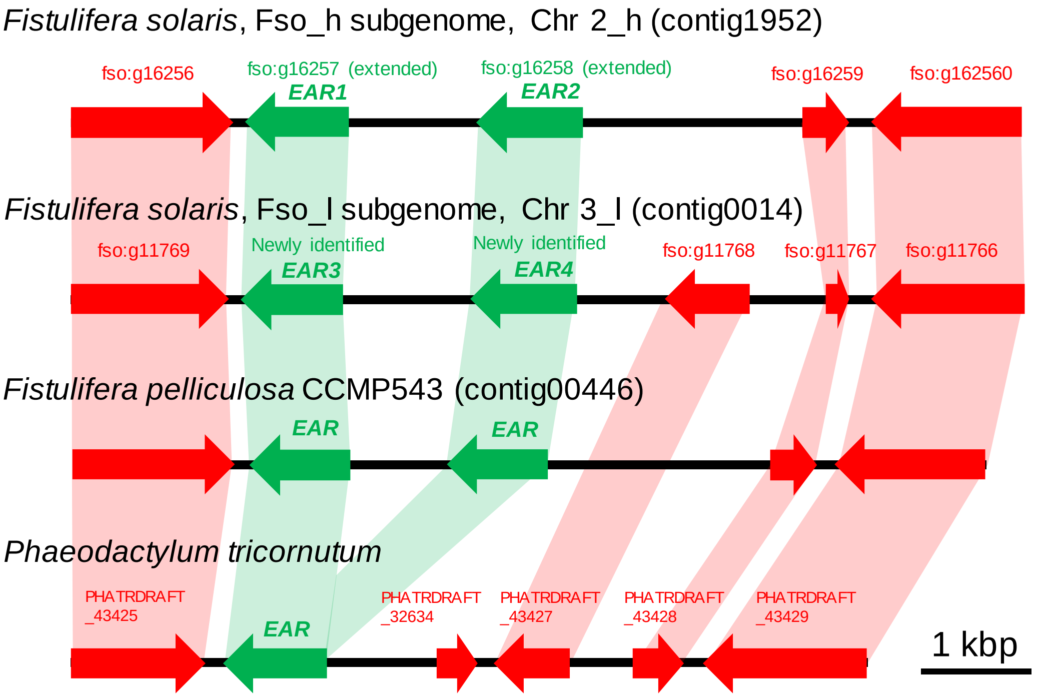

(B)

```
EAR1_Fso_h      MAKTVDLKGKVAFVAGVADSQGYGWAIARALADAGATIVVGTWPPVLKIFQMGLSKGQFD    60
EAR3_Fso_l      MAKTVDLKGKVAFVAGVADSQGYGWAIARALADAGATIVVGTWPPVLKIFQMGLSKGQFD    60
EAR2_Fso_h      MAKTVDLKGKVAFVAGVADSQGYGWAIARALADAGATIVVGTWPPVLKIFQMGLSKGQFD    60
EAR4_Fso_l      MAKTVDLKGKVAFVAGVADSQGYGWAIARALADAGATIVVGTWPPVLKIFQMGLSKGQFD    60
                ************************************************************

EAR1_Fso_h      EDSTLSDGSKMTIEKVYPLDAVFDEPDDVPEDIKNNKRYAGLDGFTISEVAKAVEADYGK    120
EAR3_Fso_l      EDSTLSDGSKMTIEKVYPLDAVFDEPDDVPEDIKNNKRYAGLDGFTISEVAKAVEADYGK    120
EAR2_Fso_h      EDSTLSDGSKMTIEKVYPLDAVFDEPDDVPEDIKNNKRYAGLDGFTISEVAKAVEADYGK    120
EAR4_Fso_l      EDSTLSDGSKMTIEKVYPLDAVFDEPDDVPEDIKNNKRYAGLDGFTISEVAKAVEADYGK    120
                ************************************************************

EAR1_Fso_h      IDILVHSLANGPEVVKPLLETSRKGYLAASSASAYSAVSLLQKFGPIMNEGGSFLSLTYI    180
EAR3_Fso_l      IDILVHSLANGPEVVKPLLETSRKGYLAASSASAYSAVSLLQKFGPIMNEGGSFLSLTYI    180
EAR2_Fso_h      IDILVHSLANGPEVVKPLLETSRKGYLAASSASAYSAVSLLQKFGPIMNEGGSFLSLTYI    180
EAR4_Fso_l      IDILVHSLANGPEVVKPLLETSRKGYLAASSASAYSAVSLLQKFGPIMNEGGSFLSLTYI    180
                ************************************************************

EAR1_Fso_h      ASEKAIPGYGGGMSSAKAQLESDTRTLAFEAGRKYGIRVNTISAGPLKSRAATAIGKSKE    240
EAR3_Fso_l      ASEKAIPGYGGGMSSAKAQLESDTRTLAFEAGRKYGIRVNTISAGPLKSRAATAIGKSKE    240
EAR2_Fso_h      ASEKAIPGYGGGMSSAKAQLESDTRTLAFEAGRKYGIRVNTISAGPLKSRAATAIGKSKE    240
EAR4_Fso_l      ASEKAIPGYGGGMSSAKAQLESDTRTLAFEAGRKYGIRVNTISAGPLKSRAATAIGKSKE    240
                ************************************************************

EAR1_Fso_h      PGSRTFIEKAIDYSKANAPLAQDLYNDDVGNAGLFLLSPMARTITGITMYVDNGLHAMGM    300
EAR3_Fso_l      PGSRTFIEKAIDYSKANAPLAQDLYNDDVGNAGLFLLSPMARTITGITMYVDNGLHAMGM    300
EAR2_Fso_h      PGSRTFIEKAIDYSKANAPLAQDLYNDDVGNAGLFLLSPMARTITGITMYVDNGLHAMGM    300
EAR4_Fso_l      PGSRTFIEKAIDYSKANAPLAQDLYNDDVGNAGLFLLSPMARTITGITMYVDNGLHAMGM    300
                ************************************************************

EAR1_Fso_h      ALDSASMVEETA------------      312
EAR3_Fso_l      ALDSASMMEETV------------      312
EAR2_Fso_h      ALDSASMVEEPVPESAKEPEMVSA      324
EAR4_Fso_l      ALDSASMMEEPVPESAKEPEMVSA      324
                *******:** .
```

(Continued)

(Continued)

(C)

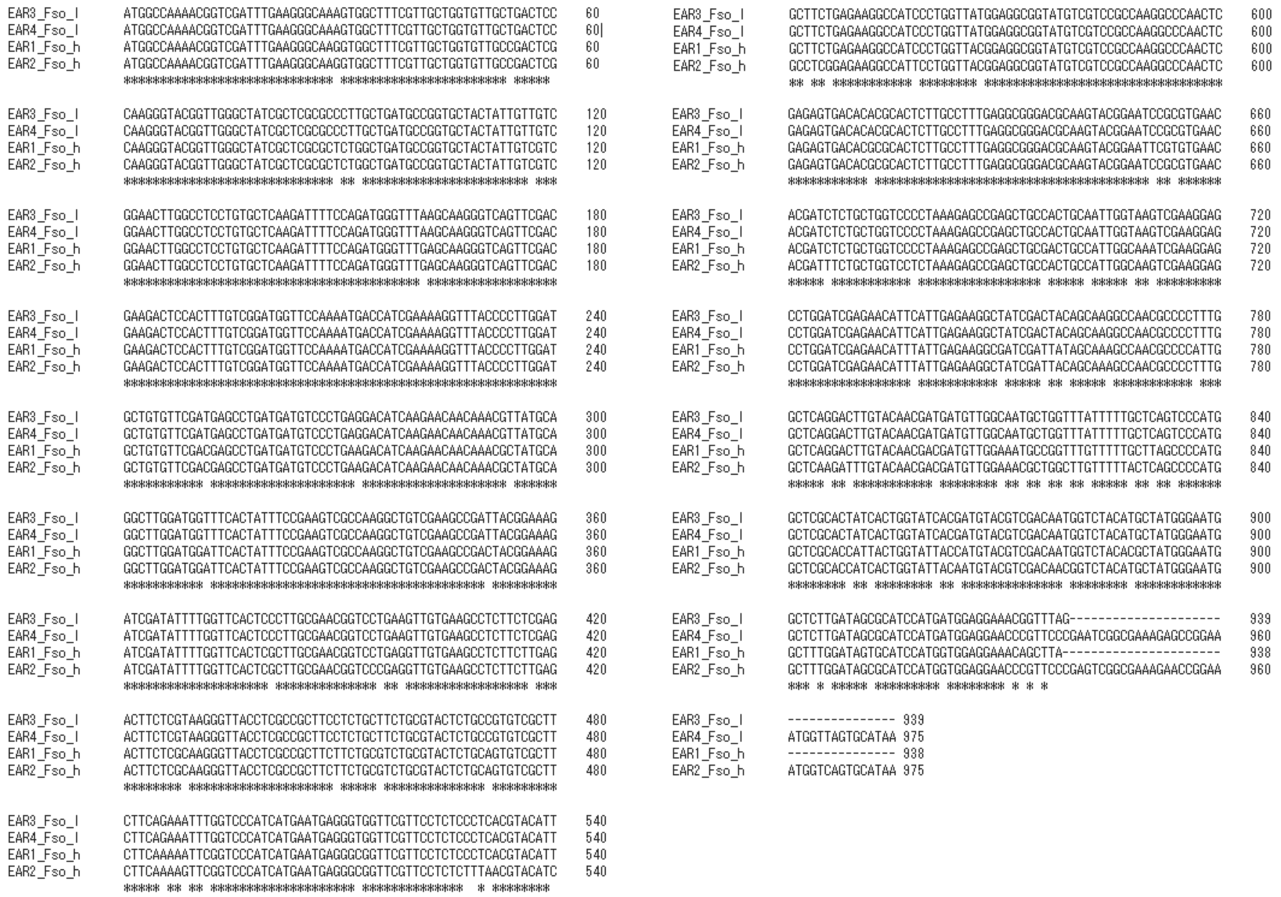

Fig. S20 Tandemly repeated enoyl-ACP reductase genes (*EAR*) found in the genomes of *Fistulifera solaris* and *Fistulifera pelliculosa*. (A) Positional relationships of the gene repeats in the genome of the diatoms. (B) Alignment of the amino acid sequences of the EAR. (C) Alignment of the DNA sequences of the *EAR*. DNA sequence identities between EAR1 and EAR3, and EAR2 and EAR4 are 94.53% and 93.95%, respectively.

### Supplementary data 1: Sequences of the predicted centromeres in the genome of *Fistulifera solaris*

>Centotomere_Chr1_h

GACTCGAGCCCAACTCCAAGAGCTGATCCGGTATACCTTCGGAACTCAATTGGGTGTTCGCGTGGACAGTTTGTTCCCGGCACAGTTCCCTGCAGATAGA
CAAACGCCGGGGGATGTTTTGCATGTGGTGACCTTGAAGATTCTTCAGACACGAGAAACTGACTGACAACCGGGTGCGATGTAGACGGCACCTACCAATA
GCGCAGTCATACTTGTCGGTGGAATTGCTGCCACCGCCAGTTTAATTTAGTATTAAGAAGAATTTGCGTTTTTATTCATCTTGAGCGTGCTATGAATAAGCAA
GTTTTCATGAAAGCGAATATAATAACCGTTGTAAAAGTCTTACTTTCTATTTTTCACGCGCTGCAACAAATTCAGCCGCCACTTCAACCGAAGGTGCTGCTGT
TGTTATGCGAAAGCTTTGACCATTGTTTGTCAGGTTCCTCGAACAGCGTGCAAAAACGTTCAGAACACAACTTTCGTCGCAAGCTAGAAGAAATACGTCGC
AAGAGCGAAAACAACATACCGTGACTCCGTTTGTCACGACCAAGAAAAATACAGGTACGGTACAAGTACCGAGCTTAAAAGGAAACTTACTTGGCGTGCTA
TAGAAAAAGTTAATACCGCTGTTGCAGTTGACTTGTTGCTGCAGAGTGAGGAACAAACAAAAGGTGATGATGATGCGGCTTGAAACAAAACACATCACGCC
ACCGACGACCTCCTTCTGCAAAATTCAGTTCCGACACCAAAACGGATCCAAGACTTTGCTGACCGAGCTCAATTAAGCGGTACATGGTTCCATAAGACGGT
TAATCATTTGGCACCAAAAGACTGCACGAATGTGACGCATCGACAAAGGATTTTTCTTGGGCGCAACATTTTGTGGCAAACAGCTTAGCAGTCGTCTGTGT
GTAAAATTTTGAAGATTAGAATACATTTTTACAAATTGTGGTACAGTGATTCAGAAATCGGGGTTTCTGAAAATAAAAAAGAACGATGAAAAATTCAGTCGAAT
TTCAACATTGCATTCTGGCTCAAGAAATAAGGAATAAAAATACGAATAAATATAGTGAAAGTTCGTTTTACATCGTAATGTCAAAAATCAAATAATGACCTTCCT
TGTTGTTAAAATGCTTTCATCTTCCGCAGCTACTTATCGACATTTTGAAAATACAAGCCGCCTCCAACATATTTCGTCGTTGTCTCAAGCTGTGTTCCTGCCG
ATTAAAAAAATACGAGGCAACACAACGCGGACTGACATCTATTTGTGAGCTTTGAACACTACATCCTGATATATATTTCTAATTTCCTAAACTCAGCATGTAATC
ATGACATTCGACAATCAGAAACTTCGCTGAGGAGCTAGTGCAACTGGAGATTTGCAAGCCAGAGCCCGTTCCCCAATATGTGAAAAAAACGTCTTGGTATT
CGTGTTGTAAACTAATCCGGAACAAAGGCTTTGTGTTTAAGCTGTGGCCCTCCCCAAGACAAAAATACGTGCTTTGAACACCTTGATGAAAATACGTGCTTA
AAGCAAATCGCGGAGCGACATTAAATGAAAGTCATTGTAAAGAAAGGTGGACGAAATACATCAGTTGAAGATATGATTGATGCTTTCAAACATTTATTTATTCA
AAATGATTTATGTTGTTACTATACTATACGGACTATGAAGTCTACCGTCATGGGATCAGAGCCAT

>Centotomere_Chr1_l

CCAACTCTCTCTAGCAATCGAGTTTCAGATCGACATGAAAAAAGCTTTTGCGGAAACCCAACGTCGACACTGTGTTCAGTCAGGCATGTCGTGGCGTGTTT
TTTCCTCTTGCTGTTTCTGTTGTTCTACCTGACGGGGCGGGCGGCCGACGGTCCGACGCTATTTGATAAGTCACGAGGAAGCATGTTTGCGACCAAAGGA
CTTGACGTCGTAAAGAGAAGTTTCGCATTGATTTTAAGATATTCCGATCCGTCCTTCGATACCGTATCATTGGAGTTCGGCACTGATTGGATGCACAGTAGAC
CCAATAGACTTGAACGGGCGACGGCTCCTCCTTGTGATGGACGCTGCGCGCGAAAAAAAGAAAGTGCTTGGAGTTTTGAGGCGCTGCGAAAGAGGAGG
AGCAGCAAGAGATAGCCTCATCTCTTGAAAAGCATTATGATTTTATTTGAAAGCACGCGCACAACACTTGTACTAGCTATCGTAGCTGTTACACGGACTAACC
GCCAATGAGGTTTGGGACACGAAACTCCCAGGGATAGTGGAAGGAAATCAGTTCAAGGACGGATATAACATGGGAAACATTCTTTGCTCTACTCCGATATG
CCCTCCTCTGCCGAGGCAAGATTGGAATGATGTTTATCAGATTTTGTCGGACCTATACACAACACCATCAGCACTCATTACATACCTAGTGAAAATAGCAAGA
ACCCCGCACAAGCGAGTTCACTATAGTCTGCCACAAGGGGGTGGGCTTGAGCCAACTAATATTGGTCCTGTGATTCCTTTCCTGCGAGACGGCTTAATCAA
TTGCGCCAGAGGCTTTACGTACATTCTACCCATGAACTCCCTTATGGGCAAGACGATGGCAGCTGAATATTTGGTTGCGAAATTACTACAAGGGCAAAACGC
ACTGTACGTCAGCGCTTCAGGTGATAAAGAGATTGTCAACACTCTTCAAGCAACTTTGAATACTACAGTGGGTAGCACCATGCTAGCTAAATGCTTAGTAGC
CGCATTGGGGACAACCGGAGTCGCTCGCGACTATGAACTCCCCATCGCTTTTGATTATTGACGAAGCGAACCGCGAGCGAATCCAATAAAGCTTTCATTGA
AGACCTCTTCCTGGAGATTAATCGTGACAGAGGTTTAACTTGTGTGATTCTGACCAATAAAGAAGAGGTAGCCGAAGCCTTGCTAGCGCTCAATGGAGGTA
AAATTCGACCATTTCCTGGCTCCGTTGTTGAACCTTGGAAGCCCCTCGAACGGCCTGAATGGTTGCGGAATGTTTATTGGACTAGTGCGCAACTCCAAGAG
CTGATCCGGTATATGTTCGGAACTCAGTTAGGTGATTGCGTCGACAATTTGTTTTCGCCACAGTTCCCTGCAGATGAGCAAACGCCGGGGGATATCCTGCA
TGTGGTGGACTTGAAGATTCAAAAAAAGAAATTTACTGATAACAGGGTGCTATGTAGACGACATCTACCAATACCAAGGTCTATACTTGTCGGTGTAATTGCT
GCAACCGCCAGTTTGATTTTATCAAGAAGAAAGTAAAGTCCTTATCTATTTGCACTTTATTTATCTCACACGTGCTATGAATAAGCAAGTTTTCATAAAAGCAAA
TGATAACCGTTGTAAAGTCTTACTTTCTATTTTTCACGCGCTGCAACGAACTAAATTGCCACTTCAACCGAAGGTGCTGCTGTTGTTATGCGGAAAGCTTTG
GCCATTGTTGTCAGGTTCCTCCTCGAACAGCATGCAAAAAAACAACGTTCAGAACAAAACTTTCGTCGCAAGAGCGAAAACAACATACCGTGACTCTGTTA
GTCACGAGTCACGACCAAGAAATAATACGTAGTATTATTGGTACGGTACCGAGCTTTTTCGAATCTGACAAGGAAAGGTACAAGGCGTGCGCTGAAAAAGTT
ACCAGCGCTTTTGCAGTTGTTGTTACTGCAGCGTGAGTGAAGCAAACACAACCAAGTGATGATGATGCTGGCTTGAAACAAGACACATCATGCTACCGACT
CATGCATACTTGAGTCCCGACAAAAAAACTGGATCGAAGATTTTGGCTAAGGCGCTCAACTAAGCGGTACTTGGTTCAATAAGACGGTTACTACGTATTCGG
CACCAAAAAGACTGCATGAATGTGACGCATCGACAAAAGGCTTCTTGGGCCCAACATTTTAGTGCCTTATATCTGTGTGTGTAAAACAAAGGAAGTAAGGAA

TAAAGTTTTACAAGCTGGGTACAGTGATTCAGAACCCGAATTCTCTGAACAATAAAAACGAACAATGGAAAATTCAGTCGAATGTCACAATTGCATTCTGACT
CTAAGAAATGAGTGAAGCTCCTTTTTATATGTTTAGGATTGTGTTGTGGAAAAAACCAATTTTAGGAATGCCAAAACGGAATCCCCTCATTGGGAAACGGGCT
CGGGACTGGCCCTCTACAAGCGCACTAAGGCTAGAGAGTTAGAACCAAAGCTTACTCAGAGGAGTGTCAGACTGAAATATATGTAAATGAAAAATGGAAAAT
TCAGTCGAATGTCAAGATTAGATTCTGACGAATAAAAAATTAGGAATAATATTGGTAGTAAAAAAAAATTTCCATTGACGGAATGATGGGACAACAGGCTATTT
CAGCATCGAGACAACTTGCAAGTAAACTTCAGCGGAAGTGGAACAAGCCATACTCCGAGATCTGTGGTTTCATCAGAAACCGCATGAGTCTTCCCCCTGGT
CAGGTCAGTTAGTACGCTTCTGCGGAATCAAAGAGAAGGATGCCAAGACGGACTCCCCCTTATTGGGAAACAGGCTCGGGACTAGCCCTCTACTAGCACT
AGAGCTAGAGTTAGAACCAAAGTCTACTCAGCGAAGTGTCAGACTCAAAAATATACTCCATATGCGAAATAATGAGATGAAAAATAGAAATATCAGTCGAATG
TCAAAATAACATTCTGACGTTAGGAAATTAGAAATAATATTTAAATTTAGTGAATAATTTTTGTTATAATTGAGGCCAACGAAGTCCAGGCCTCTTGGGTGTAAG
AGTGGCATTCCTCGTTGTAAACGCTTTCGTCACCCCAATCAGTAATCGACATTTTAAAAAACCGGCCCAAACGTATCTTCGTCGTTATCTCAAGCTATGTTTT
AAAAAACCTCGCGGACGCATATATATGTGAGATTGAACACTCAATCCGGATAATAATTATTGATATCTTAAATTCAGAATGAAACTTTGTCAAATATATTACAGTC
CGGACATCCGCTGAGAAGATAGTGCAAGCCCAGAGCCCGCATTGCAGTTGTCCTACGCTGTGTTCTTCCCGACGATACAACTACGTGCGTAAATCACCTC
GCGGAAGGAATTATTTGTTTATCAATACGGGCAGTTATTTGAAAAAGTAGATGGGCATGTGTTGCTTTCGAGTTCTCCGAAGGTCAACATAAGACATAATGAC
AACCAAGGTCCCTTTCATATTAAACTCGAATGAGCTGATAGCACAATATGTTAAATTTAAGCGATTAAAGTGTTTTCAGTTTCCACGCTCCTTCATGTAAATGA
ACACCTCACGAACGCACATTACATTGTTTTGATAATGTCAAGGTAGGTACGCCGAGAGAACCTTTAGTGAAGCTGATATTCGCATGGATATAAAAACACTTAA
CCCGTGTAGACCATCGCTTACAAGGAAAAATTCTTTGACGGCACAAGTGCTTTGATGATAAAAAATATTTCAAGAAGCAATTTCATCATTCACTAAAAAAGAA
CCTATGAGCTTAATTTTCATTGCTTTTGCAAAATCGAATTTTACATAAGAGAATATAAATATGATCTGTGTCTTACCTGGTGTCTTTATATGCATCATTCACTAAG
AAGAACCTATGAGCTTATTTTTCATTGCTTTTGCAAAATCAAAACTTACATAAGAGAATATAAATATGATCTGTGTCTAACTTTGTGTCTGTATATGAATGACGGA
AGAATCATT

>Centotomere_Chr3_h

CGTAGCTCTTTGGAGGCAGGACATCCACTCATCTCGACTGTATATATCGTCACTGGCGAGGTGCCAAACTAGTCTTCCACCAAGCGTTTCGACTGTCCAGG
CAAACTTCTCGCCGACGCTAATAACGGTGTGCGTCTGTTTGAGGTCAGTCAAAGCTAAGCGGTGCTTTTTATTTGCACTGTTCTGTTTCCTTGAGTTATTATC
ATCGTAATGATGGAGGATGTTAGACGCCAAGACAAAGTAACGGCGATTCCAGTAAAGTCTGCCGCGCGAGGATACCTCTTGTTTCCACAAATAGCCTGCTT
TCTGCATACACACAGGATTCATTTTGATAATTGTATCACTACTCACCGCCTGTAAATTGTTTTCTTTCCCTTCCAGCCCCAATCGGGCTACTTCGTAGGCTATG
TGCGCGGTAAGCCCCGTAATTTTAAAGATTTTTCGACTGCCTTTCGGTGTAAACTGATACTCTCGCAAGCAAAAGTCTTCTCCAGACCATTTGACTTGATAAC
GTGAAATGGGGATTTCAAGAAAAGATCCAGAGGTACAGCAAAAACTGCTTGTACCTCGGATACATTAATCGTGCAGTTTCCCACATCGAACGAGCCATCAG
CCCATGCAACAACGGGTGTAACACATAAATGATGTAGCGATTCAATTGCTCGCAAACAAGCCAAGGGCGTGACATTCTTGGGCTCGAGACCGACTTCTTCG
TACGCTTCCCTCAATGCCGTTGCAGTGTCATCTGCGTCTTCTTCATCGCGACGCCCTCCTGGAAAACAGCATTCTCCCGGATGTGACCGAAGCGTAATAGA
GCGTTGTGTTATCAGTGTTTGTACTTCATTTTTATCTTCCCACAACAGCACGAGAACACTTGCGCGTTTCAGAGATCGATGATGTGCGCATTCTTTCGGTTCC
AAGCCGCAAGCCTCCAGCCGTTTCCGACACGCCGAAGACATGAAGAGTAGTAGGCGGACCCAGTTTGACCAAAAACAAAGAAGAAACGCTACGAATATTC
GTTCCGCAACTTTTACAAAGCAATGCAGCTTCCGAACAATCGCTACCAGTCAACAAACACTCCGAAAAATAGGCGCTGCTTCTCCGAAAAATTGGAAAACA
GTGCAATCAAAAAAGTTCAAATTATCGAGGTCGTCGTGCGAGACGTACCCGCGTACAGACATCAAATACCTGGTTTTGTACTCCGTACTATGTTTGTCGTCG
CTGAGATGATGACGTCGAGCCTACCGAAGTCACATCTAAAAAAAATCACCTTCCCCTAAAAGCGTAGCATGTATGACGCGGACCCCCCTCGCCATGAACGC
AGACGCGATTTCACTCCCTTACCAGGTAATCTCTTCCATTGACGGAATGATGGGACAGCAAGCTGCAAGGATTCTACAGCGGAAGTCGAACGCCCTCAAA
GATTTGTGAGTTCATCGGAGCGCGCCCTCTACTAGCACTACCCCTACAGCTAGAGCCTACTCAGCGAAGTGTGAAAAAATGAAAACACAAAATGGAGAATT
TAGCCGAATGTCAGAATTACATTCTGATTTTACGAAACTGAAATTTGTTTGGACCTTCCCCGACCTGGTAACCTAAGCTTGTGTCGATAAGTCTAGAAGTACT
GATTACAAGCCGTAATGAGCATCTATTTTACCGGGCGCTCAAAACTGTGAGCTCTAAACTAGTCGAAAAGATGGAATGCTCCTGCATCAAGAAAAAAAAATT
CCGTTGAATGTCAAAAATGCATTCGAACTTTAGGAAATTAGGAATAAATATTTAGGATTGTGTTACATGTGGAAAAAAAAAGAGCTTGTGTCGACAGGCTTGA
ACTGGATTTGCGCCAGACCTCACTATTTGGTGTGTAGGTCGTAACGTTAATCGCGTTACACACTAGAACTGCAACAAAAACAGAACAAACCAAACAAAACAT
CTTGCATCGTGTAAGCGAAACACCATTTCTATTACGTTTTTGGTTCTGATGCGTGAAATCAAAGAAAGTAAAAAAGGGGGTCCTTAAAGACTGACAACAATAC
AGGGAGAATTAAAATTGTGTCGTCAACAAAAGGGGACAATTGTGTCGTCAACAAAAGTGGTAAGCATGGAGATCATAAGATGAAAAGTCAAAGCGTAAAGG
GGACGCAACTGGGGACGCTAGCTGCTACGGAGTATCAGAAATATAGTCGCTAATTATATTAAGTAAAGAATAGAGAGGAAAAGTAGTGCACTTATATGAGGCA
TGGAACTTTCACCTGATAGAGAAAGGCCCACTAACCACCCCGTTTAACCAGAGGCTTATAGGCATACGCAACTACAGCCCAAAGTGCTTGTGTCCTCGCAC
ATTACTTACTCACTAGGAGCAAGCAGAAACGTGAATTTTAAGGCGACCACCATCAAAAAACAAACCTGCTTGGAGGCATCATCCTACCAATTTTCACTTAATG

AATTCATACAACCTCTTTCCTTGTTTCCGTCCGTTCAGTTTACAAGCGATTTCAACGTCTTAGACCAGCTCTCAGAGACAATAAGCCGATATTTTCAATCACTA
TACAGACAAAATTAAAAGATAACTTCATGTAATATGGTTCCAACGAGAAATTTTTTAAACTGTTACCTGCAAGCGGCATCCGATAAAACAAACAATGGTTCGTT
TTTTAAGGGGGTCGATAGGCACCAACCCCTTTGTGAGAATCGCAAAATCGGTGAAAATTACATAAAAAACTAATACTATTTTCGATGGAAGTAAAGAAAGAAA
AATCGATCAACATCTTATCTCTGAACAATCAATGTACGCGAAGTATGTAAGATATGCGATTACCATCGTTTGAATGCAGGAAATAGATCAAATGGAAACTGCAT
CAACTACGATAAGCAATATTGTTCAATGATTATCGAAGATAAGAAACATTATTCAATGATTTTCAGAAAACCAAAAAAAAATACGAGAAAAGAGAAGTACAAACA
ATCGAGACGATAAGCATGTCAACGAAATTGAGGCACTAGTCTAAGCAAGAAAACTGATTCAATTTAGTAGTACTAAGGATTGGAATAGTACTAATGGATAAAAT
GACCGTTCATGGCACTCTTTGATTTGAACACCTTGTGTTTAAAGCAGCAAAGTTAGTGGTAAGCAGGTAAATGCGATAACACATGATTGTAGGCTAATAGGTT
AAACTCAAGGCTGACATTTTATTTTCAATGATATTAAATCTATTTGATTTTTCTTAAATGGTTGCATATCCCGAAGTCGAGGTAACGCTTACGGAAGGTCAAAGA
TATCTGGGCCATATCTAATTTGTTTGCTTCTTCGTAGTTCAAAAGACGAGGTGAATTGGCGAATATCCCAAGAAGTTACAAACTGAATCAATTAAAAAAGTATG
AAATAATGGAAAGAACTTATTGCTTGATCAACCTATCATGTATGAAATCTTAAATGCTTGCTGATTTTATTCTATAAATATTTTGATATTGCCGATAAGAAGTTTTT
GCATGCACATCCAACAATCATTCTTTGTTAATAACAACAAAAACGCGCAAGTTTTGAAAGTAT

>Centotomere_Chr3_l
GCTTTTTTTGTCAAAACTGAGTTGGCGATTTTGCGATGACTTCAAAGGTTTACGGAAAGCTACGGAAATATATTCTTTGGGGACGTAGCCATTCCAAGAACG
TTTCGTTCACTCTTACTTGGGTGCATAAGGCTCTTCCAATGCTTGAGTTGATCAAATACTTTCAAGCAAAAAGTAGTCGAAGCCACACAGCCGTTACTTGTC
AAATTTGTGCACTGTCTTCACTGCCAAGCATGCCGACGTGGCTCTTTGAAGACAGGACATCCATTCCTCTCGACTAAATATATCGTCACTGGCGAGGTGCC
AGACCAGTCTTCCACCAAGCGTTTCGACTGCCCAGGCATACTTCTCCCCGTCGTTTACAACGGCGTGCATCTGTTTGAGGTCAGTCAGAGCTAAGCGGTG
CTTCTTATTTGCAGTGTTCTGTTTCCTTGAGTTATTATCATCGTAATGATGGAGGATATTAGACGACAAGACAAAGAAACGGCGGTTCCAGTAAGGTCTGCCT
CGCGTGGATACCTCTTGTTTCCACAAATAACCTGCTTTCCGCATATGTACAGGATTTATTGTGATATTTGTATCACGACTCGCCTGTAAATTGTTTTCTTTCCC
TTCCATCACTAATTGGGCTACTTCGTAGGCTATGTGCGCGGTAAGGCCCGTGATTCTAAAGATTTCTCGACTGCCTTTCGGTGTAAAATGATATTCTCGCAAG
CAAAAGTCTTCTCCAGACCACTTGACTTGATAACTTGAAATGGGGGGCTTCAAAAATAGATCCAGAGGGACAGCAAAAACTGCTTGTACCTCGCATACATTA
GCCTTGCAGTTTCCCACATCGAACAAGCCATCAGCCCATGCAACAACGGGTGTAACGCATAAGTGATGCAGCGATTCAATGGTCCGCAAACAAGCCATGG
GAGTGACAAAATTGGGCTTGAGACCAACCTCCTCATACGCTTCTCTCAATGCCGTTGCAATGTCATCCGTGTCCCCTTCATCGCGACGCCCACCAGGAAAA
CAGCATTCTCCTGGATGTGACCGTAGCGAAAAGGAGCGTTGTGTTATAAGCGTTTGTATTTCATTTTCAGCTTCCCACAACAGCACCAGAACACTTGCGCG
TTTCAAAGATTGATGATGTGCGCACTCTTTCGTTTCAAGGCCGCAAGCTTCCAGCCGTTTCCGACAAGCCAAAGACATGACGAATAGAAGGACCAAGCTTG
ATAAAAGCAACGCTACGGATTTCGTTCCACAGCTTGTACAAAGTAATACGGACTCCCGATCATAACGAAACATTCAAAGAAAAAAGGTACCGTACCGGTAGT
TGCGTGAGACGTACTCGCGTACAGACATCAAAGTCATTTTTATTACTCCGTATTGTTTACGATGTGGTCCAAAAGACAAACGAAGCCACAATTCAACACCAC
GCTTTCCCTAAGGGCATTGTTCTCTTTTGGAAAGAATAGCAGTGTCGTCCAGATAGGATTGTTAATGCTTAAACTGGGGAAATGCAGTGCGCCAAACACCG
CAACGGACAATTTTCATAGGTCGTTACGAGCTCAGATGGTGTAGGAGGGCTGCACCGCGAGATCCCGTCAGCTCATGACAACTTCGCTTCGAATGTTCTCT
TCTTGCAAAGGCCGAAAACAGACAGATGGACTATATAAAAAATCTAATAGCATCAGCACAGAAAACAAAGCTCTGAAAAGCCCTTCGAAACTTGATATTGAAA
TCCAACTTAAGCAAATAAGGGAATAAAATTATAAAGGATAAGTGAATGTGGACAACAGCTGGCAGGACTGCTCACAGCTTGTCAAGCATTCATTTTTGAGATA
GTGCACGCGGAACTATTACAACGAGACCAAATCTGAAAACGGCGCACCATAAAAAGGGGTCCCCTGGCAGCAAAAATTCAAGTTAAGCCTATAATCAGATC
CTTAACCAAAGGCTAAAAAAGTACTGTTACGATTCGTAGACAATAAATTTTATAAGCGTTATTAAACAGTTCTGCTCATCGAGCTGCCCCGTGCGCGTCGTAG
GACAAGTCTGAAGTAATAAATGGGGTCTCCTGCCGGCAACAAATGTAGGCTAAAAAGACTATAATCAGATCCCCTATCAGCAATGGCTAATATATGTGTCCTC
GACCAGCTTTCAGTCATTTCATCGCAATGCCTACAAATGACGTCGGGATTCTGTTCCTTCAGTCCTTACATCTGCGTTCTCCGATGCCATGCAAAGCAACAT
TATACATTTTCTATTAAAAATACATATTATATTTGCATTTGCGTCACTTTGTTTGACTGATGTACTTGATATCGTGACGGCCGAATGAATCGTGTGCAATAGAAGC
TCCTTTGTTGGAAGCATTATTATTTTTTTAGCAGGACTATTCTATTCTCTTGATATTTCATATTATAAAGGTTAACACCAAGTTGCCCACAGTTGTGAGCATCCTT
CGCTGGTATGAATTGGGAATTGGGCGTTGGTACCTATAAAAATTTCATATCACGTGTGTTTGGGACTGTAAATTTCGCTAGGTATGATTTTCCGCCGGGAAAA
TAGAATCTGAATTTTTTATCGTCGCATCTGTCTTTAGCTGGACTGCAAATCGCAAATGTAATGGTGTTTTGAATCTCCTTCAATGCTTTTGAAACAAGACGGAT
TTATCGTAAACACATAACATAATTTTTTCCGCTGTCTTGCCAAAGCCGAATTTTTTACATCTTAGTAATCATCTCATCCTAGTTCCGATGAAGATAAAAACGAGA
TGAAAGAACGGATCATGTATATTTAAGATTCATTGTTCTTTTTTAATGGAGTTTGAGTTATATGCAGGAAGAGTTCACCACCTTAGTCAATTTCTAGATTTCGAG
AAAATCGCGTTGATTTCGTGTTTCTTCAATTCCGAAAACTTACACACAATGATTTTCGGCACTGAAATCGATTCTTTTCGATGACAGCATAAAAAAAAACCATTG
CGATTTAATTTGTTTTTTTAAGAAGCTTATATTTCCTGAAAACACAAAGTATATTGATGATAAAGATTATTCAAGAAGCAAGGTACTACGGTTTACATTTTTAGAA
AATGACAACGGCATTCATTGGGACTTGGCTAAGTCTTTCTTCGGTTTAGACATTACGCATTCAACTTTATTCGGAAAAGTGAGTCTGTGAAGGATAAGCTCA

GAATATCAAATTAGATAGAAGATAGGGTAAAGCAATACATATTGCTAACAAGCTTACAAATGATAAAGTCATTAAAACGTCTTCGATTTATTATTAGTAAAGGGAG
AGCAGCAAGAGTAGGCGAGGGGTCGCCGTACAATGACAAATATGGCTGTATAAAATTTTTTTCTCCACAAAGGGTGCATAAACCTGTGTCGATAATTTCCAT
GCATTTCACGCATTTCGTTTTTGTAGTATTAAATAGTCTCAATTATTTACTTAATTTCTTCTTCACAAGTTCGTAAGGCTCCCAGCTTCGAAATGGTAAATGGTG
AGGTGATATGCGACAAAACTGTTCTTCCTGGATTATGCTGATGAAAATTGGAAGGAATAAATACAACTGAATTAATGTTCCAGAATGGATATTTTACCTGGCTC
GTCGTGATTAGGTGGATTGTTTCTGCCTCAACCATTTTGTCAACACGCATTTCACAGTGATCCTTACTCCATTTAACAACGGAATGGTAATCTGCAATACTCC
ATTTTTGGTTGCTTTGGTAACATTTAAAGGAATAACCCTGGGTTTATCTTGAAATATAAAAGCTTTGGTTAAGTACACTTTCTGCAAATGATTCAATCGCGGCA
AAGCCTGGATTGATGAAAAAGTGATACATCAAATCATTAAACAATTAACTTGAGTTCTGTGTAACTCATTTAAGGATTCCAATGAGCACTTTGAAATGTGTAATT
TCAT

>Centotomere_Chr10_h
AACGAAAGCCGCAAAAAGTCTATATAACAGGCGTAGCTTAGCTAAATTTTGACACTTCGCACTTAATTTTAATAAAAAAATTGGTTCAAACAAAATCTTTTGATC
TTGTCATGTAAAAGAGGCGAAATATCCACAAGTCGGATAAAGGAGAGATAGCGCTGCAATAACTCCCAGCATACGATTTGCCCGAGAGGGGGATGACGAAA
GTGTTCGGCTTCAGCCTTTCTAACCACAGATAATCCTATTATTAACCCTAATTTCGAAAAAAAAATGAATCCTGTAGATGGTAAAAACTGACTTCAAAGAGCAG
ATCATAATAACAGCATCAAAGTCCTCTCGAATGTCTAATTCAACCCCTATTCGTTCTCGATGTGTTATCCCCAATGACGACATCAACACCATTTAGCGAGCGAA
CAAAAATAAGTCGAGACACGTTGAAAAATATCTTATCAATCGAAACAACGTGATCTATCTCATAGTTAAGTGTAGAAAGGAAATTTATTGGGAAGTTGTCAACA
CGTCAACGATGTCCCCTTCGCTTCCATCAGGCACCAGCTCAAAGCATGAAAAACTCCGTCTGTATTGTGTTTTGTGTCCTTGCGTCGTCACGGATGCTTTC
CCAATCATTGGTTTCGTCAAGTCGACCAATACTTTCGCTATTCGAACATCTAAGTTTATGAACACGTAGGTTACGAAACTCTGTATTTTCTTGGTAAGAAAAAC
AGTAAGCGCGTTGCCTACCCGATCAACCATTAAGTGAGCGATCAAACATCAGTAAAGAACCTTAAAAAAATATCAATTTAGTCGAAACAACGTGATCTCTATTT
TGAGTAAAATGGTGAAAGGTTGCGAACACGTTCAACGAAGCACAAAGCATATCAAATTCCGTATCTTCTTTCTTTCAGTCCCCACCTGAAATCAAAATATATC
ATCGCATTTAACAACCTACAAATTAGGTGAAAGCGAAGACCTTGGACGTTACATCTCTTTCGTTAAAAATTCGGGGAAACTTGGCCGTCCAATTCATTGCGA
CGTGAGAAACGAGAGAAAATTTTATTTTGCTAATACAACTGCTCAATGGGATGGAAGTGCTTTCACAATGTTGTTACCAAGGATAACTACCTTCTGATGCATA
GAGTGATAAGGAACTATGAGAATGCTATATCGTGGATACTTTTCTACCCGAAAAGATTATCTGTTTGCATTTTGAATTCACTTGTCTCCATAGCAAGCAAAGTC
TTAACATCACTAGTCTTTATGAGACATTATCAACATTCTTGCTTTAGAGATTATGATAATTGATCTCACTGGACCAGCTTCTAGGATGAAACGATAATGCTGCAT
AGTCTAGCTTAAATAACGAAATGATAGAAGAATTTACTTTACAACCTTTATAAGAATAATGACCTAGATCACCGAAAACATAGGTTTTTGTGCAGAAAGGATATA
AGCGGTGTTTTCAGCAATTAATATCTCGGTGATACAAGCACGTTGGCATGGCGAAGATTATCTATTTAAAATATTGGGCTCTGAATCACCCACAAAATATGTTT
TTGTGCTCGAAGGAAATATGCGCTGTTTGCAGTTAATGATATCTCGGTTATTCAACCACGTTCGCATGTTTTATAAAATTATCGCACTGGATCAGAACTAAATAT
GGTTATGTTCTCAGAGAATGTAAGTAATGTTTTGAGCAATTGATATCTCGGTATGGTAAAAATGTTGGCCTATCTTTTTACATTATCTCAACCATATTAAATTCAC
CAATATGATTATATCCCGCTCATTTATATGAGGCTAGTCATTTTGATTTAATTGAGCCAGGAAAGATGTCCAGTATTGTTGCCAGTGCGCCATACCCAAGTTGC
TTGCATAAGGACGGTTCAATGATTAGTGAATAGTCACAATGAAAGCATATGCTACTAGCATACATATGATCATAACAGCCCACACTCGATGCTTTAACACACCG
CGCTATAATGATACAGAATGAAATCGACGTCAACATGAATAACTAAAAACGGCAAAGATCTTCGCAGCCCCAGCCAGCGTTGCAGTCACAGCAGATATGAAA
AAGCATGTAATTTTTCTGCAACGCCATCTTTCACCTACGCGACGAAGCCTGACGCAAACTTTCTGAAGTAGAAGCATTATCGACTTAAAACACCGCCAGTAC
GACCTCACGGCGGCCACATTAATACCTAAAGACAAATCTCGCCAACATCAACCAATCCAAGCGTTCCTCGCGACGGCCATGCCAGATTGTACTGGTAGAGC
ATGGGTTCTCACCGTTAAGATTGCAAAGGAAGCCCACGAACAGGACCGGTTCGATAAGGATGAACAGGACCTTTAGACCTAATACCACGCTTTTCCAATGA
ATCTCGGACGCATCTACGGTGCAATATACACCTATCAACCTTGAGTTGGTACAACGTAGTTAACACTTTCGGGTCTCTCACATCAAGCTTTAGAGTCGGTATA
GTTAATAACTATATAAATCCTTGTAATATAGTAGTAATTATGATTAAATTGAACACAACGTCGAGTTTCTCTGGGA

>Centotomere_Chr10_l
AACTAGAGCCGCAACAGTTAATCCAGTGAAAGCTTCATTTAGGCCAAGTTCTGGCGCCTAGCACACAATAAAATAATAAAAATGCAACAGAAAGAGTTCAAA
CATATATCTTTCAATCTTGCCGTGTAAAAAATGCGAAATATTCACCGAAACTAGGTAATGACTATGCCTTGCGTCCGATTAGATCAGTGCAAATCGAATAAAGG
AGATGAAAACGCGCAACCTCCCACGAGTAATTTGCTAGGAGAAGTGAGTGACGAAAGCGTCCGCCTTCAGCTTTTTTATAATTATGAGTCACTAAAAAATTT
GTGGTTTCATCAGAACTTACATAAGTCTTTCCCTGGTCCGGTCAATTAGTATGCTTCTGCGAAATCCAAGTGAAGGAACGCCAAGACGGTCTCCCCCTTATT
GGGAAACGGGCTCTGGACTGGCCCTCTACTAACACTAGAGCTGGAGCTAGAACCAATGCCTACTCAGCGAAGTGCAAATATATGTGGAAAAGTAAAACAAA
AATGGAAATTCAGTCGAATGCCAATTTATAACATACCGACGTTAGGAAATTAGGAATGAATATTAATAAAAAAGCTAAGATTTAAGCTTAAGCTCTAGTGCTTTC
ACGACAAAATTGTATTTTAATCGTAATCCGAAATTAACTCAAAGCGAACCAAAGCATCCGAGAAAATAAACTTAATAAGATTCAAGGGAAACGAAGCGCATGA

CATTAAAGATCATAACAAGATCACGAGTCTTCCACCGATTTTTAACTCAGCCCCGATGTATTCTGACAGGTGTCATCACCCACCCGTTATACAATTTCGGGTT
AGCGTTGAACCATTAGAAATGCTTTGTTTAAATGATTACAGCCTGAATATCTCCACCGAAACAAAATATGCCTACAAGCAAACGCAACCATTTCAAACGACAC
AAGAAACAAAAGCAAATTGCATGAAAATATTGTTCCAGGAATTTGGAAAAGTTAGCTAATGACATTTTTAGATGTCTTGTTGTAAAATTGAATAAGTTCCTTAAT
TTGTAGACGTTATGATAAGAGAGCTCTAAGAAGAGAAGCCACAAGTAATCTTTTAAAAGAGTGATATTTCTTCATAGAATATAACTAATATTAAGCTTGTTATCAC
GGATTCTCATATCATCATTTTAGGGGGGGGGGGGATTTCCAAACGTTGAAGGTTGATGTTTTACATTTTTTTAAAAGAGATTTGATAAGGATCGATATGTGATCA
TATTCATGATTTGAACGCGGCGGGATGGAAAAGAGATGGAATCATTTTGAACCATGCTGGTTGTCTTGACGATTAAAAAAAAATTAAACAAGCACAACGTTAA
GTTTGATGAACAGAGTTATAAATAATTTTTTCGTTGAGTCTGGAGAGTTCTTTAAACTTTGAGTAGCCTCAGGAGAGATTACAACATGATATTTTTATGACTATT
CAACCAAGCCTTTAAATGTTATGTCTAACGAAGAAAAAAATTTCTGGTCAGCGGAAAGGTCTTCTTCGCATTCCGTACATACAAGTGTATTATATTAACATTGGA
GAAAAAACGCAACAATACATGATAACGTGTGAGAATAAAACATAGATGTGTTTTGTTTTTAAAATTCTAGTGAAGCAAAATTCTGATTTTATGTATTCGTTAATTT
TCGGAAAGCTATTGAAAATAACGGAAAAACCTGGCATCTAAAGTTTTTTGTTTTTTATACAATTCGTTAATAGACAAATAAGAGCCCTCATCAACTGCTTGGAT
ACGTTGCATTTGAGTCAACGCTTTTGCAAGTTCAGATATAGTTTTCTGAATAATAAGAGCCAGCTTTTCTTTTCAATTTTCAAATCGAGGGACTCAGTCTAACG
CATGATGGATTGATCTGGTAGGTGCTCCGAGCCGAAGGAAAGATATATGTATTGAGCTGTTATCGCAATGGTCAATACCGCCATTGCGAATACAGCACTTTG
GATTTCGAATATCCTTTTACGTATGCGTATGTGCTTGGTCGATTGATCATTCTTCATTTTTTAAAATACCTTACTATTTATATCTTTAACACATAT

>Centotomere_Chr11_h
ATAGTCAACTTGGTTACTTTTGACCGCTTGAACACGTAGATCAACACTCGCGTCGGCAAGACGCAGAAGCGCATATCTCGGGAAAAGAGCTGAAATTTGTC
TTTTTCAAAGACTTAATTAAAAGGATAAGAGAAGTATAACACGATTTCACTAATATTTATTCGGAATGTTCCTAGAGTCAGAATGCAATTTTGACATTCGACTGA
AATTTTCCTTTCTTCCAAAAAAAACTTGATTTCTAAATCACTGTACCAAAGTTAGTAGTAAAGCGACAAACCTGCTCCGAGAGCTCGCTCCCGAATCATCAAT
TTTAACTGCGCTCTAGTTTGATCCTTCAAATGCTCCAGCGGGCGTTCAACAGAAAATTTTGGGCGTTACAGACTTACCAAATGAGCTCCAACAACTTCGTAT
TTGGACCAAGTTATCTTCTGCTTTATCAGTTACATTCTTCTATAATGGTACGCTTACCTCTTGGACCACTTTAAGCAGTGCATTCGCTACTGCTTCACCACACA
TATTCGACGCTAGATTGACTGTCACAACACGTCCGTCCGTGAAAAAGAAACTTCTGCAAATTTTACCTCGTAGATAGACACTCCTCTTAGTTTTCGCAAACAT
TCGATTGCATTACGCCGCTTCTTTACTTATGATCTTCAATCGACCTGATGCAGTTCTTACACCTTATTTCGTCGTGGCCTAAAGTGCCGCTCCCTTATAGGCA
CGATCTTTCAATCCGATTGCTTCTCTGAACATTGGAAATAAACAAAAGAATCCAGCACCAAAAATTAAACAGACCCATCTAGCTACTCCACTTTTGAACAGAC
TAAATGTGGTAAGTGCTTCATCAAGAGGCTTCTCAGAACACCATAACACCTCACGGGTAGTATCTATGAAGTTATAGAAGAAGAAGAAGGTAAGCTGAGTAA
GAGGATTTCGCTTTTTGAATCAATCCTGACATCATTTCCGATCGGGAGTCGCTAGCGTATCTGAAACAAAAATTTTTCACTACGTACTCAACGGACTCACTAC
TTGACTGCAGTAAGAATCGCTAGTGCGATTATGTAATCTGGACAGTGAAAAGCAGTAAAGAAGTTTGTGATAATTAATCCACAGTGCGCGATGTAAGCAACAA
TTTCATCGGAAGATTATCGTATTGAATAAGTTGAAAAGTTTTTTTAAAATAACCCTGCAACTAAAGCAATAAAAATTTGGCGTATTCATGCAGCTGGATACTTA
CAC

>Centotomere_Chr11_I
ATAATGAACTTGGTTAACAAATAATTGAAACTAATATTTATTCCTAATTTCCTAACGTCAGAATGTAATCTTGACATTCGACTCAATTCTTTTCATTAAGCACGTAT
TACACTCGCAAAAGCAAGACGCAAAAGCGCGTATTTAGGAACGGAGCTGAAATATCTTGTCCGAGCTTGAAAAGCCTGAAAGGTCGCTTCTGAATCAATAA
AACCGCGATATGGTTGAACCGTCTCTTATTTGTGCGTGGGCAGTGCTAGACCCTCAACAGAGAGAATTTTAGCACCTTAGACTTGGATCCATCTAGTTAACT
AAGCTAAGCGATTGCAGAATTGTGATCACCGCCGACCACTCCCTAGATTTAATAAAAGCTTTTCGTTCCTGAGGTCGATCTGAACCCGTCCGTTACAGCAAC
TTATAGCTGCCTCACTAGGATACGGATTCAGTATTGCACTTTATGAGCATAGAAGCAAGAGCTTACTCAGCCCGGCGCCAGCTTAAAATGAAAATGGACAAC
GAAAAATTCAGTCCGAAAGAAGGAAATTTGGATATAAAATTATTAAGTCTTTCACCTAAAAATATATCAGCTTCATGGTACCCGATGCAAAAAATATCAAACCAA
ATTTGCTGAGGCGCCTAATCTACCTTTCAGAAAGCATAAACAACAAACTTGAAAACTGAAAGTAAAATACTTTCAATGAAGAGAGCATGGCTTCGTGTTGAAA
TTAGCAGAGCCGTTCGCAAAGGCCAAACCCGTTTCTAAGCAAAAGAATTGCTTGGCCTTTTTACAGATCTGGACTTGATTATCAGGTTCCCCGGAAGTTATA
AATGAGGAGATCTTGAATCAAGTGTTCTGTTTATGGCGTCTGAAAGAGGATCGAATGTCGCCCATGGGTATGGCTGTCTTCATGTTTGTTGTTTAGGACCCA
GGCCTTCAGAGAATCCAAAGGTTGTCAAAGGCGTTGGGTAAAGTAATAAGCGAGAGGATATTAAGCGTTTTAACCGGTAAGATTTTTGAAGTTTACTTTTCTT
TTTTGGTTGCTTTTTGCTTCCCACAATAAAATGAATATAGGCAAATAATCCTTATATCTGTGTTATGAGAGAACAATTTCATTTTAGACTGCACACAAGATTTGG
TTTACCCTATAAATTTTTGAAAATGTTCTATCATATTGCTCAAGTAATTAAACAGCCTAATTGAAATAATGATGATTTTTGTGATCAAACTATGCATTTATTTATGAC
AAAAACTTATGCTTGACAGTGATTTGAATAAAATCTGATAGCAAGCAACAAGCTTAAGGGATCAATACTTTTGCTTGATCAAAAAGTCGAAAACATAGAAAAG
GCACTGTATACTAGGTTACTGTTTTCTTATGCTCGATTTATTAATAACAGCAGCTTGTACACCTCTATTTGATCGCGGATTGATTTTGATTGTGCTAAAGCGTTT

TTAATTGCCTTGTTCTACGTTTTAATCGGCTTAATACGTGAAGCTTGAATTTCTGTTCAATAAGGCTATAGCCCGCTATTATTCCTGATTATTTTTTCAGTTGCAA
ATGCTTTTTTTGGTTTTGATTTTGTAGACTTGGTTGAATATACCACTGTTGTTTTGACAACGCAACGTTCATTTTCATATTTTTTCCTTTTTTGGGAAACAAAGA
TTCATTTTTGGGAAAGACAACATCTATTAGAGAACTTCTGCAAATGAGCTTATAATGCTCTATTGACCTTGGTTTTGGTTTCCAAATGATTGCTTTAGTATTAAC
GCAAAAGGATTTTGTCGTTTGAAAATAAAATGGACAACGTTATAATAAGTTTGAATTTAAAAATGGAA

>Centotomere_Chr13_h
ACTGCCTAGATAGGAAAGTGCACTGAAGCAACGAAGCGCGGTCATAATCGTGTTTTTGTACATGCGAATGACGATGGGGAAGGCACAAGTTAAAGATGCGA
AACACCCTTAAGCTAGAAATTTTCTGCTACAACAGCCATAGTATGCCTTAGCATAGGTGGTCGCTTGTCGCATGAGATGGCCATGAAGAAGGTGATAGGCAT
TCAATCGTACGAGCTGCTAGTGACTAGTGAATGAAACTCGTTAGCTAGATCTAGGTTCTATAAGTCGGCACCATAATTCTCGAATCTTGCTTGGCATTCGATC
AAACAGCTGGAGCAACTGACTTAAACTTTAACCCTTTCAGTACCAAAAATTTTTAAGAGTAATATTTGACTCTTTGCTAATAAACCGTTGATTTTCTTGTCGATA
AACCAATTTATCACGAAACAAATCTCACAAATCCAAACAATAACTATAGTTGAAGAAGAAATAATAATTATATTTCTTTGTTTTGGCTTAGGATAAATCAAAAATG
GCAACATCTAGATTGCAAAGAGTAAATATTTAGTCTAGTGTTCGACACCGTATTAATACCAAAAACAGAAAACCGTACTATACTGTACTAAACCGGTTTTTGGT
TTTTATGATTTTGAAAAAAACGGTTTTTTTCGTTAGACGTCTGTACGCCGTAGACCCTTAAGTTTCCGGTGGATATCTTTTTTAGCCTAGCACGCCTTTTTCCA
AGCAATGGACTATCAGAGCGCAGCAGATCAATAACCAATCACCAGTCTTGATTAAATGTTGAGTTTTATATTCCTTGCACAGTTTCGTTACGCCAGTAAATAGA
CATGAATTTTGAGAGAACTTAATGTTTTATTTTCGCTAAGCTTCAAGTAAGCCAAGAATGTTTTAACCTGGTCAATCAATTGATCACATTGGAAATAATCTCATT
GCATAAAAGATCAAATTGGGAAGCTTTTCTTCTTGAAAAGCTTCCTGTAGCTGTAGCTACAGGGACGTAATGACGGTGTTTTGAATAATACGGAATAAAACCA
TAAAATACGATAAAACCGGTTTAATACGGTTTAAAACCGTTTTGGTTTTTTCAATTTCAATAAAACGGTATTTTTCGAACACTAATTTAGTCTACTCTTAGGCTTT
TTGGTCAAAATATCGGGGCTGTAGATGTCCGTTGAAATAAT

>Centotomere_Chr13_l
TTGTTTCTTCCATGTTCCATGGTCGTGGTTTTTGCGAGGACCGAGCGGTGGTGCCTTTTACAGACTAGACTAAGGTTAGTGGATTTACAAAAGGGCTCATTT
TTTGTTTTTACTTTGGTACCAGGTACCGGTACCGGTACCCGTCGCGTGGAAAGCACCACGATCTCCCATAGGCCTCATTGAAACCCAAGAAACAATACCCT
GTAATGGCAAATTGTCGCATAAAATTGTAAACTCAATTCAATGACAGGGGGTAAGGGATTCCGTCGCGGCTTCTTGCTTGAAAAGCCGAAAAAAGCGATAAA
GAGAGCATCTCCCAAGAACACTAACGTATCATCAGCACTTTTAGATATAGAAAGTCCTTCGCGTTCTGTTTTCGAGGTGACTGCTGTTGACCAAGGAACTGA
CATTTTCGCAGCTGAGCCTGAGCCTGAGCGGCCTTGGAAAAGGATGAGTCGTCGTCGTTATTTACCGCGGTAAAGACAACTTCCCGATCACCTCCTTTCGT
CAACAGGTTAGAGGTTCCCGAAGAACAGCTTATAACCGAAAACGCGACAACAACAATTTTGAAGACCTCGGTTGCAAATGAAGCAGTAACCCTTGTGATTC
CCGAACCAAGGCCAAAATGGCTACTTGCAGAAGAGCCCTCTCGAAATGACGACGACTCTCACGTGCCAAAAAGCGAACTTACAATTCAGGAATTTCTAATG
GAGGATGACAACGACATAAACGAAGATCTTGCAAAAACCCTTCTTCGGATCAAACGAAGTAATGACTGGCGGAAAGTCGGATCGCATTTCGTCCAGGCGC
ATTTGCACACTGCGAAAAAGCGAAATAGCGTTTGGAATCTACTTTTCGATGGTGACTTTACTCTACCGAAATCTCGTCTCGCCCTGTGCTTATTATTGCATCA
TGGACTCGATCCACTAACAAAAGGCTTGGAAGATACAGGAAGCAAATCAGCGCGTTGTCGTGTGATGAAATATATTGCTCTAATTGAGTATCATCTGGATCAA
GAGGGTGTTGCTTCTGATCTCGTTGAAATTTTGCTACCTAAAATTGCTTATGTGGTAAAGTCAGAAAAGAAAAGGACCTATCTAGCACAACAGGCTATGGAG
GTTGCCATTAAACTGATATCTGCAGCAGCCATTATTGGCACAAGCGATACCGATATTGCCACGTTGCTTCACAATTACGTAGCAACTGTTGACGAACTTCTCT
GCACGCAAATGGCTTGGTTCGGCGATGATGTAAACTGGTTCACTGCTGAAGGCGCACGACAATGGTGCAAGTTCGTTGTTTTAACATCCTGGCAGGATGT
GAACGAAACTCATAGAGGCCGGGGAAAGGCAATAGATTGGTGTAGAGCGCTGCAGGGGACTAGCTCACACAACCTGAAGGTTTCAAGCTTTTCTGCGACT
CTTCTTTCACCCCCGAATTTCCCCTTAATAGCTTCATTACCTCACTCCGGGTATCGCTCGAAGCAAATGACCTTAAATATACAAGTTTCTTTAATGTCTTCCTA
GCCTGGTTAGGTCGCAAAGCGAGCCGGGTTCAGATCGACCTCAACGACAAGCGAGATTGTATTGAGGATCTTTTGCTCGAGCTCTTGAAGCAGACAGCTC
AACCACTCTCACTTGTTGCCGCTCTGTAAGTGATTTGACAATCCTATTGTAAAATCATGCGCAAAATCATCTGATCGCTGTCTTGTTCCAGGAATGCTCTATG
GTCAAGTAGTGGTTTCGTTCCAGGCGATGTGCTTCGCACGCCGATTCCAATCACGCGCCTTCCGCACTCACTGGAATGCATTCTTTCTCTTGTTTCGAGTA
AGAAAGCTGATACTACCAAAGAATACGGGATGGTTCGATCCTTTGCGCTAAACACGCTATCCTGCTCATTTAAACTAGTAGGGTCGAGTTTATTGGATTTTCT
GCCTCGTCATGACATTGACCCAGATTTCATCGAAGAATGTGTATCTCATCAAAATTAACCACGCTTCACACCTTTTGCGGATCCAGAGTCGGCTTTCACTTTC
TGACAACATCGACAGAGGCGCGTGACTATAGTCGACTCGGTACTTGCCCACTAATACAATAAAACAAAAACCGAGAATGTATCATACGCATGTCCTATCCCTA
TAAGATACCAGAAAACAACAACATCAGTAAAATTTGCCAATTGCTAGACCAATGATGAATGTATGCATCATGCCTGTCGTCTGATTAAGACTGACATCTTGACT
AGGTCCGCGGCCGACATTGAGAATAACATGTTGACTTAACGAAAGTCAATAAAGATGACATTTGCTTGAGGCGCTTGCGAAATATTAGAACATGCTGTGTAA
ATTTGTCAAGAAAATCTTGATGCTAATTGCTAGACCAATGTTGAGTGTATTCATCACGGCTGTTATCTTGTCCAGACCGGCGGCCGAAAATTAGAATTGCATG

TTGACTTGGCAGCAGTCAGTAAGGATGACGTTTGCTTGAGGCGCTTGCGAAAAATTGGTACAGACTGTGTAAATTGAGTATATCATTTAACATTTTTGTGCAG
CCAAAAACCAACCAGAATGTGAGTTGATTTTTTCAGGACAAAGCATCGCGATCAAACTGAGCATCTTCTGTGAATAGATGCTCACTTAATTCAAGAACGCGT
ATTATGGCATAACTTTGTATGCCGTACACTTGATTCTGCCGTGACTGATGGGTGCGACTGTCAATTCAAGGGTGCGACTATATAGTTCAATGTGCGTTAAAGG
CTCTACTAGCTTATCATAAGCCTTCTTAAGCCCGTACAATTTTTTTCAATATTCAAGAAAATAAGAATTAAATTAATTTCGAAGTATATCGGACAATGGATCAAAA
GAAAAAATATTAGAAAAACAATAATGTTTTTTCTTTGAGGTTAAATCATTGAATTTTTGTGTAAGTAGCTCGTCTGTTCAACTAGTTATTAAGATTTATCTTCTCG
CAACATTTTAAAGTTTAAACAAATTCAAACTCTCCTAAAAAAATCGTTTTTACTTGATATCGTAAACGAAGGGGTAAATCGACCTGCAAAAGTACTCCTGATGA
ATAGGAAGGGGTTGCGAAAAGAGTGGTGAATCGGATCGGGAGAAGCCTCGACGAAACAATAGAACTTATGTACAAATCCGGAAATAAGCAGTCTTTTCAAA
GATAAGAATCTTAACAGAGACAAAGTGCATATATATGATTTTACCTGCACTTGAAATTATTTGGTTATTTTGACGAAAGGGTTATTTTTACCCCGGATGAACAGG
GTCTCAAATGTTTTGCTTTAATCGTCGAAAAAGTCGATTCGCTCTTTTCCAAACCTTGAAATATTTGAAGTCATGTGGCTTTTTCTTTATTGCGAGAAAAGCAC
TTGTCACTTTCGAAAGACTACGTCTCAATAGAATTAACGATTCATGTTTATTTGAGCTCGACAGCAGAATTATGTTTCACACCATGGCATGCAAGGTTGAATTA
TTTTCGAAACTCCTTTGCTAATAGGGGGATTCAGTCATTAATTCTTAAAATGCTTAGTTAACGTAGAAAATTTCTTGAACATGGTCAGATACCTTTCAGATGATC
ACCCTATGAGACAACGGATCACAAAAATGCATAACTGTTATTTGAATGGAGAATTCGGGGAAATAGGTCACCGGACATCTTTTTCAAACACAAAAACTGACAT
GTTTGCATTTACGTTGGACATTTCCTAGGGTTTACCTGACAGAAATAGACGATGTCTTACTACTTATTCTGTTTTGACTTCCGAAACGGCTCAGTTATACCTTT
TTGGTTTACTTTTAAAACAAGAGAATCTTTTTTGTGATTGCTTGTTTTAGGAT

>Centotomere_Chr14_h
AAAATCTAGCTTTGAATTTACGTTTCTATTAGTACTACGTATTATCATCATTGATGCTTGCTCTGTAATCCTCTTCCGGGGGTACAGAAATTGCAGAAGGCAAA
CTTACATAATAAGACGTACTATGTCATCAAACCGCAATAACACTACGGAGCGCTCACAAAAGCTTGAAGCCTAAGTGCGACTTATTAGTCAACCAAGTTTCAA
AGATAGAGCACCGTGCACCGTATCAGGTACTTGGTAACGGAGAGGAATGAGTTGATCAAGCCGGTCGTTTCTAGGTCAAGGAAGACACCCGGTAACAGGA
AATCAGTTGTTCGGTTGAATTAGCTGTTTGACAATTAAGAGCACAATAGCTATAATGGCTAGAATATTTGAATCGTCTAATTCGCTTAGACTGCTGTCAAAAAG
TCTAAAATATACTGGCTATGGTAATTCGTTTACCTAGCTTTCACGATCTCATCCAAGTGACCGTTATTCGTGACAAAGAACAGACAGCGGGACTGGATGGAGA
GCTTGTGGTGCTTTGCTGTCGTCCATAGCGAGTGGAGTATAAGTCGTTGGCAATGCTGTTGTTTCATTGCCTCGTATAAAGCATAAATAGCACAATCCACAG
CAAGTCAATCTTTGAACATAACTTCTAAAGAATGAAACGCTTTTATCGTTATCTCGTTGGCTCTAAAAGCCAACAGCCTGTTCAAATCAACGGAGAAGCCCAC
GAGCGGACAGTTTTACTCAACGCTGGCGAATCTTCGCAATCTACGCTTTCAAGAACGAGATGTTTACAAACAGATGAGGATAATATGAGGTCAGAAAAATGT
TCTATAAATAATGAAGCGAGCCACGCGTCTCTTGAAAGTTTTTTCTTTCCGGCTGACAACCCTTCGATACAACGATATTATCGATTTGAAGCTACCCCAATTAC
TCCGGTTGTCGCTCTTTATAAGCAACCCAATACAGACAGTGGAGTGACGGGTCTGCTGAGAAGATCAGCCGTTGTTCCTTCTCACGGGACATCGGATCAC
GATTGGATTTTAGTGTCGGTAGGAGGCCGGTCAGGGTGGGCTCGTCGTCATCCAGCGGGGCCTTTTCAGCCCGCTGATACATTTAAAGCTCATCAGGCTT
GGATGGGAAACCACGCTTTTTTATTCAAAGGAAAGTGTATGCTGGGATCGGACGCACCAAGTTTGGTTTTAACGAACGCTTTGTTGCTGGGTTGTGGGCTC
TATCACTTTCTCGTTGTTTTACCGAGACTCGCAGCAGTACACACGCAAACATCTATTTGGTATGGAGCGTCAATTTTTCTCTTACTTGCTTCCTTCATACTTTT
ATGGATCACAGCTTGTTTGGATCCAGGAATTATACCCGCCCGGTCTTCGCCCCTTCGACCTGAGCCTCCCGCCGGTGTAGTGGGTCATTCTTCAAACTCGT
ATCGCTACTGCTCAACATGTAACATATTCCGACCTCCCAGATCGAAGCATTGCAATTCCTGCAACGTGTGTGTTTCAGAATTTGACCATCACTGTCCTTGGA
CCGGAAACTGTATTGGCGAGCGAAATCACATGTGGTTCGTGTGCTTTTTAGTGGCTGTCACTACACTCTCATCGTTCGTTACCGGATGTGTAGCTATACTTG
TGGCAGATAATTTTTTCGAACTTTTGGAAATGAACACGGGAGGAATCGGGCCCGTCATCAAGTCAAAGGACAGCATGGATTCGACAGCCATACCGACATCA
TTACTTTACGACAAAGATGCGTGGTCTACTTTGTGGGAAGCCATTTCAGCTATGCCCGTTACCGTTTTGTTCGGGTTATTCCTGTTGCTATGCGCGTGGTCA
CTTTTGTCCTTACTGGCATACCATGCTCTCTTAATATCCGTTGCGCAAACAACCAATGAAAGGGTACGCGGTGTTTATTTAACATCAAGAAACCCCCGCGACA
AGGGCTGCTGCCTCAATTGTGTTACATTCTTTACGCGGAGAACGTCTCCAAGCAGACTGCCACCTGATTTTTCCAAAGAAGTAAAGCTGCAGCATGCTCGC
GTTGAGGCGCCATGGAGTGATAACTCGCCATAGCACTGCCTCAGTCACGGAGTTATTAAACAATTTAACTGTAACAACTCTGCTCTTGCTTTGAAAAGTCGT
AGGCGGTAAATGGAGAGCGAGATTGCCGGTATCTCTAGCTAGAGAGAGATAAAACTGTTCGCAAATGCGGAGAAACCTTTCTGAGAAAAGATAGATCACCT
ATGGAGATAACAGATAATCAAATGATCTTCTACGACCAGAAATACCAATTAACGTCAAAAAGACGAAAACGCAACCTCACCTTGTGCAATTCAAGGCTCTTCC
GCTCACGTCCTAAGCAATGCATTCCCCCACAAACAACAAAAATAAATAACCGGCGATTCACACCTTCCATATTTTTTTTAAAAATTAAGATACAGCAAATACTT
CTAAGTATTTATTCCTAATTTCCAAAAGTTCGAATGAAATTTTGACATTCGACTGAATTATCCACTGTTTTTCCTACTTTTTTCATATTAGAATTTTTATTTACTAGA
TTATTGAATTTTTGAGATTCGACTGAATCCTCTTTTACATGTGCAAAGATTTGTAAAGGCAAAAAAAAAAAAACTGGAAGTACTAGCTACAAAGTTTTCCCATTT
CTGCCCCGATGATAGCCGATGCACTCGACAAGCGCTTACAAGTGGAAAACGGTCAATGTAAAGCAACCATGTTAGAAATCGTAGTTACTATCCAAAGCAGCT
ACAATCGACGATAATTTCAAAAGCAGTTTCCATTTGAAACCATGTCATAAAGTAGAAAGTGGCACCGTAGTTAATCAATCAAGGCTAACATAACAAATCAAACG

TTTCGACAGGAATCATTAGCAGAGTTGCTTTACAATCTAATCTAACGGTATCGAAAAGATATTTGTAATGAAAGCGTCTTCTGGTATCAAGGCGCGTATCAACA
TTATCAAGTGATTATTTTTGCTATGATATGGGTTAAACACCCAAGGTACATGGTTGACAGTTTTTGGAAAGCAGAACTGCCTAAGTAGACCTATATACTGCTGT
TGTAGCAATCGTCCTTTAATATGTATTTCGTGTTGCTGGGCAAAAAAACACTTTTGTCAACCAGACGGCAAATTGTATCAACGATTAACGATCTATTATTCCTA
GAAGCCATTCGATTTGTTCTATCCACGAAAATTTTCTCATCTCACACGGTAGACAACCGTCTTGACAATAAAGTCACGAAATTACAATTCATATAAGCCGCTCG
CGGCCGGCTAGGGGAGGGGAAGAAGGCAATGTTGGAAACCAATCTCGACACGCTTTAAATCCCGATGTCACCTGGATAAAGGCGCCCAAAGATATGGCTT
CTCGGGAGAAACTTGATGCCGTGAAAAAAAGAAGAATCGTGCAAATATGTTGAAATATTTTCTATATCATTCGCTCATGGAGCCAGTATGATCCAGCCTTTTC
GGTCGCATGTTACATTCCTAACAAGAACGGTCTTGTGCCGCAGAGAGTCGGATCCAAGCTATTCAAATCAATCAGTTTTTCACCACGATCTGCAATTTCAAA
AGTTCCAACTGTTGCTTGCAAATTGGTTATAACGCTCAGATAATCAGTAGCATCTATGTATCTTAACCTTACAAGCATTATGGATGCGGATCAATACCCACATTT
TTGTCCCTAGGCTGTTCTATTCCCGTCAAAAGTGGACAAGAACAGTACAAATGAGGTACGGATCTGGATCGTTAACAAGTATTCGTGCTCCTTTTCTTACATA
TTTTGATTGATTTGTACGCAAAAACGACGAATCGGAAGTGTTAAAACACAGTCCGCGAGCTTCCAAAGCTTAATCAATTTCCTTAGAAACGCCAACCTAATTG
ATTTGGTCTTATTTTTGTTTTGCGTCTATCACGCATAGTCACGGCATGAAAATATTCTTATATTGTTTTAAATTTCTTGAATGTAGTGTTGTAAACCAATCACTGT
CTGTTAATCTCAAACCGGGGTCAAAAATCTCTCATTTTATGT

>Centotomere_Chr14_I

TTTCGTCTACTATTCTCATCATTATTACTTGCTTGTAATCTTCTTAAGGAGTACATCTTTTTATGAGTGCCTAGCGGGTCGCAAACTTATACAACAAGGCGTAGC
TTATCAAATCGCACTAGCAATGGGCAACGCTTGCAAAAGTTCAAACTCTATCTACGATTCATCAAGCAATTCAGTTGCATGGATAGTGCACCGTGTTAGGAAA
CGGTTAGAAGAATGAATTGATCAAGCCAGTGGTTTTAAGGTCGAAGAAGATACCTGGTAACAGGAAACCAATTGTTCGATTGGAGTAGCTGTTGGAAAATTA
AGAGGACAAGATAATGGAAGCCGAATTAAATTTGCTAGCTCGCTTAGACGGCTGTCGAAAGTCTAGATGAACTGGCTGTGGTAATTCGTTTACCTAAGTTTC
ACCATCTCATCCACGTGACCGGGATTGTTCACGAGGGACAGACAGCGGGACAGTTTGGAGCTTGGAGCTGGAGGCTATGGAGAGCAGGAGATCGTGTTC
TTGCTGTCGTCCGTCGAGGAGTAAGTTTTCATTCGTCAGCAATGATGTTGTTTCTTGTTTCGTTTTAAGCATAGATCGCATAATCCACCACTATTCAACCATCA
AGGCGTAGCTGTAAAAAATGAAACGCTTTTATCGTTATATCGTTGGATCTAAAAGCGAAAAGCCTGTTCAAAGCAACGGAGAAGCACACGAGCGGACAGTT
TTACTCGACCCCGGCGAATCTTCGCAATTTACGGTTTTATCAATATCAAGAACGAGATGTCTACCAACAGATGAGGTTAACGTGAAGTCCGGAAATTGTTCTA
TGGATAACGAAGCGAACCACGCGTCGCTTGAAAGCTTTTTTCTTCCCATCTGACAACCCTTCGATACAACGATATTACCGGTTTGACGCTACGCCAATTACT
CCGGTTGTCGCTCTTTACAAGCAACCCAACACAGACAGTGGAGTAACGGGTCTGCTGAGAAGATCAGCCATTGTTCCTTCTCACGGAACATCGGATCATG
ATTGGATTTTAGTGTCGGTAGGAGGCCGGTCAGGGTGGGCTCGTCGTCATCCAGCGGGGCCTTTTCAGCCCGCTGATACATTTTTAGCTCATCAGGCATG
GATGGGGAACCACGCGTTTTTATTCAACGGAAAGTGTATGCTGGGATCGGACGCACCAAGTTTGGTTCTTACGAACGCTTTGTTGCTGGGTTGTGGGCTC
TATCATTTCCTCGTTGTTTTACCGAGACTCACAGCACTACACTCGCAAACATCAATTTGGTACGGAGCGTCAATATCTCTTTTACTTGCTGTCTTTGTACTTTT
ATGGATCACAGCATGTTTAGATCCAGGAATTATTCCCGCTCGCTCCTCGCCCCTTCGACCTGAGCCTCCAGCGGGAATAGTGGGTCATTCTTCAAGCTCAT
ATCGCTACTGCTCAACATGTAACATATTCCGACCTCCCAGATCAAAGCATTGTAATTCCTGCAACGTGTGTGTTTCGGAATTTGACCATCACTGTCCTTGGAC
CGGAAACTGTATTGGCGAGCGAAATCACATGTGGTTCGTCTGCTTCTTAGTGGCTGTCACGACACTCTCATCGTTTGTTACCGGATGTGTAGCTATACTTGT
GGCAGATACTTTTTTCGAACTTTTGGAAATGAACACGGGGGGAATAGGGCCCGTAATCAAGTCAAAGGACAGCATGGATTCTACAGCCATTCCACCATCATT
ACTCTACGACAAGAATGCGTGGTCCACCTTATGGGAGTCCGTTTCAGCCATGCCTGTTACAGTTTTGTTCGGGCTTTTTCTTTTGCTATGTGCGTGGTCACT
TTTGTCCTTACTTGCATACCATGCTCTCTTAATATCAGTTGCGCAAACAACAAATGAAAGGGTACGCGGTGTATATTTAACGTCAAGAAACCCCCGTGACAAG
GGCTGCTGCCTCAATTGTGTCACATTCTTTACTAGAAGAACGTCCCCAAGTAGACTACCACCTGATTTTTCCAAAGAAGTAACGCTTCACCATGCTCGCGTT
GAAGAGCCATGGGGTGATCGCTCGCCCTAACACTGCCTAAATCATGGAGTCGTCCTCCATATTAGACGCTACTTTTACGATAACCGGCGCTGCTTCTTGTCT
TGGAAAAGTCCGAAAGGCGGTAGAACGGTGTGCGATTCTGCCAAGCAACCCGCAAACTCTTCGTCAATGTGACTCTCAAGGAGACATTTTCTTCGATAATG
TGACTCTCCAGGAGACATTTTCTGAAAATATAGATCGCTTACGAAGACTTCGGCAATCAATTTTAAACTGCTCCATTTGGTGACTAATTATTTTATTCCCTAATT
TCCTTGGCAGAATGCAATTTTGATATTCGGACTGATTTTTTCGTCGTCCATTTTCATTCTCACATAAAGTTCAAAGTGGCCGCGTGCTGTGTAACTCTCCAGC
TACTAGTGCTAGAGTAAAAGGCGAGTCCTAAACCAGTGTCCTAGTGAGAAGAGAATCTCCGTGGTGCTCCTTTTCTTGGATATTTTGACGAAGCATACTTAAT
TGTCCGGAACAGAGAAAGGCTCAGGCGAGCCCTAATAACCCCATAAAAATGTCTGACTAAGGTTTATCGCAAATGCTTGTTGTCCCATCGATCCGTCAATAA
ATATCTACAAACATATGAGAATGCAACCTAAAGTCAAAATTGCATTCTGACTTGAGGAAATTGGGAATAAAAAGGTTTAAGTGGATATTTCAAACATTTACAAAC
GAAACTGCCTCAACGAAAGCGAGTTCAAAAGAATCTTCAAATCAAGCCGATGCACAAAAAACAAAAAATGCGATATAAACGTAAAAACCACTTTAACACAAC
CAACTGCAATATATGCGAATGAAAAATGGAAAATTCAGTCGAATGTCAAGATTAGATTCTGACGATTAAAAAATTAGGAATAATATTAGTGGAACTGTAAATCAC
TTGTGAATTCTCAAATAACGCTTAAAGCACAGTAGTTCAACAGAACACAAAGCATAGGGTAAGCGATGACTCAAATCCTCATAAAAAAGCAGAAAGCGAAAC

GAATAAGGAACTTTTTAAGAAAATGAGGCTACTGGGCAAAAACCGAACAAAATCTTCATTAAAGTCGATAACAAAGGTCCATAATTGGTTGAATTCTTCGTGG
GGGTTCAGTTTCAACCTATTCGCCAGCACTTTGAAAATTGCAAGCCATCAACCCAACAACATAAACAGCAGCTGACTTTAAAGGGTAAAAAACAAAGCCCCC
AAAATTATTGTAATAATCACAATATGATGAAACCTTCGTTTACTTTTCGCGGGTACACTATGTCTGTTTGAGGACACGTATTCAAATGATTGTTGTGAAAAATTG
GTTGTTGAATCATATGTATAATCGTTCAACTTTGATTTATTTCCTCTTTCATAAAATATACACATTGAAG

>Centotomere_Chr20_h
AACAGTTTGAATGAGGCAAAATTCGGTTTTCTTGTAGGGACATAAGCATAGTAAAAGTAGTAAATTCGTGCCACTTCAATACTAAAGCCATAAGCAGAAGATG
CAACCCCGCCATATTTATGCGATGAGCTTTATACTTTATAGACCAAAAGAACAAAATATCTTACTTAATCGCGATGGAATTGTCTGTCTAAATTACACGAGATTT
CTTTGTCAGAAGCAATGGGACCGATGTCTGATTTCTTCAACAGAGGTTAGACATTCAGATCCTTACGCCATAAAAAAAAGACTTTGCGAGTTTATAAAGTGG
GTAAACTATAGATGCAATCGTGTGCCATGCTGTAGAATCATCAATCCTTTAAAAACTTATGTTTGAGATAAGCTGCCCGAAGGCAAGGCAACATTAATTATAAG
TGTTAATCAGTTCGTAAATCATGCGAAGATGAAAATTCTTTTGTGTGGCCTGTTCCCTCTAGCCCTCCTTGCAAACAGTCAATGTGCGGGTAGACCAGCGAA
AGTCCTCTTTGTCTTGTTGTCGATAAAACTTCGCCGGAATTATTGCTGTAGCCCCTAGATAGAATACTTTAGCACCGTAGTTACAAATATGGTTGGCAGTTGT
CTTTATGGAGAAATGAATTTCATTGGTGGCGAATAATGCCTTCATAGCAAGGCTTGACCCCCGCGTATCTTCTCAATTCATACCATGTGCTTTATGCGAAAAA
ACAATTACTTACTTATTGTAAAAGCTTGTGTTCTCTTCGATCTCAATCCGCGTCTATAACATTATTATTATTATTGTTATTCACAATGATCTGACGTGTGAGAACC
GAAAGTGTTCCCTACGTTGAACCAACTCGAGGTTGATAGAGTGTGTTGTGCATCATAGATGCGTCCAAGTTTCCAACGTGGTAGTATGTCTAAAGGTCATGT
TCCTCCTCATCGAACCAGTCCTGTTCGTAGGTTCCCTTGCAATCAGAATGGTGTGAACCCATGCTCTGCAATCTGGCATGGCACTCTCGATGACCGCTTGG
ATTGGTTGGTGTAGCAGGTCTTTGTCTTTAGGCAGTAATGTGTCCTCCGTGAGGTCGTACTGGCGGTGTATTTCGGTAAATAATGCTTCTACTTCAGATAATT
TGCGTCCGGCATCGTCGCGTAGGTGTAAGATGGCGTTTCGGCAAAACCACATGCTTCTCGTAACTGCCGTGAGTGCAAGGCTGGCTGTGGCTGCCTAGG
TCTTTGCCGTTCTTCGTGATTCATTAGCCAGATAATACAACGCGATAACGATACAGAACGAGATCGACGTCCACAACACCCTACTGGGCCGCCTAGCGGAA
GGCTGGGACTATAACAATATTTCGTTATAGACACAAGCTTGCTTGACGCCTCCATATAAGTCGAAATCATCCAATAATACAAAACAAAACTATTACGATATTGAG
GACACGCACAGTTATCGTATTTTGTTTTTAAATGTTCGGTTACTTTTAACATTGCTTTGCAGTTTGAGTGATGCCCAATCTACGATGGTTTGAATATCTATTCTT
GGCTTCAATTATTCACTCGAAAGCCCACCGCTTGGGATTTTACAGGTGGTTCGATACGATTTGCTAGCTGGGAAAAAGGAAAAGATGGCAAATGTAAAATCG
AAAACTATTAATTGTTAAAAAATAGAATGTCTTTATGGAGAACTGGAATTCATTCGTGGTGAATCAAGCTCTGAAAGCAAGGCTTGACTCCACCATATATTCTC
AGTTCCCACCATGTGCAATTGGCGAAAAGCATGTTCGGGGTATGCTTTGCCCGCCTATCACCCAACCATTTCCTAACTTATTGTATAAGCGTATGTTATCTTC
GATCTCAATCCGCGTCTACAAAATATTTTGTTATAAACATAAGCTGGCTTAGCGTTTCCATATTAGACGAAATCATCAAATCATACGAAATAAGAACTCATCATTT
TTTTGGTTTTTTAATGTTCGGGTGTTTTCGCGTCTATCAAAATACTTCGTTATAGACAAAAGTCCAAATCATCCAAACATCTATACATTGA

>Centotomere_Chr20_l
ATAGAGAACTATACTACCCATGTGTGGTGTGGTGCATGTTGGGTGCATCACATTACCGGCGCCGTCTACGTCTGACACGGCGTTGTAGGAATGTTCTTATGT
CTGTTTGTCTCAGGCTGTTTCGTTGCTTCCATCTGGTGTGACTCCGATGGATATTGTCCCTATTTTGTTCGATATCCACGTTTGTTTTCATCCCTCGTCCAAT
GGTTTAGCGAATGGGTATCGGATTGTGCAATGCGGAATGAGAGCATAGAGATTTTGGGCTGTATGTTGTGCTTGTCTTCGAGCCGTTCGTTTCGCGGCAGT
TTCCGTGTGTTCATGGTGAAGCGTCGTGCGACTTGTCCAAGCTAATAGTGTCTGTTGGATGATAAAGTGCATGAGATTCTTCGTAAACCAAGTCCGATCAAG
TACTTCGTGTTCAGCGTAATGCTTATCGATCATGTTTTCCCACTATGTGGGGATAATGCCGCAAGGGATCAGTCGGACTGGTATTTATGAAAGGGGTCGTGC
ATCTGGGTAGTGCCTTATATAGAGACAGTGTGATATATCCCGTGCAACCCTACATCACGGAAGCAATAGGCTGTAGACCGAAAAGCAAATAAAATAATCATTAT
CATGCGATTAATCCCCGTCGGGAACAATCGTATGGTACTCTCATCTAATTTATTTGTCGAGCGGGTGATCCTTAAAAAAGGATAATGATCGGTAAAACTTTTAG
CTTCTGCTTGCACGATTAACTTTCGAAAGACAATTTTATGTGTCAATGTAAGAATTGAATCGTTATACATCTGCTCATTGGTCCATATCTTAAAACAAAGAAAAT
GATTGTCAACGGTGGACTTTCTGCCATGAAAAATATAAAGAATAAGTCTTCATTTGACCAGATCATTGACTGATACGCTTTGTTTCATGCAACCAATTGTCTTT
TGCCTTTCTTCCATTAGACGATCATCGCAAATATAGCATAAATTTAAATTAAATTTGCATTGTTCTCCTGTTGTAATTGGCAGAGAGATCTAGGTTGGGGTTAG
GTTTCATCAATTTTGTGATTAAGGAAAGCGTCCGCCCAAGTTTACCTTTGATTCATTGTGTTCGCCTCGACACTGACCGTTAATTTTACTATTTTTTACTAAAAA
GACTACCAGGCCAAAAATACCATCTGCTCCAAAGTCTACCGTTAAGAAATAAAACACGACTCCTAAATAGGCATATTATGCTAATTGATTCCAAGCATCGTTCA
AAGCTGATCTAGCGTCCAAACTGAGTTTCCTACTCAACGCTTAGATAACTATTTCTTTCAGAGAACGTCAATGTTGTTACAAAATAAGCTATCCGAATGCTAGT
TAATTATTTTCTGTCATCCAAGCTAGTTAAGTGGTTGAGAATTCAAGAGAAAAATGGCGCTCCTCGGAAGTAGATCGACAATCGACAAAAGTATCATATGATTT
GTTTTACCAAGAAGGAAGAATGATTAAACATGGCAAATTTAATTGTAGAGAGAAAGAGCGTTCGAGATAAGTTATTGTATTCTACTTATTAAAAAATTGCAAACT
TTCGTATGTATAATTTAATCTTTTGCTGTATTTCCGCAGGACAGTTGCATCTCTTTGCTTTGAGCCAAGGACTTGGACGAACGCTTTTTCTAAAATCATGCAAA

ATTCCAAAGCACAGAGCCTTTAAAAGGAACATACTCAAAACGCTAACGGGCCTGGAGGGCATATTTCGGTTTAAGACGTTTAAACATCAGATTTTAAATGCA
GGTTGTAGAACCAGTATGCTTAAATAAAAGTGAATAGTTTTAGGAAGCAATCTTAATGACGCATAAATATAAGCGAATTTGAAACTTGATGGCGGATTAGACTT
TATTTAGACGTCCATTATGGCTAATTTGAGTGTTATTTAAAAAGCCAC

>Centotomere_Chr22_h
AACATAAATTTTGACATAAGTCTCCAACTCAGCGTGAAAAAGTGATGGGGTCTTAGAAGATTGTCTCCTACCCGCACCCTAGACTTAGCATTTGGGTCTTCTA
CCAGCAGTCTCTGCGCCTATGATAATGCTATGCATACAAAGTAACTTTAAACAGTTCAGTGAATTTTCCATCGTTTCATTTTCAATTTCCACATACTATTAATCC
AATACCGATCTGGCACTTTGCCATTGGCAATCCTTCTCTTGGATTTTAACAAAGCATTCTAACCGACCAGACCAGAGAAAGGCTCCCCTCATGCAAGCTCAT
TGTACAAATTTCAGAGTAAGGTGTATTCCATTACCGTTGAAGTTTCTTTGCCAGTTGTCTCGATGCTGGAACAGCTTGCTGTTTACATTGTTTACACGGTACG
GATATTCACAGCGAATAACGTCCGCCAGCCAATCAACGGTAAGATCTTACCATCAGTCATAGAATTCAAGAATTCTACTTGAATTTAATCAAGTAAAAGGCTTT
ACCAAATGAATCTGAAAACTGATCAGCTTAGAGTGGCAGCAGGAGAACTTGTTTTTGATGCCGATTTCTCGGGGATGGAATGAATCTGTTTTGCGCAACGC
TCTCGTAACTTTTCCAATTGAGGTGAGATACTTTTCTTCAAAATGACATTGCAAGTTTCGATAGTAGAAATCATTGCAGACAATCTCAAAACACCAAATCAAAT
GAATACCAAACATTTACCAACAAGTTTTTTTGGTGTTATTGTGACATTACTGGGTAATGAACACTTACGACAATGAGAGACAACAGTCGGTCGTTAATGCTCG
CTATTAAGATTCGATAGCGGATGAAGTTGAATATAATGTCAAACCATAAAATCGCAAGCAACATTGAAACCAAGAACTAAATATGCACTTCAGTTTTTGAGTTT
CGGAATCGGAGTGATTGTATACGATCTAGAATCACTATATCGCACGTAGTCCGAACAATGGCATTTCAAGTTTCGATTGTGGAAATCATTTCAAACGATCTCA
AACCTCCAAATCACATGAATTCCATATATTTACCACCAAGTTTCTTTAGTTTAATTGTGACATCCTTAGGTAATGAACACGAACGACAATGTGATACAACAATCG
GTCGTTAATGCTCGATCTCAAGATTCGATAGTGGATAATGTTGAAAATGGTCTTAAACCAAAGAATCGCAAGCAATATTGAAACCAAGAACTTATGTTTGTGTG
AGTGTTCAATGCATCATACGCACTTCAAATTTCGAGTTTCGAAATCGAAATGATTGCATACGATCTAAAACCATTATATAGTGTTTATGAAATCAGTTGGTAAAT
GATTTTAAAAATGACCCGCAGTACTTGCTTCTAAAATTTCGATAGTGGGAATTATTGCAAAGGATCTCCAGGCGCCTAATCGCATGATATCCGTACAATAGCCA
GTTCTCTCAAATCAATACATCATTTGCACTCTCTTACAATTGGCCGTAAATAAACGCTATGCTATCAGGTACATGCTTATTTGTAGCTTTGTATAGCCGCTTGAG
GGTAGAGATTGGGCGTCATTTTCGATATGAAGAAATGTGGAGGACATAGTTTCGAAATCATTGGATACGATCTCAAACACCTGTATCGCATGTCAGCTGTACA
ATAACTAATGTTTCACCATTAATGTTCACTTTAGCATAAAGATGTTACGCAGTGCAATTTGGTTTTTATAGCTTAACGTAGCCAGGAAATAACATCGATTGTTTG
TAATCATTTGATGCTCGCGACTAAGGGTCATAGTGTTTTAAGGAAAAAGGTTCTAACAAACTTTCAACATTTGAGCTGTACAGGGTCTAAATACACTTACTACA
TAAAGATCTTACGATTCACCATCTTAGGTCGCTTGATCATATTGATCGTGTTTTTGTTACTTAATGATTTGACCAAAGCTCAAAA

>Centotomere_Chr22_I
AACATAATTTTTGATAGATCACAAGGGGCGGCAGCGCTAATGTAAAGGGTCCCCCTCCTGGCAGCACATAGGCCATCTCAAGAAAACGTTCTCCTACCCGC
ACCCTAAACTAAGCATTTTGGGTCTTCAACCAACATTCTGCAATAAATTTTAGGCCTATGTGGTTGCTTTTTATTCTTTTCAAATATAATCCATTTCCCATCGGG
CACTTTGTTGATAAGATTCAAGAGCTCTTGAGCTCTCGTCTATAAGAGAGGAGGTCAAAAGCCCGTATGACAAAGAGAAAGGAATCGCCGTGGAATTTGCT
TCTCTTGGATGCCGACGAAGCATACTTACCGACAGAACGAGAAAAAACTCGTGCGTTCAAATGAGCAACAAATCCTTGACTTTGGCTTGTTCCATTGGCGC
TGAAGTTTCTTTCCTAGGTGTCTCAATCCTGAAACAGGTTCAAGTTAAAGTGGATCACGGTTAATTTACAACAAAGTAAGCACAAAATCCCACCACGAGCAA
AGGAAGTCAAGAAGCTTTACTTGAATATCAATAAGTGCACTGCCGAAAAAAGAAGAATGTAGTATTAAGTACAACGTAGTGCCTTAACTGTTGCCGACAGCA
AAATCAAACGTCAAATGTTCATCTCGCAAAACTTAATGCTCAATAAAACGCTCCAAGGCTTACGGATAAAATTCTCTGTGAAAGATCGATGTCAAGCCACCGC
ATCGCAATTGAAGTGGCAAAAATGTTAACGGAATAGTTCATTGCCTGCATTTTTGTACTACAATATCGTTTCTGGATATTTAAATCAATTTTTTTATATGCAAATT
TTCACGACAAAAACACATATATCGTTGGATGAAAAAGTTTATAGGTTTTGAATATTCAAATGTTAACAAGTGTTGATTTATATTCAATTTGTGTTTGTTTAATTACA
TGATTTAAATTCAATTTGTGTTTCTTTTATTATTACTTTTACTGAAACGTTAAAAAATAACTAAATAATCGTTTAAATTGTCGAACAAAATTGAAATCAGCATTTTAC
TACAGATGTCTATTGAGGTCGTCCTATTGCGACAGACGTAAAATAGGATAGCAGTGTCTTCAAGAAGATCACAAATTCTTTCAAACAAATGCTTTGACCAGAC
TGTGCCAACAATTATATCATTTTCTTGTACTTTGCTGATAATTATAAAGGTTACTATTGTCTGGATGCGCTAATGCTACCCAAGTCTTCTTATCTGTGTAAATACG
GATAGCATTCTTTTACTCCTTTATTAAGCAAAATAGTGTAAGAGCCACTCTGCGTAAAGATTTCTTGAAGTTATCTTGTCTTTTATTGTCCTCAAAATTGTGGAT
TGCTTCATTTTCCTTACCAAAGGAATCTAGTTTAACAATTGGAATT

### References for Supplementary Information

Osada K, Maeda Y, Yoshino T, Nojima D, Bowler C, Tanaka T. 2017. Enhanced NADPH production in the pentose phosphate pathway accelerates lipid accumulation in the oleaginous diatom *Fistulifera solaris*. *Algal Res* **23**: 126-134.

Tanaka T, Maeda Y, Veluchamy A, Tanaka M, Abida H, Marechal E, Bowler C, Muto M, Sunaga Y, Tanaka M et al. 2015. Oil accumulation by the oleaginous diatom *Fistulifera solaris* as revealed by the genome and transcriptome. *Plant Cell* **27**: 162-176.

Tillich M, Lehwark P, Pellizzer T, Ulbricht-Jones ES, Fischer A, Bock R, Greiner S. 2017. GeSeq - versatile and accurate annotation of organelle genomes. *Nucleic Acids Res* **45**: W6-W11.
